# The ketamine metabolite (*2R,6R*)-hydroxynorketamine rescues hippocampal mRNA translation, synaptic plasticity and memory in mouse models of Alzheimer’s disease

**DOI:** 10.1101/2023.08.03.551808

**Authors:** Felipe C. Ribeiro, Danielle Cozachenco, Elentina K. Argyrousi, Agnieszka Staniszewski, Shane Wiebe, Joao D. Calixtro, Rubens Soares-Neto, Aycheh Al-Chami, Fatema El Sayegh, Sara Bermudez, Emily Arsenault, Marcelo Cossenza, Jean-Claude Lacaille, Karim Nader, Hongyu Sun, Fernanda G. De Felice, Mychael V. Lourenco, Ottavio Arancio, Argel Aguilar-Valles, Nahum Sonenberg, Sergio T. Ferreira

**Author notes:** These authors contributed equally to this work (Felipe C. Ribeiro, Danielle Cozachenco). These authors contributed equally to this work (Elentina K. Argyrousi, Agnieszka Staniszewski, Shane Wiebe). Corresponding authors: Sergio T. Ferreira, Nahum Sonenberg and Argel Aguilar-Valles.

## Abstract

Impaired synaptic plasticity and progressive memory deficits are major hallmarks of Alzheimer’s disease (AD). Hippocampal mRNA translation, required for memory consolidation, is defective in AD. Here, we show that systemic treatment with (*2R,6R*)- hydroxynorketamine (HNK), an active metabolite of the antidepressant ketamine, prevented deficits in hippocampal mRNA translation, long-term potentiation (LTP) and memory induced by AD-linked amyloid-β oligomers (AβOs) in mice. HNK activated hippocampal extracellular signal-regulated kinase 1/2 (ERK1/2), mechanistic target of rapamycin (mTOR), and p70S6 kinase 1 (S6K1)/ribosomal protein S6 (S6), which promote protein synthesis and synaptic plasticity. Stimulation of S6 phosphorylation by HNK was mTORC1-dependent, while rescue of hippocampal LTP and memory in HNK-treated AβO-infused mice was ERK1/2-dependent and, partially, mTORC1- dependent. Remarkably, treatment with HNK corrected LTP and memory deficits in aged APP/PS1 mice. Transcriptomic analysis further showed that HNK rescued signaling pathways that are aberrant in APP/PS1 mice, including inflammatory and hormonal responses, and programmed cell death. Taken together, our findings demonstrate that HNK induces signaling and transcriptional responses that correct deficits in hippocampal synaptic plasticity and memory in AD mouse models. These results raise the prospect that HNK could serve as a therapeutic to prevent or reverse memory decline in AD.

## INTRODUCTION

Alzheimer’s disease (AD) is the most prevalent form of dementia, affecting over 55 million people worldwide [1]. AD patients progressively develop memory loss and cognitive impairments, mood alterations, and disorientation. At the molecular and cellular level, cognitive decline in AD is caused by structural and functional deterioration of synapses [2].

Hippocampal mRNA translation (i.e., protein synthesis) is crucial for synaptic plasticity and memory consolidation [3], and becomes defective in AD [4, 5]. Control of both initiation and elongation steps of protein synthesis is deregulated in the brains of AD patients and mouse models [6, 7]. Developing drugs capable of rescuing brain protein synthesis, synapse function and memory remains a major challenge.

Effective pharmacological approaches to stimulate brain protein synthesis remain elusive. One potential candidate that recently emerged is the ketamine metabolite, (*2R,6R*)-hydroxynorketamine (HNK). Ketamine is an FDA-approved drug [8] that, at subanesthetic doses, triggers fast-acting and prolonged antidepressant responses linked to stimulation of protein synthesis [9–11]. However, potential side effects, including addictive/dissociative actions, may limit its widespread clinical application [12]. HNK elicits similar antidepressant responses [13] in the absence of significant detrimental side effects in mice [14, 15]. As a result, HNK has gained significant attention in the field of major depression, and it is currently in clinical trials in treatment-resistant depression (NCT04711005).

Even though complete mechanistic details remain to be determined, HNK activates mRNA translation and the mechanistic target of rapamycin (mTOR) signaling [13, 14, 16]. Here, we hypothesized that HNK could rescue deficits in hippocampal protein synthesis, synaptic plasticity, and memory in AD mouse models. Our findings suggest that treatment with HNK may prevent cognitive deterioration and restore memory in AD.

## RESULTS

### HNK prevents deficits in hippocampal protein synthesis, synaptic plasticity and memory induced by A**β**Os in mice

We first investigated whether HNK could protect against impairment in hippocampal protein synthesis induced by amyloid-β oligomers (AβOs), soluble neurotoxins that accumulate in the AD brain and trigger synapse failure and cognitive impairment [17, 18]. To this end, we treated acute hippocampal slices from WT mice with HNK (20 µM) for 20 min prior to exposure to AβOs (1 µM; 2 h), and assessed global protein synthesis using SUnSET. In line with our previous report [19], AβOs reduced protein synthesis by ∼ 20% **(Fig. 1A, B)**. HNK alone had no effect on hippocampal protein synthesis, but pre-treatment with HNK blocked the inhibition of protein synthesis induced by AβOs (**Fig. 1A, B**).

**Figure 1.**
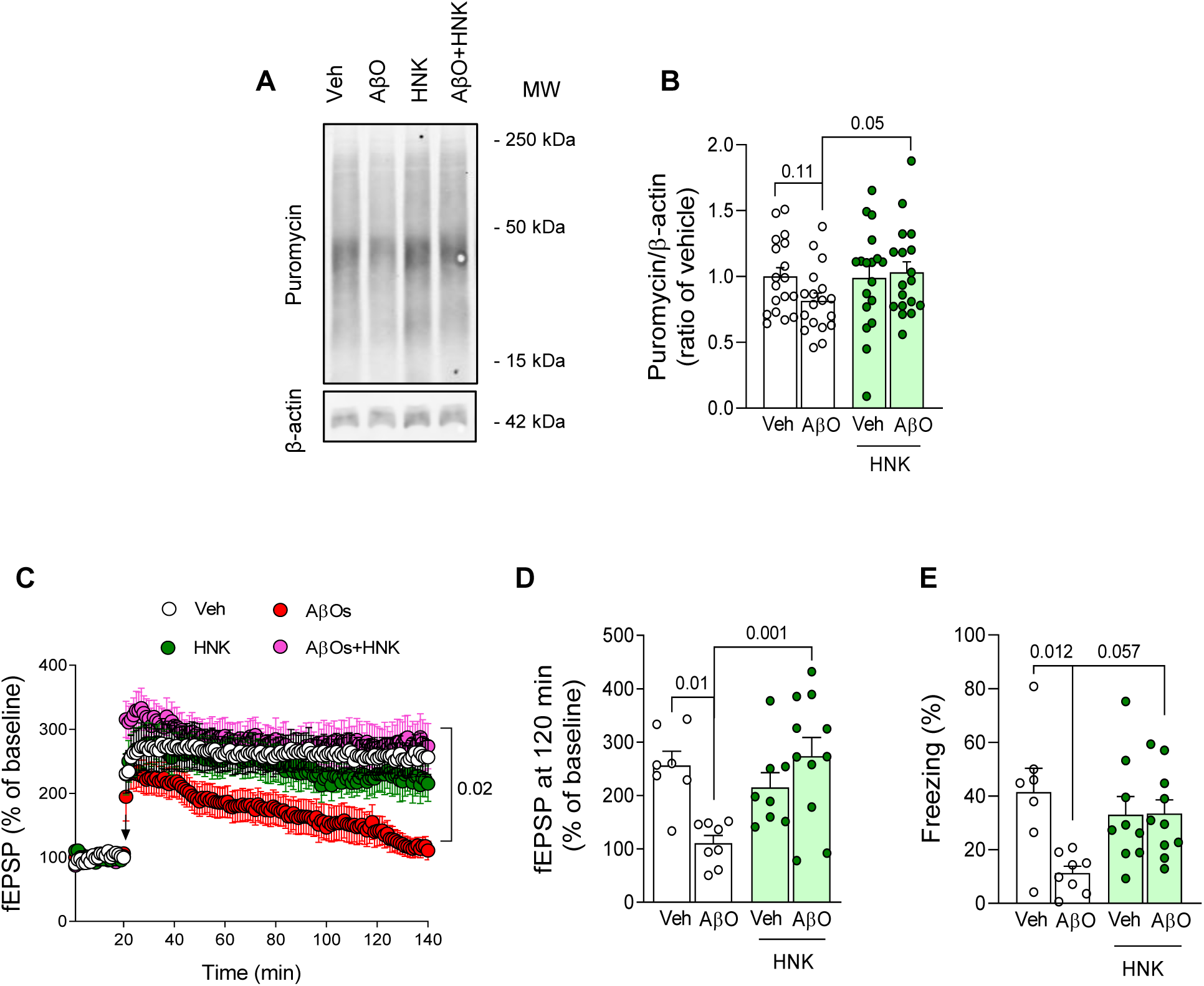
HNK rescues defective hippocampal protein synthesis, LTP and memory in AβO-infused mice. (A, B) Acute hippocampal slices were treated with HNK (20 μM; 20 min) prior to AβOs (1 μM; 2 h), and were then exposed to puromycin (5 μg/mL; 45 min) for SUnSET. Puromycin levels were evaluated by Western blotting and normalized by β-actin levels (N = 18 slices per experimental group). Two-way repeated measures ANOVA followed by Tukey’s post-hoc test. Interaction: F (1, 17) = 4.406; p = 0.0511. (C) Field excitatory post-synaptic potentials (fEPSP) recorded in acute hippocampal slices (from C57BL/6J mice) exposed to saline or HNK (20 μM; 20 min) prior exposure to vehicle or AβOs (400 nM; 20 min) (N = 7 Veh, 8 AβOs, 9 Veh/HNK, 11 AβOs/HNK). Arrows indicate tetanic stimulation. (D) fEPSP slope measured 120 min after tetanic stimulation. Two-way ANOVA followed by Tukey’s post-hoc test. Interaction: F (1, 30) = 11.68; p = 0.0018. (E) Six days after i.c.v. infusion of AβOs (10 pmol) or vehicle, mice were treated with saline or HNK (20 mg/kg, i.p.) 1 h prior to contextual fear conditioning (CFC) training. The CFC test phase was performed 24 h later (N = 7 Veh, 8 AβOs, 9 Veh/HNK, 10 AβOs/HNK). Two-way ANOVA followed by Tukey’s post-hoc test. Interaction: F (1, 30) = 6.311; p = 0.0176.

Next, to determine whether HNK could counteract AβO-induced inhibition of synaptic plasticity, we treated hippocampal slices with HNK for 20 min prior to exposure to AβOs and assessed long-term potentiation (LTP) at Schäffer collateral-CA1 synapses. Whereas slices exposed to AβOs failed to maintain LTP following tetanic stimulation, slices treated with HNK prior to stimulation showed normal induction and maintenance of LTP even in the presence of AβOs (**Fig. 1C, D**). HNK alone had no significant effect on LTP (**Fig. 1C, D**).

We further investigated the effect of HNK on AβO-induced memory impairment. We treated AβO-infused mice with saline or HNK systemically (20 mg/kg, i.p.) 1 h prior to training in the contextual fear conditioning (CFC) memory task. Basal freezing responses were not altered by AβOs or HNK (**Fig. S1**). While treatment with HNK *per se* had no impact on memory, it significantly prevented long-term fear memory deficits engendered by AβOs (**Fig. 1E**).

### HNK activates ERK/RSK and promotes mTOR-dependent S6 phosphorylation

HNK activates mTOR complex 1 (mTORC1) signaling, BDNF release, and increase synaptic function [13–15]. However, the mechanisms involved in these actions are largely unknown. To gain insight into the mechanisms of action of HNK in the brain, we examined the impact of HNK on signaling pathways implicated in synaptic plasticity and memory in acute hippocampal slices. We assessed the activation of mTOR, extracellular signal-regulated kinase 1/2 (ERK1/2), p70S6 kinase 1 (S6K1), p90S6 kinase (RSK) and ribosomal protein S6 (S6), all of which have been implicated in translational control, synaptic plasticity and memory (**Fig. 2A**) [20–22]. Similar to previous findings with ketamine [23, 24], we found that HNK triggered a rapid (5 min) and transient phosphorylation (activation) of mTOR at Ser2448 (**Fig. 2B, Fig. S2A**), and a sustained increase (10-60 min) in ERK1/2 phosphorylation (**Fig. 2C**). This was accompanied by an approximately 50% increase in phosphorylation of S6K1 at Thr389 (**Fig. 2D**), and by a marked (250%) increase in phosphorylation of RSK at Ser380 (**Fig. 2E**) 10 min after HNK treatment.

**Figure 2.**
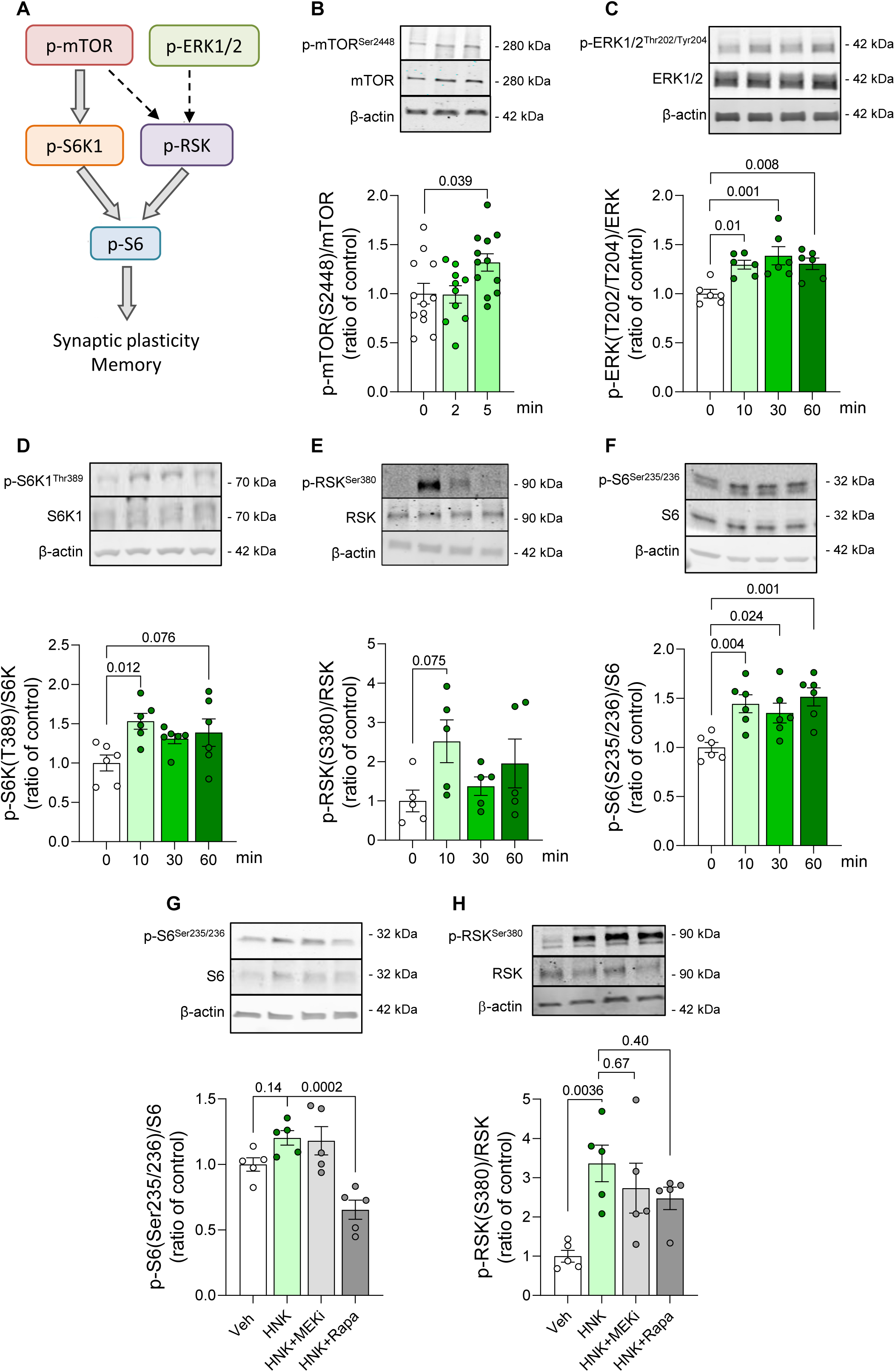
HNK activates hippocampal signaling pathways associated with translational control and protein synthesis. (A) Schematic diagram of signaling pathways that mediate synaptic plasticity and memory processes. (B-F) Acute hippocampal slices were treated with vehicle or HNK (20 μM) for the indicated time periods. Phosphorylated mTOR, ERK S6K1, RSK, S6 were evaluated by Western blotting and normalized by respective total protein levels (N = 6-12 slices/experimental group). One-way ANOVA followed by Dunnett’s post-hoc test. (G, H) Acute hippocampal slices were treated with vehicle, rapamycin (Rapa; 100 μM) or PD98059 (MEKi; 20 μM) for 30 min and were then exposed to vehicle or HNK (20 μM) for 10 min. Phosphorylated S6 and RSK were evaluated by Western blotting (N= 5 slices/experimental group). p-values are shown for the indicated comparisons; One-way ANOVA followed by Sidak’s post-hoc test.

Because S6 is a direct target of both S6K1 and RSK, we determined the phosphorylation status of S6 upon HNK treatment. HNK engendered a sustained (10-60 min) phosphorylation of S6 at Ser235/236 (**Fig. 2F**), residues targeted by both S6K1 and RSK [25], but not at Ser240/244, which are phosphorylated only by S6K (**Fig. S2B**).

We next investigated whether ERK1/2 or mTOR mediated HNK-induced activation of S6 at Ser235/236. We treated hippocampal slices with HNK in the presence of PD98059 (MEKi), a MEK1/2 inhibitor that blocks ERK1/2 activation, or rapamycin (Rapa), an mTORC1 inhibitor. Blocking mTORC1 activity, but not ERK1/2, blocked the increase in S6 phosphorylation at Ser235/236 by HNK (**Fig. 2G**). We further found that neither MEKi nor rapamycin alone blocked HNK-induced Ser380 phosphorylation of RSK (**Fig. 2H**), suggesting that both ERK1/2 and mTOR may be involved in RSK activation by HNK. As expected, MEKi attenuated HNK-stimulated ERK1/2 phosphorylation, but caused no changes in HNK-induced mTOR activation (**Fig. S2C, D**). Conversely, rapamycin robustly blocked mTOR, but not ERK phosphorylation (**Fig. S2C, D**). Collectively, these results suggest that HNK engages ERK1/2 and mTOR to phosphorylate RSK, and mTOR to phosphorylate S6.

### Correction of hippocampal synaptic plasticity and memory by HNK is mediated by ERK signaling

We next investigated whether HNK-induced restoration of synaptic plasticity and memory upon exposure to AβOs was mediated by ERK1/2 or mTOR signaling. We first treated acute hippocampal slices from WT mice with MEKi (20 µM), Rapa (100 μM) or vehicle for 20 min, then added HNK (20 μM) or vehicle for 20 min prior to exposure to AβOs (1 µM), and assessed LTP at Shaffer collateral-CA1 synapses. Consistent with our findings above, AβOs inhibited CA3-CA1 LTP, which was prevented by pre-treatment with HNK (**Fig. 3A, B**). Inhibition of MEK1/2 abolished HNK-mediated rescue of LTP, whereas mTOR inhibition did not (**Fig. 3A, B**).

**Figure 3.**
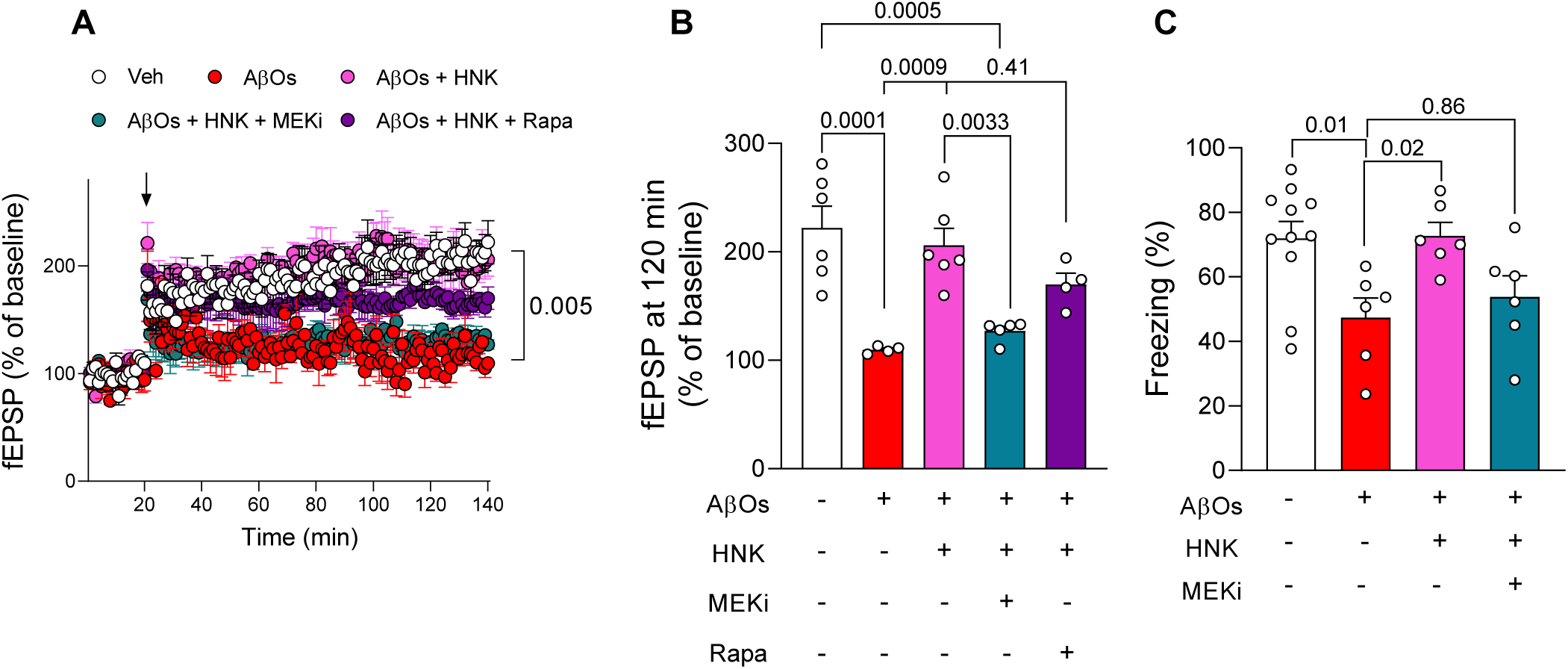
Correction of hippocampal synaptic plasticity and memory by treatment with HNK is mediated by ERK signaling. (A) Acute hippocampal slices were treated with PD98059 (MEKi; 20 μM), rapamycin (Rapa; 100 μM) or vehicle for 20 min, then HNK (20 μM) or vehicle was added 20 min prior to AβOs (1 μM). Field excitatory post-synaptic potentials (fEPSP) were recorded in hippocampal slices (N = 6 Veh, 4 AβOs, 6 AβOs+HNK, 5 AβOs+HNK+MEKi, 4 6 AβOs+HNK+ Rapa). Arrows indicate tetanic stimulation. (B) fEPSP slope measured 120 min after tetanic stimulation. One-way ANOVA followed by Sidak’s post-hoc test. (C) Six days after i.c.v. infusion of AβOs (10 pmol) or vehicle, mice were treated with MEKi (10 mg/kg, i.p.) or saline for 1 h, and then treated with HNK (20 mg/kg, i.p.) for 1 h prior to contextual fear conditioning (CFC) training. The CFC test phase was performed 24 h later (N = 11 Veh, 6 AβOs, 6 AβOs+HNK, 6 AβOs+HNK+MEKi). One-way ANOVA followed by Sidak’s post-hoc test. F (3, 25) = 4.752; p = 0.0093.

We then assessed whether ERK1/2 signaling was necessary for memory rescue by HNK. We treated AβO-infused mice with MEKi (10 mg/kg, i.p.) or vehicle, followed by treatment with HNK (20 mg/kg, i.p.) or vehicle 1 h prior to training in the contextual fear conditioning paradigm. While treatment with HNK rescued memory in AβO-infused mice, pre-treatment with MEKi abolished this protective action (**Fig. 3C**). Collectively, these data indicate that HNK-induced rescue of LTP and memory is mediated by ERK1/2 signaling.

### Transcriptional alterations induced by HNK in the mouse hippocampus

To further investigate mechanisms underlying the protective actions of HNK, we examined global transcriptional changes induced by HNK treatment in the mouse hippocampus. For this, we treated mice with HNK (0.5 mg/kg, i.p.) daily for 14 days, and assessed hippocampal gene expression by RNAseq (**Table S1**). Among differentially expressed genes, HNK promoted the upregulation of 469 genes and downregulation of 433 genes (**Fig. 4A**). Consistent with previous findings with ketamine [26], gene ontology (GO) analyses revealed that HNK induced the expression of genes associated with mitochondrial metabolism and transcription (**Fig. 4B**). At the same time, it downregulated genes linked to GTPase activity (**Fig. 4B**). Network analysis indicated that HNK upregulated several mitochondrial genes involved in the electron transport chain (e.g., mt-Cytb, mt-Co1, mt-Co2, mt-Nd4), RNA metabolism and ribogenesis (e.g., Wdr42, Ddx10, Ddx18, Esf1). In contrast, genes such as Src and eIF6 were downregulated (**Fig. 4C, D**). Altogether, these results suggest that HNK promotes a transcriptional reprogramming that modulates mitochondrial energy metabolism, transcription, and ribosomal biogenesis in the hippocampus.

**Figure 4.**
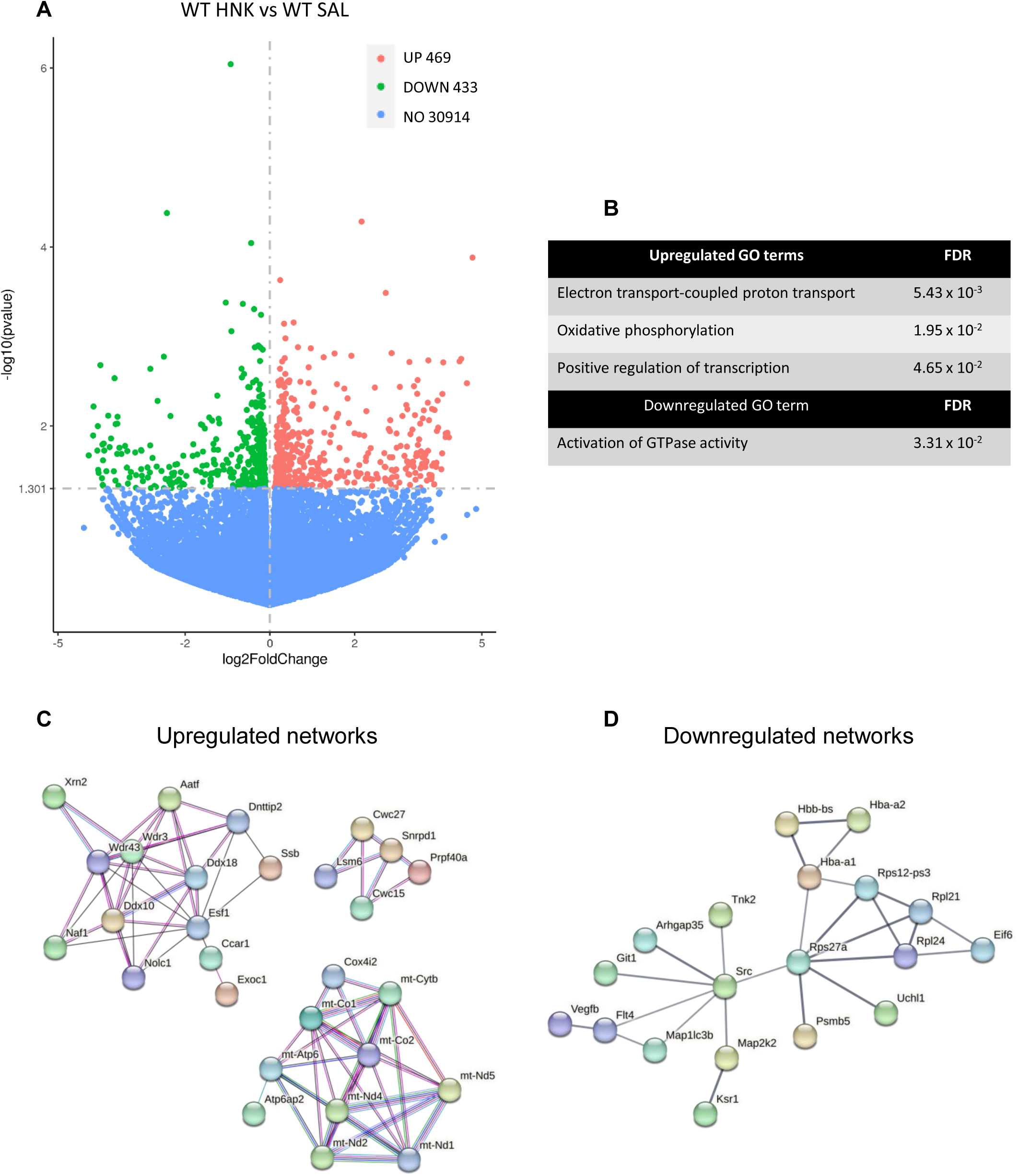
Treatment with HNK modifies the hippocampal transcriptional profile in mice. (A) Volcano plot from RNA-seq analysis of HNK-treated vs saline-treated WT mice. Downregulated genes are shown in green and upregulated genes in red (p < 0.05). (B) Gene ontology analysis (molecular function) for upregulated and downregulated genes (FDR < 0.05). (C, D) STRING analysis of upregulated and downregulated networks. The most connected nodes are shown (N = 6 mice per experimental group).

### HNK corrects synaptic plasticity and memory deficits in aged APP/PS1 mice

Our finding that HNK prevented synapse and memory deficits induced by AβOs prompted us to investigate whether HNK could reverse pre-existing impairments in synaptic plasticity and memory in aged (19-24-month-old) APP/PS1 mice, used as a transgenic model of late-stage AD. HNK (0.5 mg/kg, i.p.) or saline were administered daily for 12-14 days, and hippocampus-dependent spatial memory was tested 8-9 days after the onset of treatment using the two-day Radial Arm Water Maze (RAWM) paradigm, a memory task for which aged APP/PS1 are impaired [27]. Following the RAWM, we obtained hippocampal slices and measured LTP at Schäffer collaterals.

Remarkably, HNK reversed both the inhibition of hippocampal LTP (**Fig. 5A, B**) and spatial memory deficits (**Fig. 5C**) in aged APP/PS1 mice. Control measurements showed no differences in input/output (I/O) curves between WT and APP/PS1 mice, or between animals treated with vehicle or HNK (**Fig. S3A**). Additional control experiments revealed that treatment with HNK had no impact on body weight (**Fig. S3B**) or hippocampal Aβ load in APP/PS1 mice (**Fig. S3C**).

**Figure 5.**
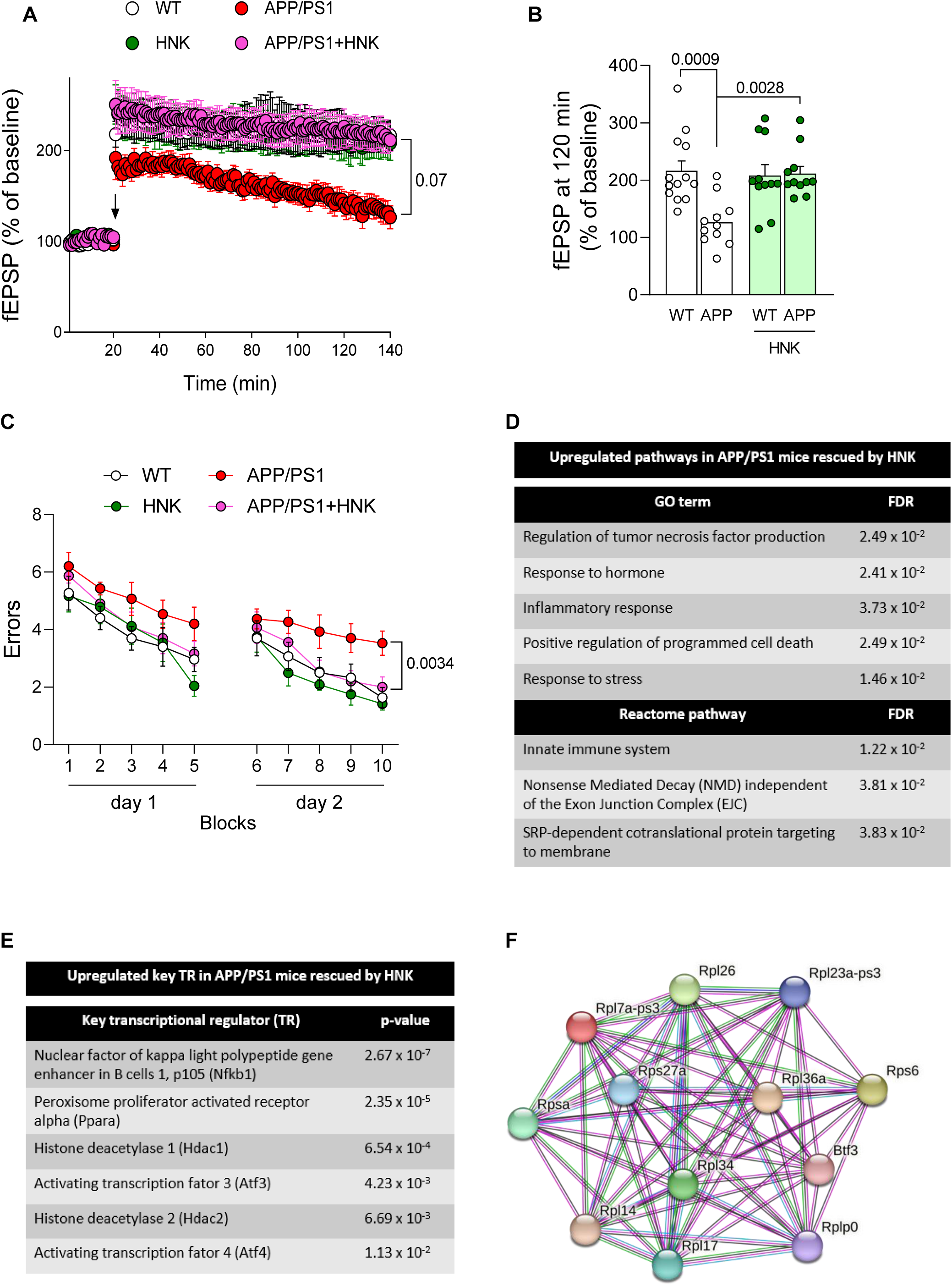
Treatment with HNK reverses LTP and memory deficits, and corrects hippocampal transcriptional alterations in APP/PS1 mice. (A) fEPSP recording in hippocampal slices from 19–24-month-old APP/PS1 (or WT) mice treated daily with HNK (0.5 mg/kg, i.p.) or vehicle for 12-14 days (N = 13 WT, 11 APP/PS1, 11 WT/HNK, 11 APP/PS1/HNK). N = number of slices (derived from at least 3 mice) per experimental condition. Arrows indicate tetanic stimulation. (B) fEPSP slope measured 120 min after tetanic stimulation. Two-way ANOVA followed by Tukey’s post-hoc test. Interaction: F (1, 42) = 8.872; p = 0.0048. (C) Spatial memory assessed using the Radial Arms Water Maze (RAWM) test on 19–24-month-old APP/PS1 (or WT) mice treated daily with HNK (0.5 mg/kg, i.p.) or vehicle for 8-10 days (N = 10 WT, 10 APP/PS1, 8 WT/HNK, and 10 APP/PS1/HNK). Statistical analysis was performed using the values of the testing day. p-values for appropriate comparisons are indicated; Two-way ANOVA followed by Tukey’s post-hoc test. (D-F) RNA-seq analyses of genes that were differentially expressed in saline-treated APP/PS1 (compared to WT mice) mice and whose expression was corrected by treatment with HNK. (D) Gene ontology (molecular function and reactome pathway). (E) Key transcription factor analysis by TRRUST (FDR < 0.05). (F) Representative STRING cluster of ribosome-linked genes that were differentially expressed in APP/PS1 (compared to WT mice) and were normalized by treatment with HNK. (N = 6 mice per experimental group).

### HNK rescues hippocampal transcriptional alterations in APP/PS1 mice

Finally, we investigated hippocampal transcriptional alterations induced by HNK in aged APP/PS1 mice. Mice were treated with HNK (0.5 mg/kg, i.p.) or saline daily for 14 days, and hippocampal transcriptomic changes were assessed by RNAseq. GO analyses revealed significantly upregulated pathways in APP/PS1 mice (compared to WT mice) that were corrected by HNK treatment (**Fig. 5D**). This included regulation of programmed cell death and response to hormones and stress (**Fig. 5D**). Reactome pathway analyses further implicated the innate immune system and, notably, three pathways associated with RNA metabolism and translation that were aberrantly regulated in APP/PS1 mice and were rescued by HNK. These pathways were nonsense-mediated RNA decay, co-translational protein targeting to membranes and formation of a pool of free 40S subunits (**Fig. 5D**). These are consistent with previous observations of neuroinflammation, deregulated hormonal signaling, TNF-α-dependent neuronal stress responses, and memory impairment in AD models [17, 28–30].

By performing Transcriptional Regulatory Relationships Unraveled by Sentence-based Text mining (TRRUST) analysis, we further identified transcriptional regulators known to control gene networks that HNK rescued in APP/PS1 mice. Among those regulators, we found molecules involved in inflammatory responses (NF-κB1, PPARα), transcriptional repression (HDAC1, HDAC2) and integrated stress response effectors (ATF3, ATF4) (**Fig. 5E**). Network analysis on STRING revealed a key cluster of ribosomal genes whose aberrant expression in APP/PS1 mice was corrected by HNK treatment (**Fig. 5F**).

In addition, we analyzed differentially expressed genes that were upregulated in APP/PS1 mice and were corrected by HNK treatment. Network analysis on STRING and GO revealed that HNK rescued transcriptional programs involved in cytosolic ribosomes and regulation of immune system process (**Fig. S4A, B**).

Downregulated genes in APP/PS1 mice that were corrected by HNK were also analyzed. We found that HNK rescued mRNA levels for genes coding for RNA polymerase II holoenzyme subunits, calcium channel complex and synaptic proteins in APP/PS1 mice (**Fig. S5A, B**). Altogether, these results indicate that HNK rescues transcriptional programs associated with inflammation, impaired proteostasis, calcium signaling and synaptic proteins in aged APP/S1 mice.

## DISCUSSION

In the present study, we investigated the effects of HNK, a ketamine metabolite whose mechanism of action is thought to involve activation of protein synthesis pathways [14], on hippocampal synaptic plasticity and memory in AD model mice. We demonstrate that HNK prevents deficits in hippocampal protein synthesis, LTP and memory induced by AβOs in mice. We further found that HNK triggers phosphorylation of mTOR, ERK1/2, S6K1, RSK and S6, implicated in protein synthesis, synaptic plasticity and memory [25, 31], in hippocampal slices. Interestingly, HNK-induced phosphorylation of S6 depended on mTOR activity, whereas rescue of LTP and memory in AβO-infused mice depended on ERK1/2, a key signaling effector that stimulates translation of immediate early genes necessary for synaptic plasticity [32, 33].

Even though HNK induced S6 phosphorylation, which may act as an inducer of mRNA translation and ribosome biogenesis, HNK *per se* did not increase global protein synthesis in hippocampal slices. In this regard, it should be noted that current evidence indicates that S6 phosphorylation *per se* does not increase mRNA translation [21, 34–36]. Moreover, even though LTP stimulates S6 phosphorylation [21], it has been reported that activation of mTORC1 signaling is not necessary for hippocampal LTP [37]. This may explain our finding that ERK1/2 signaling, but not mTOR, is required for the rescue of LTP impairment by HNK.

We further found that chronic treatment with HNK upregulate hippocampal genes involved in oxidative phosphorylation, in line with previous evidence indicating that mitochondrial energy metabolism and the antioxidant defense system mediate the actions of ketamine [26]. AD is associated with reduced energy metabolism and decreased expression of mitochondrial electron transport chain subunits [38], suggesting that rescuing mitochondrial function could be beneficial in AD. Of note, a very recent study reported that treatment with trazodone corrects translational attenuation of mitochondrial proteins in prion-diseased mice, and that this is accompanied by improved mitochondrial and synaptic function [39].

Strikingly, HNK reversed pre-existing impairments in synaptic plasticity and memory in aged (19–24-month-old) APP/PS1 mice. HNK further corrected aberrant transcriptional signatures related to inflammation and translation/proteostasis in APP/PS1 mice, suggesting that enduring changes in brain gene expression may contribute to the neuroprotective mechanisms of HNK. Our findings on hippocampal transcriptional changes *in vivo* further suggest that HNK impacts translation by upregulating translation-associated pathways, which may promote ribosome biogenesis and increase translational capacity. Collectively, these findings support the notion that HNK has therapeutic potential even in the presence of substantial brain deposition of Aβ.

In conclusion, our findings show that HNK alleviates synaptic deficits and memory impairment, and restores hippocampal protein synthesis, in AD mouse models. The results demonstrate that the mTOR and ERK1/2 signaling pathways mediate the neuroprotective actions of (*2R,6R*)-HNK by impacting energy metabolism, and inflammatory responses. HNK has been implicated as a mediator of the antidepressant actions of ketamine without the latter’s detrimental side effects [40], and is currently in clinical trials in treatment-resistant depression (NCT04711005). If HNK is proven safe and effective, like the related FDA-approved drug esketamine, this would raise the prospect that HNK could be repurposed for treatment of cognitive decline in AD.

## MATERIALS AND METHODS

### Animals

Adult male or female C57BL/6J mice (3-month-old) were obtained from the animal facility at McGill University and were housed in groups of 2-5 mice per cage, with food and water *ad libitum*, in rooms with controlled temperature (21°C) and humidity (∼55%) on a 12 h light/dark cycle. Male and female APPswe/PS1ΔE9 mice on a C57BL/6J background (Jackson Laboratories, strain code #005864) were bred at the animal facilities of McGill University and the Federal University of Rio de Janeiro. All procedures followed the Canadian Council on Animal Care guidelines and were approved by McGill University Committees and by the Institutional Animal Care and Utilization Committee of the Federal University of Rio de Janeiro.

### Biochemistry in hippocampal slices

Hippocampal slices (400 μm) were obtained and allowed to recover in artificial cerebrospinal fluid (aCSF; 124 mM NaCl, 4.4 mM KCl, 1 mM Na_2_ HPO_4_, 25 mM NaHCO_3_, 2 mM MgCl_2_, 2mM CaCl_2_, 10 mM glucose) for 2 h. For time course experiments, slices were treated with (*2R,6R*)-hydroxynorketamine (HNK) (Tocris Biosciences, #1430202-69-9; 20 μM) or saline for the time intervals indicated. Newly synthesized polypeptides were detected using surface sensing of translation (SUnSET), as described [41]. For SUnSET experiments, hippocampal slices from WT mice were treated with HNK (20 μM; 20 min) and then exposed to AβOs (1 μM; 2 h). Slices were then exposed to puromycin (5 µg/mL; 45 min) and collected for Western blotting. Puromycin incorporation was evaluated as a measure of newly synthesized polypeptides, and β–actin was used as loading control. A negative control lacking puromycin was added to each experiment.

For mechanistic studies using HNK, hippocampal slices from WT mice were pre-exposed to vehicle, PD98059 (MEK inhibitor; Sigma Aldrich, #P215; 20 μM) or rapamycin (mTORC1 inhibitor; Sigma Aldrich, R0395; 100 μM) for 30 min and then exposed to HNK (20 μM) for 10 min.

### Western blotting and ELISA

Hippocampal slices were dissociated using Bio-Plex Cell Lysis Kit (Bio-Rad), centrifuged for 10 min at 10,000 g at 4°C, and the supernatant was collected for Western blot analysis. Protein concentration was determined using Bradford Protein Assay Kit, and samples were prepared to a final concentration of 3 μg/μL. After boiling for 5 min, 50 μg of total protein were loaded per lane and resolved in 12% Tris-glycine SDS-PAGE gels. Proteins were then transferred to a nitrocellulose membrane and incubated with respective primary antibodies. Primary antibodies used were: puromycin (#MABE343, clone 12D10, 1:1,000; EMD Millipore), phospho-ERK1/2 Thr202/Tyr204 (#4370, 1:1,000; Cell Signaling), ERK1/2 (#9102, 1:1,000; Cell Signaling); phospho-p70S6K1 Thr389 (#9205, 1:1,000; Cell Signaling), p70S6K1 (#9206, 1:1,1000; Cell Signaling), phospho-S6 Ser235/236 (#4858, 1:1,000; Cell Signaling), phospho-S6 Ser240/244 (#5364, 1:1,000; Cell Signaling), S6 (#2217, 1:1,000; Cell Signaling), phospho-4E-BP1 Ser65 (#9451, 1:1,000; Cell Signaling), 4E-BP1/2 (#2845, 1:1,1000; Cell Signaling), phospho-mTOR Ser2448 (#2971, 1:1,1000; Cell Signaling), mTOR (#2983, clone 7C10, 1:1,1000; Cell Signaling), phospho-eIF2α Ser51 (#3597, 1:1,000; Invitrogen); eIF2α (#9722, 1:1,000; Cell Signaling) and β–actin (#A2228, 1:10,000; Sigma Aldrich). Immunoblots were developed by HRP-conjugated secondary antibodies (anti-mouse or anti-rabbit, #31430 or #31460, 1:5,000; Thermo Fisher) followed by ECL plus exposure, or by IR dye-conjugated fluorescent secondary antibodies (1:5,000; LiCor). When phosphorylated and total proteins were detected, membranes were initially probed for the phosphorylated epitope, then stripped with Restore stripping buffer (Thermo Fisher, 21059), and finally probed for total protein with the antibodies listed above. Quantification was performed using ImageJ (NIH). ELISA for human Aβ_42_ (Thermo Fisher; KHB3441) was performed according to manufacturer’s instructions, as previously described [19, 42].

### A**β**O preparation and intracerebroventricular (i.c.v.) infusion

Aβ_1-42_ peptide was purchased from California Peptide (Salt Lake, CA). Aβ oligomerization and quality control were performed as previously described [17, 42]. I.c.v. infusion of AβOs (10 pmol) in mice were performed as a one-time dose, as previously described [17, 43].

### Contextual Fear Conditioning Task

Mice were gently handled for 1 min/day for 3 days prior to behavioral experiments. To assess contextual fear memory, a two-phase protocol was used as described [44]. In the training phase, mice were placed in the apparatus, in which they received a foot shock (0.35 mA for 1 s) at 2 min. After 30 s, they received another foot shock (0.35 mA for 1 s), were kept in the apparatus for an additional 30 s, and then placed back in their home cages. For the test phase, carried out 24 h later, mice were placed in the same apparatus for 5 min, with no shock. Freezing behavior was recorded automatically on Freeze Frame (Harvard Apparatus, Holliston, MA) and was used as a memory index. For experiments using HNK, 20 mg/kg HNK (or saline) was injected i.p. 1 h prior to training. PD98059 (10 mg/kg, i.p.) or saline was injected 1h prior to HNK injection.

### Two-day Radial Arm Water Maze

Radial Arm Water Maze (RAWM) was performed in WT or APP/PS1 mice as described [14, 16]. Briefly, spatial memory was assessed by the ability to find a platform located in a defined arm of a 6-arm maze placed inside a circular pool. On day 1, fifteen trials were performed with alternation between visible and hidden platform. On day 2, all five blocks had the platform hidden. A training block comprises 3 trials of 60 s each. The number of entries into incorrect maze arms (errors) was recorded for each session and the number of errors per block was averaged.

### RNA sequencing

Total RNA was extracted using the RNeasy Micro kit (Qiagen), following manufacturer’s instructions. The purity of RNA preparations was checked using the 260/280 nm absorbance ratio. Only preparations with 260/280 nm absorbance ratios higher than 1.8 were used. RNA concentrations were determined by absorption at 260 nm. Messenger RNA was purified from total RNA using poly-T oligo-attached magnetic beads. After fragmentation, the first strand cDNA was synthesized using random hexamer primers, followed by second strand cDNA synthesis using either dUTP for directional library or dTTP for non-directional library. Library sequencing was performed on an Illumina platform and paired-end reads were generated. Quality control was performed; the index of the reference genome was built using Hisat2 v2.0.5 and paired-end clean reads were aligned to the reference genome using Hisat2 v2.0.5. featureCounts v1.5.0-p3 was used to count read numbers mapped to each gene. Differential expression analysis of two conditions/groups (two biological replicates per condition) was performed using the DESeq2 R package (1.20.0). The resulting p-values were adjusted using the Benjamini and Hochberg’s approach to control for false discovery rate (FDR). Genes with an adjusted FDR p-value ≤ 0.05 found by DESeq2 were considered differentially expressed. DEGs were analyzed using Gene Ontology (GO) (http://geneontology.org) using the mouse database. Pathway enrichment was analyzed by Fisher’s Exact test and corrected by FDR. For STRING analyses, DEGs were analyzed for all interaction sources except text mining, and limited to high confidence scores. DEG detected in saline-treated APP/PS1 but not in HNK-treated APP/PS1 mice (compared to control) were also used for STRING, GO and TRRUST analyses. DEGs were loaded onto the mouse Transcriptional Regulatory Relationships Unraveled by Sentence-based Text mining (TRRUST) database (https://www.grnpedia.org/trrust) and used for identification of putative master transcription factors. Results were considered significant with FDR values ≤ 0.05.

### Electrophysiology

Transverse hippocampal slices (400 μm) were obtained from 3-month-old C57BL/6 mice and placed on a perfusion chamber on aCSF (124 mM NaCl, 4.4 mM KCl, 1 mM Na_2_ HPO_4_, 25 mM NaHCO_3_, 2 mM MgCl_2_, 2mM CaCl_2_, 10 mM glucose). Slices were allowed to recover for at least 90 min, and were perfused with 20 µM HNK or saline for 20 min, followed by AβOs (400 nM) or vehicle for another 20 min. Stimulation was performed in CA3, and field excitatory postsynaptic potentials (fEPSPs) were recorded from hippocampal CA1 region as described [42]. For mechanistic studies, hippocampal slices were pre-exposed to vehicle, PD98059 (Sigma Aldrich, #P215; 20 μM) or rapamycin (Sigma Aldrich, R0395; 100μM) for 20 min and then exposed to HNK or vehicle (20μM), followed by AβOs (1 μM) for 20 min prior to tetanic stimulation.

fEPSPs were also recorded in hippocampal slices from 19–24-month-old WT or APP/PS1 mice that were daily treated with vehicle or HNK (0.5 mg/kg, i.p.) for 12-14 days prior to slice preparation.

### Statistical analysis

Data are expressed as means ± S.E.M. and were analyzed using GraphPad Prism 8 software (La Jolla, CA). Sample sizes and statistical tests for each experiment are indicated in the corresponding Figure Legends. p-values for relevant comparisons are indicated in Figures or Figure Legends.

## ACKNOWLEDGMENTS

This work was supported by grants from Fundação Carlos Chagas Filho de Amparo à Pesquisa do Estado do Rio de Janeiro (FAPERJ) (STF, FGDF and MVL), Conselho Nacional de Desenvolvimento Científico e Tecnológico (CNPq) (STF, FGDF and MVL), National Institute of Translational Neuroscience (INNT/Brazil) (STF and FGDF), Alzheimer’s Society Canada (NS and STF), International Brain Research Organization (IBRO Regions Connecting Awards to MVL and AAV), Alzheimer’s Association (AARF-21-848798 to FCR; AARG-D-615714 and the Blas Frangione Early Career Achievement Award to MVL), National Institutes of Health (NIH-NINDS/R01NS049442 to OA), Serrapilheira Institute (R-2012-37967 to MVL), and by travel grants from International Society for Neurochemistry (ISN), International Union of Biochemistry and Molecular Biology (IUBMB), American Society for Biochemistry and Molecular Biology (ASBMB) and Company of Biologists (to FCR). JCL holds the Canada Research Chair in Cellular and Molecular Neurophysiology (CRC 950-231066). We thank Dr. Newton G. Castro (Federal University of Rio de Janeiro) for access to equipment in his laboratory and for help with experiments in acute hippocampal slices.

## Author contributions

F.C.R., D.C., A.A.-.V. and S.T.F. designed the study. F.C.R., D.C., E.K.A, A.S., S.B., S.W., J.D.C., R.S-N., M.C., E.A. and A.A.-.V. performed research. F.C.R., D.C., O.A. and A.A.-.V. analyzed data. K.N., M.V.L., J.C.-.L., O.A., F.G.D.F., A.A.-.V., N.S. and S.T.F. contributed reagents, materials, animals and analysis tools. F.C.R., D.C., F.G.D.F.., M.V.L., A.A.-.V., N.S. and S.T.F. analyzed and discussed results. F.C.R., D.C., M.V.L. and S.T.F. wrote the manuscript with inputs from the other authors, notably by N.S. and A.A.-V.

## Data and materials availability

All data supporting the conclusions are presented in the manuscript and main Figures, or in Supplementary Materials.

---

**Figure S1.**
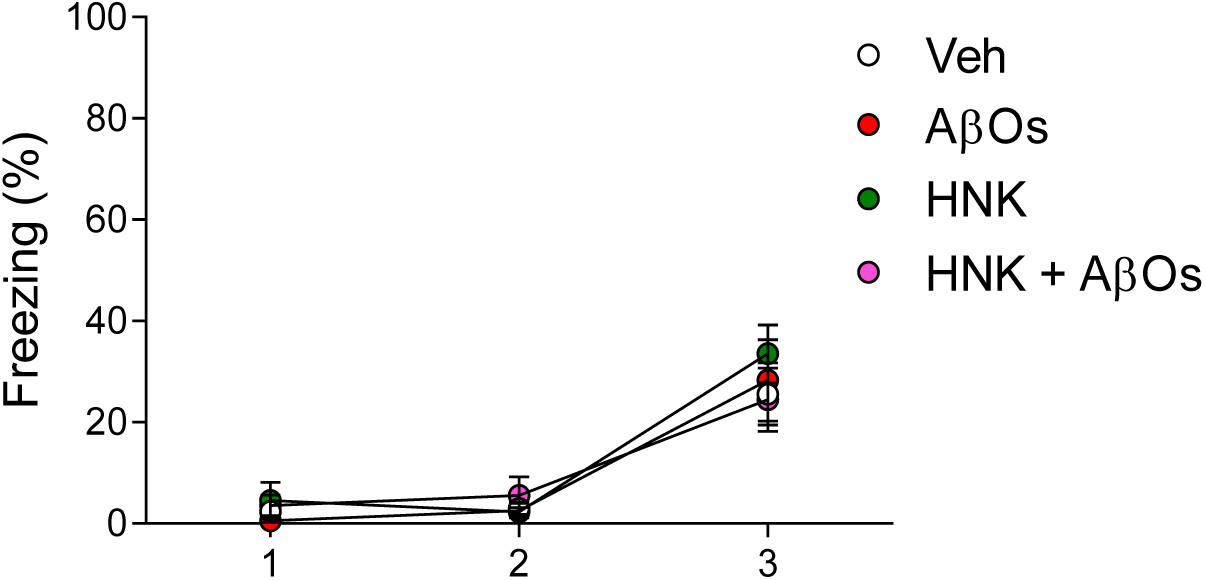
Freezing responses during contextual fear conditioning (CFC) training. Percentage of freezing during minutes 1, 2 (pre-shock) or 3 (during/after shock) in WT mice infused with AβOs and/or HNK (N = 7 Veh, 8 AβOs, 9 Veh/HNK, 10 AβOs/HNK). Two-way ANOVA followed by Sidak’s post-hoc test.

**Figure S2.**
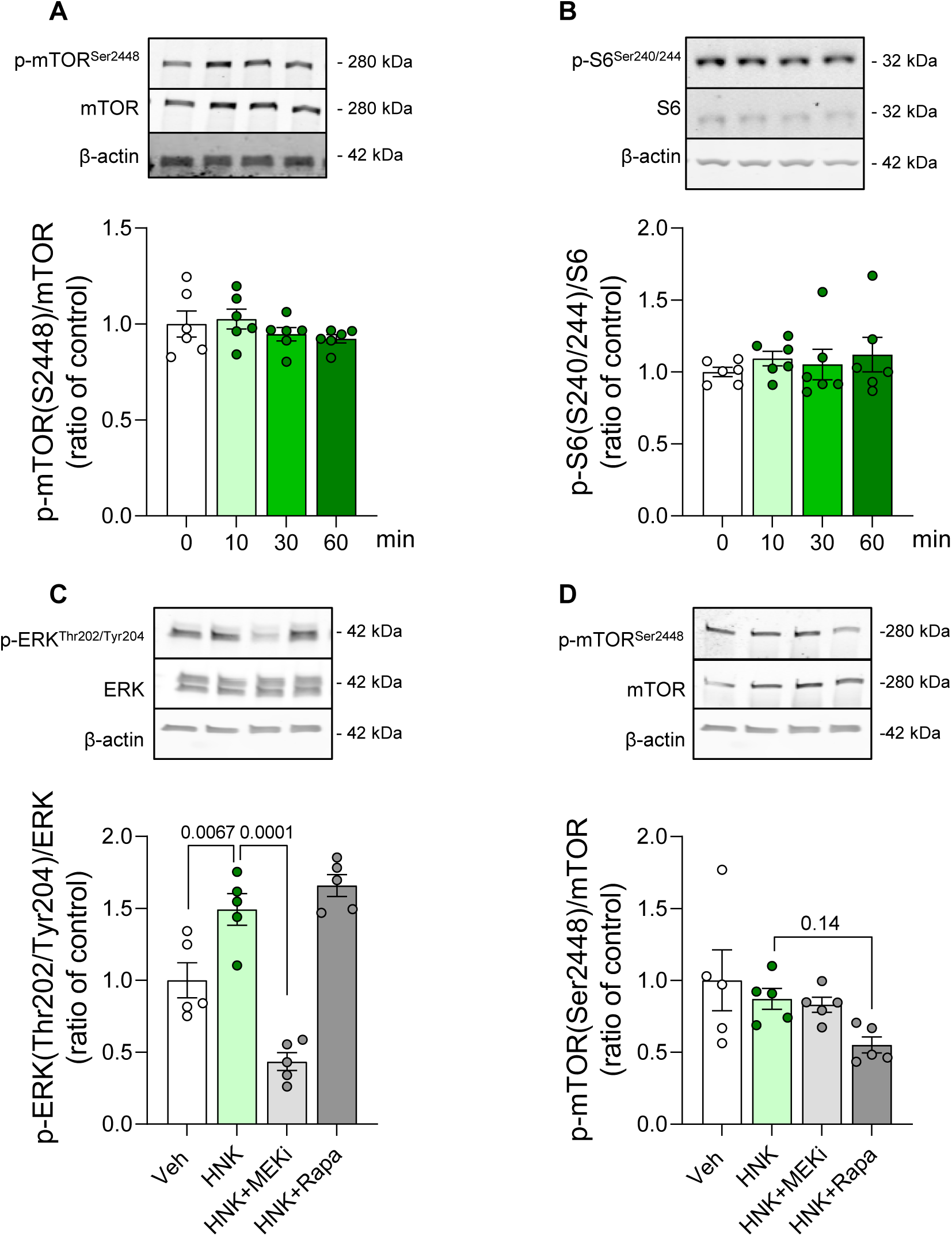
Hippocampal signaling pathways not altered by HNK. (A-B) Acute hippocampal slices were treated with vehicle or HNK (20 μM) for 10, 30 or 60 min. Phosphorylated mTOR and S6 (Ser240/244) were evaluated by Western blotting and normalized by loading control or by the respective total protein levels when appropriate (N = 6 slices/experimental group). One-way ANOVA followed by Dunnett’s post-hoc test. (C, D) Acute hippocampal slices were treated with vehicle, rapamycin (Rapa; 100 μM) or PD98059 (MEKi; 20 μM) for 30 min and were then exposed to vehicle or HNK (20 μM) for 10 min. Phosphorylated mTOR and ERK were evaluated by Western blotting and normalized by respective total protein levels (N = 5 slices/experimental group). p-values are shown for appropriate comparisons; One-way ANOVA followed by Sidak’s post-hoc test.

**Figure S3.**
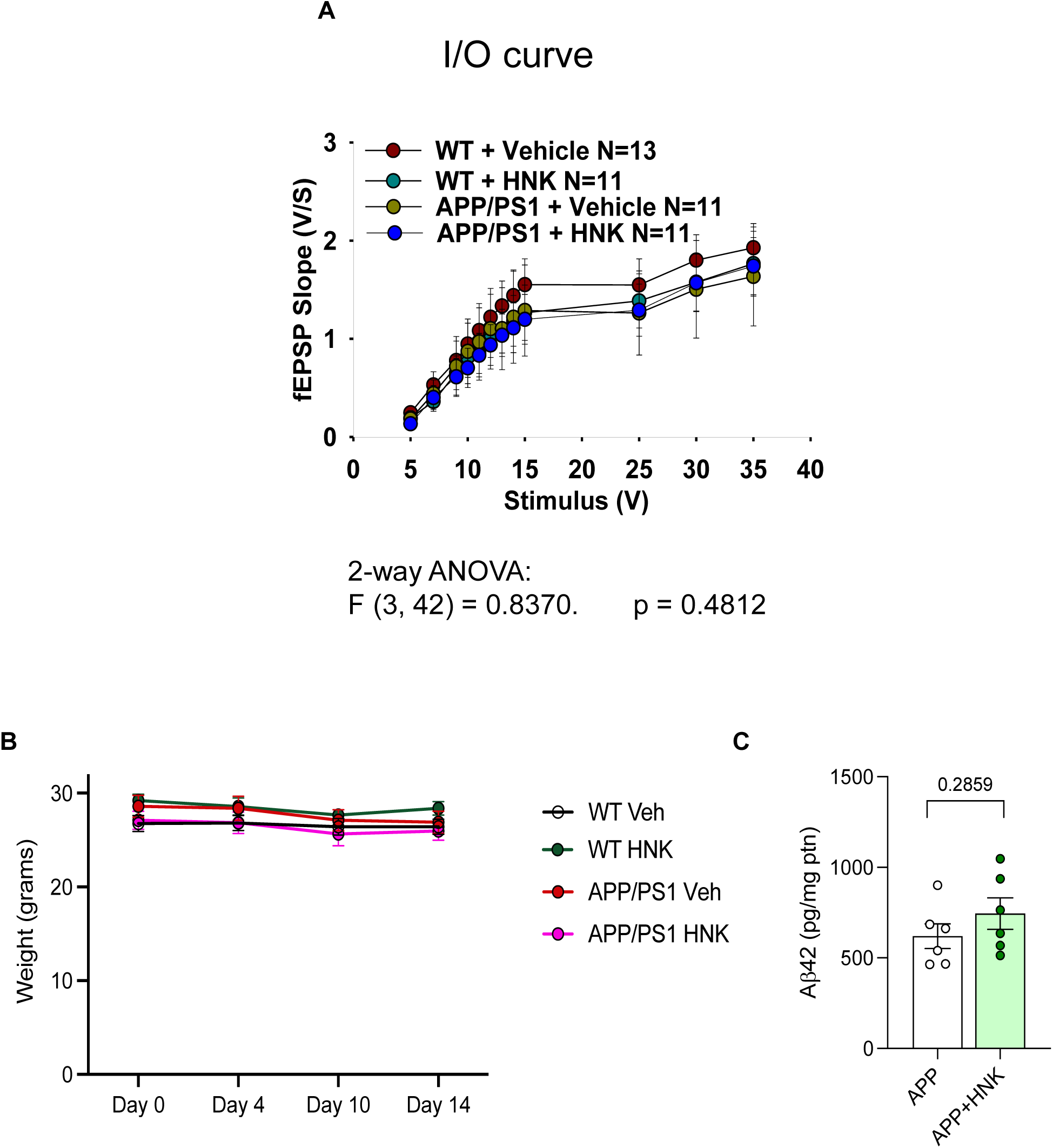
Treatment with HNK does not alter hippocampal basal synaptic transmission, body weight, or hippocampal Aβ load. (A) Input/output curves were analyzed in hippocampal slices from saline-or HNK-treated mice (WT or APP/PS1). (B) Body weights of saline-or HNK-treated mice (WT or APP/PS1) were evaluated on days 0, 4, 10 and 14 after daily HNK (0.5 mg/kg, i.p.) administration. Two-way ANOVA followed by Tukey’s post-hoc test. (C) Hippocampal Aβ_42_ was measured by ELISA in saline-or HNK-treated APP/PS1 mice (N = 6 mice/experimental group); Student t-test.

**Figure S4.**
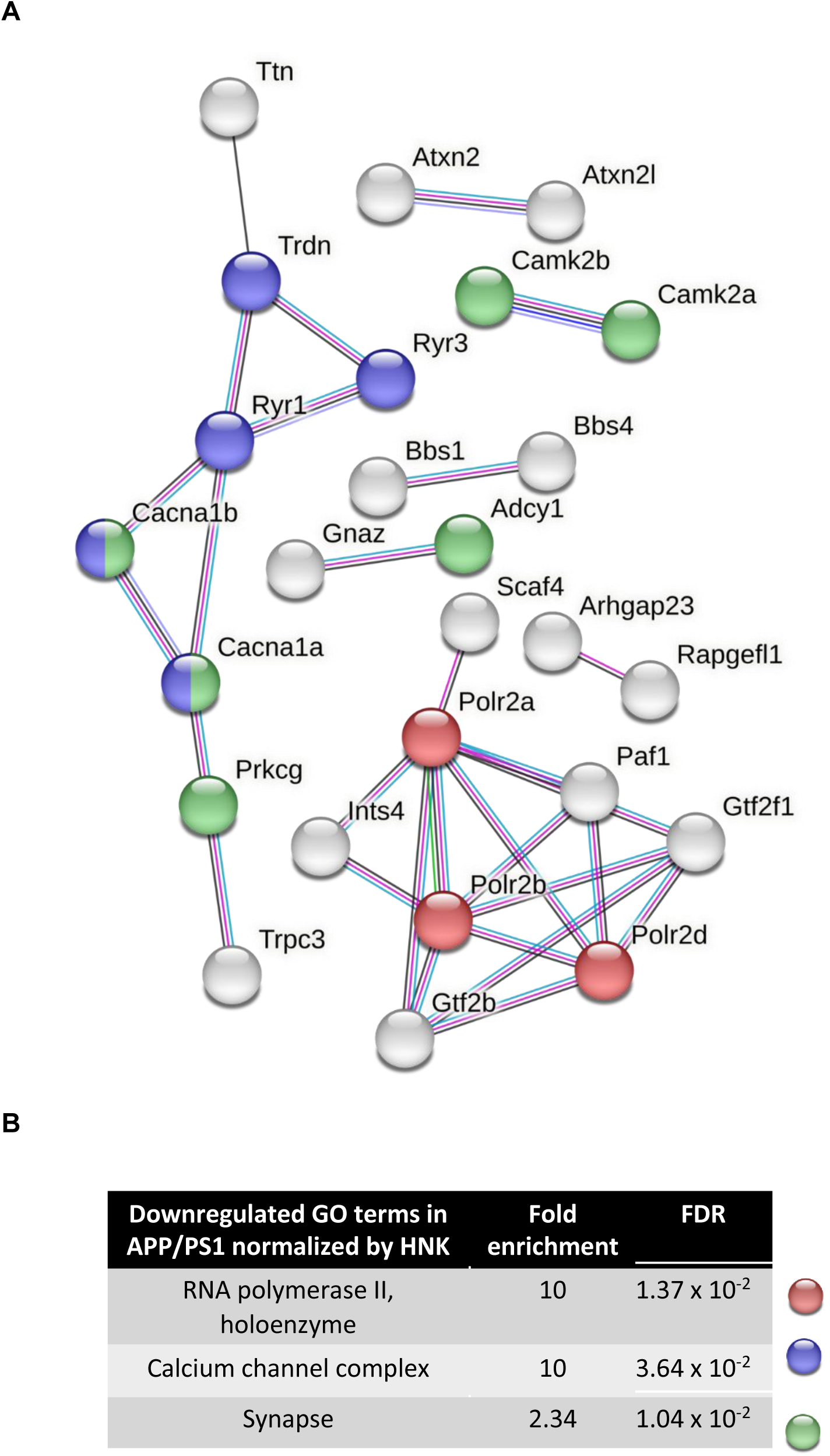
Treatment with HNK corrects the expression of cytosolic ribosome genes and genes involved in programmed cell death and immune system processes in APP/PS1 mice. (A) STRING analyses of genes that were upregulated in saline-treated APP/PS1 (compared to WT littermates) that were corrected by treatment with HNK. The most connected nodes are shown. Genes coding for ribosomal subunits are shown in Fig. 6F. The gene network is color-coded by GO terms. (B) GO analysis using STRING. (N = 6 mice per experimental group).

**Figure S5.**
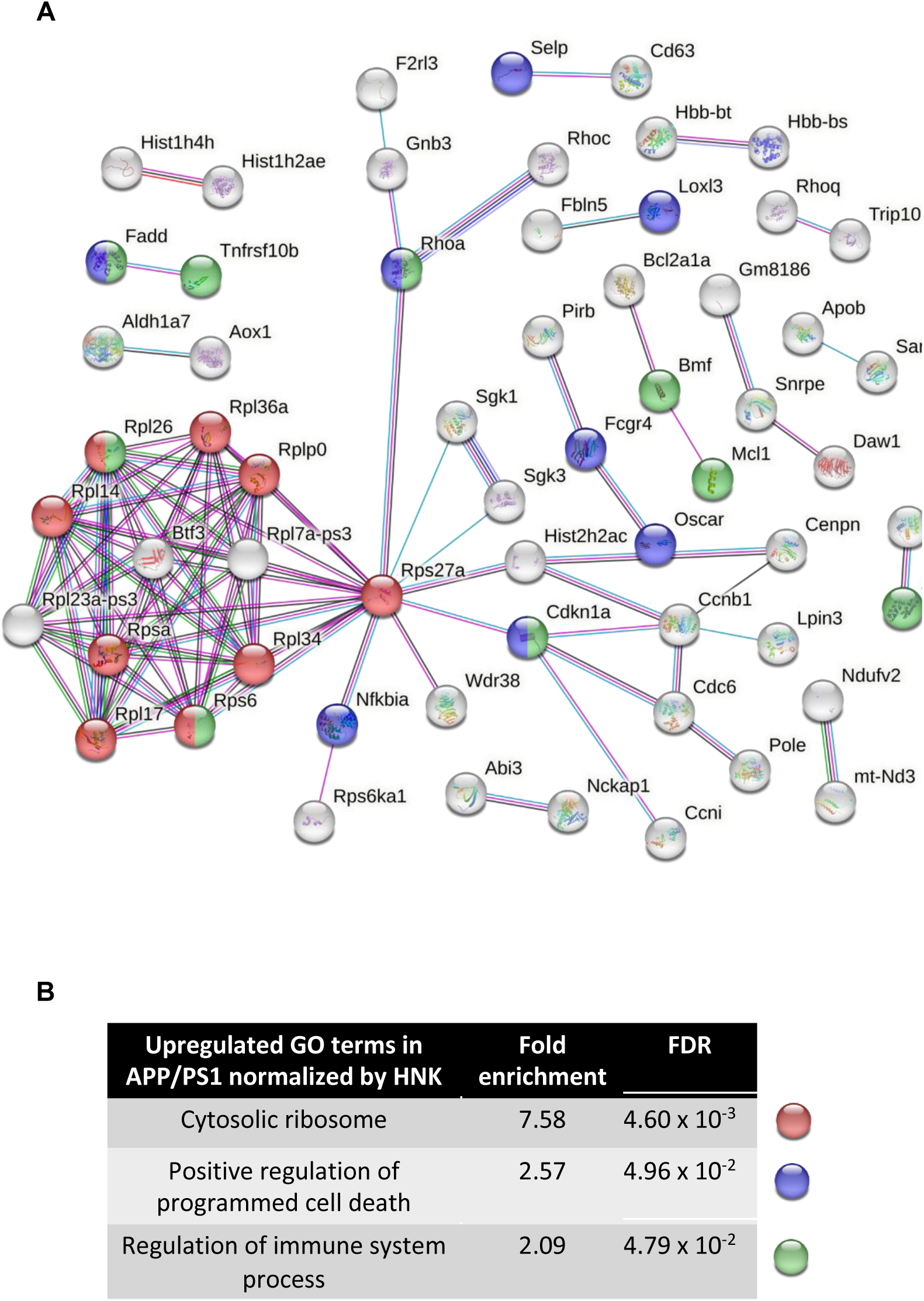
Treatment with HNK corrects the expression of RNA polymerase II, calcium channel complex and synaptic genes in APP/PS1 mice. (A) STRING analyses of genes that were downregulated in saline-treated APP/PS1 mice (compared to WT littermates) that were corrected by treatment with HNK. The most connected nodes are shown. The gene network is color-coded by GO terms. (B) GO analysis using STRING. (N = 6 mice per experimental group).

**Table S1. Differentially expressed genes (DEGs) in WT mice treated with HNK.**

**Table S2. Differentially expressed genes (DEGs) in APP/PS1 mice (compared to WT littermates) that were corrected by treatment with HNK.**

