## Supplemental Table 1 for "The ketamine metabolite (*2R,6R*)-hydroxynorketamine rescues hippocampal mRNA translation, synaptic plasticity and memory in mouse models of Alzheimer’s disease"

### Supplementary Table 1: Differentially expressed genes (DEGs) in WT mice treated with HNK

| gene_name | log2FoldChange | pvalue | gene_chr | gene_biotype | gene_description |
|---|---|---|---|---|---|
| Hba-a2 | -0,9240 | 9,070E+07 | 11 | protein_coding | hemoglobin alpha, adult chain 2 [Source:MGI Symbol;Acc:MGI:96016] |
| Capn11 | -2,4280 | 4,178E+09 | 17 | protein_coding | calpain 11 [Source:MGI Symbol;Acc:MGI:1352490] |
| C7 | 2,1647 | 5,203E+09 | 15 | protein_coding | complement component 7 [Source:MGI Symbol;Acc:MGI:88235] |
| Zbed3 | -0,4412 | 9,041E+09 | 13 | protein_coding | zinc finger, BED type containing 3 [Source:MGI Symbol;Acc:MGI:1919364] |
| Gm48230 | 4,7806 | 1,313E-04 | 12 | TEC | predicted gene, 48230 [Source:MGI Symbol;Acc:MGI:6097634] |
| Npm1 | 0,2440 | 2,346E-04 | 11 | protein_coding | nucleophosmin 1 [Source:MGI Symbol;Acc:MGI:106184] |
| Gm35248 | 2,7337 | 3,258E-04 | 15 | lincRNA | predicted gene, 35248 [Source:MGI Symbol;Acc:MGI:5594407] |
| Tnfrsf8 | -1,0392 | 4,183E-04 | 4 | protein_coding | tumor necrosis factor receptor superfamily, member 8 [Source:MGI Symbol;Acc:MGI:99908] |
| Srpk3 | -0,6369 | 4,320E-04 | X | protein_coding | serine/arginine-rich protein specific kinase 3 [Source:MGI Symbol;Acc:MGI:1891338] |
| Vcp-rs | -0,3715 | 4,943E-04 | X | processed_pseudogene | valosin containing protein, related sequence [Source:MGI Symbol;Acc:MGI:894298] |
| Smg5 | -0,2083 | 5,722E-04 | 3 | protein_coding | Smg-5 homolog, nonsense mediated mRNA decay factor (C, elegans) [Source:MGI Symbol;Acc:MGI:2447364] |
| Jpx | 0,5613 | 6,978E-04 | X | lincRNA | Jpx transcript, Xist activator (non-protein coding) [Source:MGI Symbol;Acc:MGI:2180008] |
| Klf10 | 0,3363 | 7,201E-04 | 15 | protein_coding | Kruppel-like factor 10 [Source:MGI Symbol;Acc:MGI:1101353] |
| Mutyh | -0,9063 | 8,727E-04 | 4 | protein_coding | mutY DNA glycosylase [Source:MGI Symbol;Acc:MGI:1917853] |
| Trim59 | 0,3670 | 1,049E-03 | 3 | protein_coding | tripartite motif-containing 59 [Source:MGI Symbol;Acc:MGI:1914199] |
| Zgpat | -0,2715 | 1,262E-03 | 2 | protein_coding | zinc finger, CCCH-type with G patch domain [Source:MGI Symbol;Acc:MGI:2449939] |
| Catspere2 | 0,6629 | 1,317E-03 | 1 | protein_coding | catsper channel auxiliary subunit epsilon 2 [Source:MGI Symbol;Acc:MGI:5589632] |
| Nrde2 | -0,3533 | 1,319E-03 | 12 | protein_coding | nrde-2 necessary for RNA interference, domain containing [Source:MGI Symbol;Acc:MGI:2670969] |
| Zdhhc18 | -0,2251 | 1,324E-03 | 4 | protein_coding | zinc finger, DHHC domain containing 18 [Source:MGI Symbol;Acc:MGI:3527792] |
| Gm44623 | 0,9667 | 1,355E-03 | 7 | antisense | predicted gene 44623 [Source:MGI Symbol;Acc:MGI:5753199] |
| Arhgef4 | -0,1676 | 1,401E-03 | 1 | protein_coding | Rho guanine nucleotide exchange factor (GEF) 4 [Source:MGI Symbol;Acc:MGI:2442507] |
| Gm37048 | 2,8720 | 1,534E-03 | 1 | TEC | predicted gene, 37048 [Source:MGI Symbol;Acc:MGI:5610276] |
| Efcab9 | 1,5272 | 1,547E-03 | 11 | protein_coding | EF-hand calcium binding domain 9 [Source:MGI Symbol;Acc:MGI:1916556] |
| Gm14391 | 1,9237 | 1,642E-03 | 2 | protein_coding | predicted gene 14391 [Source:MGI Symbol;Acc:MGI:3709324] |
| Skor1 | -2,5020 | 1,680E-03 | 9 | protein_coding | SKI family transcriptional corepressor 1 [Source:MGI Symbol;Acc:MGI:2443473] |

| | | | | | |
|---|---|---|---|---|---|
| B3gnt7 | 1,2698 | 1,705E-03 | 1 | protein_coding | UDP-GlcNAc:betaGal beta-1,3-N-acetylglucosaminyltransferase 7 [Source:MGI Symbol;Acc:MGI:2384394] |
| Gm6563 | 0,3523 | 1,745E-03 | 19 | protein_coding | predicted pseudogene 6563 [Source:MGI Symbol;Acc:MGI:3646907] |
| Gm45667 | 4,5134 | 1,780E-03 | 7 | TEC | predicted gene 45667 [Source:MGI Symbol;Acc:MGI:5791503] |
| Gm10252 | 3,7398 | 1,849E-03 | 7 | TEC | predicted gene 10252 [Source:MGI Symbol;Acc:MGI:3704292] |
| St6galnac4 | -0,2322 | 1,877E-03 | 2 | protein_coding | ST6 (alpha-N-acetyl-neuraminyl-2,3-beta-galactosyl-1,3)-N-acetylgalactosaminide alpha-2,6-sialyltransferase 4 [Source:MGI Symbol;Acc:MGI:1341894] |
| 4930563M21Rik | 4,4761 | 1,894E-03 | 9 | protein_coding | RIKEN cDNA 4930563M21 gene [Source:MGI Symbol;Acc:MGI:1922508] |
| mt-Nd5 | 0,2907 | 1,900E-03 | MT | protein_coding | mitochondrially encoded NADH dehydrogenase 5 [Source:MGI Symbol;Acc:MGI:102496] |
| Gm15414 | 3,2929 | 1,929E-03 | 7 | antisense | predicted gene 15414 [Source:MGI Symbol;Acc:MGI:3705307] |
| Gm9748 | 4,0909 | 1,947E-03 | 14 | antisense | predicted gene 9748 [Source:MGI Symbol;Acc:MGI:3642166] |
| Gm34280 | -3,9976 | 2,090E-03 | 7 | antisense | predicted gene, 34280 [Source:MGI Symbol;Acc:MGI:5593439] |
| Xrra1 | 0,9157 | 2,208E-03 | 7 | protein_coding | X-ray radiation resistance associated 1 [Source:MGI Symbol;Acc:MGI:2181647] |
| Tmem200b | -0,6670 | 2,290E-03 | 4 | protein_coding | transmembrane protein 200B [Source:MGI Symbol;Acc:MGI:3646343] |
| mt-Nd4 | 0,2398 | 2,291E-03 | MT | protein_coding | mitochondrially encoded NADH dehydrogenase 4 [Source:MGI Symbol;Acc:MGI:102498] |
| Ltb4r1 | -2,8193 | 2,294E-03 | 14 | protein_coding | leukotriene B4 receptor 1 [Source:MGI Symbol;Acc:MGI:1309472] |
| Rif1 | 0,3645 | 2,622E-03 | 2 | protein_coding | replication timing regulatory factor 1 [Source:MGI Symbol;Acc:MGI:1098622] |
| Gm10819 | -0,6087 | 2,624E-03 | 19 | processed_pseudogene | predicted gene 10819 [Source:MGI Symbol;Acc:MGI:3642135] |
| Alas2 | -0,6588 | 2,864E-03 | X | protein_coding | aminolevulinic acid synthase 2, erythroid [Source:MGI Symbol;Acc:MGI:87990] |
| Cldn15 | -3,6633 | 2,923E-03 | 5 | protein_coding | claudin 15 [Source:MGI Symbol;Acc:MGI:1913103] |
| Btaf1 | 0,2625 | 3,043E-03 | 19 | protein_coding | B-TFIID TATA-box binding protein associated factor 1 [Source:MGI Symbol;Acc:MGI:2147538] |
| Gm22613 | 3,4867 | 3,068E-03 | 2 | misc_RNA | predicted gene, 22613 [Source:MGI Symbol;Acc:MGI:5452390] |
| Pisd-ps1 | -0,3245 | 3,076E-03 | 11 | transcribed_unprocessed_pseudogene | phosphatidylserine decarboxylase, pseudogene 1 [Source:MGI Symbol;Acc:MGI:3842428] |
| Xrn2 | 0,2569 | 3,081E-03 | 2 | protein_coding | 5'-3' exoribonuclease 2 [Source:MGI Symbol;Acc:MGI:894687] |
| Rfesd | 0,4593 | 3,088E-03 | 13 | protein_coding | Rieske (Fe-S) domain containing [Source:MGI Symbol;Acc:MGI:2145198] |
| Snrpd1 | 0,2448 | 3,124E-03 | 18 | protein_coding | small nuclear ribonucleoprotein D1 [Source:MGI Symbol;Acc:MGI:98344] |
| mt-Cytb | 0,2014 | 3,183E-03 | MT | protein_coding | mitochondrially encoded cytochrome b [Source:MGI Symbol;Acc:MGI:102501] |
| Haus6 | 0,4209 | 3,193E-03 | 4 | protein_coding | HAUS augmin-like complex, subunit 6 [Source:MGI Symbol;Acc:MGI:1923389] |
| Tnfsf10 | 0,7213 | 3,208E-03 | 3 | protein_coding | tumor necrosis factor (ligand) superfamily, member 10 [Source:MGI Symbol;Acc:MGI:107414] |

| | | | | | |
|---|---|---|---|---|---|
| Zfp800 | 0,4506 | 3,248E-03 | 6 | protein_coding | zinc finger protein 800 [Source:MGI Symbol;Acc:MGI:1889334] |
| Hgh1 | -0,3963 | 3,256E-03 | 15 | protein_coding | HGH1 homolog [Source:MGI Symbol;Acc:MGI:1930628] |
| Grip1 | 0,2830 | 3,316E-03 | 10 | protein_coding | glutamate receptor interacting protein 1 [Source:MGI Symbol;Acc:MGI:1921303] |
| Gm26901 | 0,9155 | 3,320E-03 | 1 | lincRNA | predicted gene, 26901 [Source:MGI Symbol;Acc:MGI:5477395] |
| Gm43815 | 4,6494 | 3,326E-03 | 5 | TEC | predicted gene 43815 [Source:MGI Symbol;Acc:MGI:5663952] |
| Zfp788 | 0,2941 | 3,343E-03 | 7 | protein_coding | zinc finger protein 788 [Source:MGI Symbol;Acc:MGI:1914857] |
| Smarca1 | 0,3021 | 3,399E-03 | X | protein_coding | SWI/SNF related, matrix associated, actin dependent regulator of chromatin, subfamily a, member 1 [Source:MGI Symbol;Acc:MGI:19351[illegible] |
| 5330406M23Rik | 1,5829 | 3,413E-03 | 6 | TEC | RIKEN cDNA 5330406M23 gene [Source:MGI Symbol;Acc:MGI:1923921] |
| Zdhhc14 | -0,2346 | 3,427E-03 | 17 | protein_coding | zinc finger, DHHC domain containing 14 [Source:MGI Symbol;Acc:MGI:2653229] |
| mt-Nd1 | 0,2188 | 3,521E-03 | MT | protein_coding | mitochondrially encoded NADH dehydrogenase 1 [Source:MGI Symbol;Acc:MGI:101787] |
| Gm47023 | 3,4005 | 3,590E-03 | 10 | TEC | predicted gene, 47023 [Source:MGI Symbol;Acc:MGI:6095711] |
| Gm3943 | 3,0673 | 3,656E-03 | X | processed_pseudogene | predicted gene 3943 [Source:MGI Symbol;Acc:MGI:3782117] |
| Dek | 0,3434 | 3,660E-03 | 13 | protein_coding | DEK oncogene (DNA binding) [Source:MGI Symbol;Acc:MGI:1926209] |
| Gm17586 | -0,5101 | 3,673E-03 | 11 | lincRNA | predicted gene, 17586 [Source:MGI Symbol;Acc:MGI:4937220] |
| Saysd1 | -0,4747 | 3,699E-03 | 14 | protein_coding | SAYSVFN motif domain containing 1 [Source:MGI Symbol;Acc:MGI:1914759] |
| Gm34552 | 2,4781 | 3,718E-03 | 12 | lincRNA | predicted gene, 34552 [Source:MGI Symbol;Acc:MGI:5593711] |
| Gm44684 | 1,7570 | 3,818E-03 | 7 | processed_transcript | predicted gene 44684 [Source:MGI Symbol;Acc:MGI:5753260] |
| Tctn1 | -0,2564 | 3,899E-03 | 5 | protein_coding | tectonic family member 1 [Source:MGI Symbol;Acc:MGI:3603820] |
| Gm10222 | 1,0408 | 4,163E-03 | 1 | unprocessed_pseudogene | predicted gene 10222 [Source:MGI Symbol;Acc:MGI:3642643] |
| Triobp | 0,3796 | 4,356E-03 | 15 | protein_coding | TRIO and F-actin binding protein [Source:MGI Symbol;Acc:MGI:1349410] |
| B130006D01Rik | 3,4762 | 4,369E-03 | 11 | protein_coding | RIKEN cDNA B130006D01 gene [Source:MGI Symbol;Acc:MGI:2444371] |
| Gm23608 | 4,0714 | 4,381E-03 | 2 | snoRNA | predicted gene, 23608 [Source:MGI Symbol;Acc:MGI:5453385] |
| Cant1 | -0,1822 | 4,472E-03 | 11 | protein_coding | calcium activated nucleotidase 1 [Source:MGI Symbol;Acc:MGI:1923275] |
| Ltbp2 | -1,2405 | 4,580E-03 | 12 | protein_coding | latent transforming growth factor beta binding protein 2 [Source:MGI Symbol;Acc:MGI:99502] |
| Zfp938 | 0,2846 | 4,646E-03 | 10 | protein_coding | zinc finger protein 938 [Source:MGI Symbol;Acc:MGI:3621440] |
| Galt | -0,2516 | 4,712E-03 | 4 | protein_coding | galactose-1-phosphate uridyl transferase [Source:MGI Symbol;Acc:MGI:95638] |
| Tlr3 | 0,3290 | 4,882E-03 | 8 | protein_coding | toll-like receptor 3 [Source:MGI Symbol;Acc:MGI:2156367] |

| | | | | | |
|---|---|---|---|---|---|
| Ghr | 0,5123 | 4,906E-03 | 15 | protein_coding | growth hormone receptor [Source:MGI Symbol;Acc:MGI:95708] |
| Zfp973 | 1,8323 | 4,918E-03 | 2 | protein_coding | zinc finger protein 973 [Source:MGI Symbol;Acc:MGI:3615331] |
| Pms1 | 0,5727 | 5,088E-03 | 1 | protein_coding | PMS1 homolog 1, mismatch repair system component [Source:MGI Symbol;Acc:MGI:1202302] |
| Gm11186 | 3,9566 | 5,200E-03 | 11 | antisense | predicted gene 11186 [Source:MGI Symbol;Acc:MGI:3649729] |
| Prss41 | -2,6484 | 5,246E-03 | 17 | protein_coding | protease, serine 41 [Source:MGI Symbol;Acc:MGI:1918253] |
| Tbc1d16 | -0,1683 | 5,263E-03 | 11 | protein_coding | TBC1 domain family, member 16 [Source:MGI Symbol;Acc:MGI:2652878] |
| Zmym6 | 0,2215 | 5,358E-03 | 4 | protein_coding | zinc finger, MYM-type 6 [Source:MGI Symbol;Acc:MGI:106505] |
| Ydjc | -0,3339 | 5,457E-03 | 16 | protein_coding | YdjC homolog (bacterial) [Source:MGI Symbol;Acc:MGI:1916351] |
| Efcc1 | -0,4189 | 5,487E-03 | 6 | protein_coding | EF hand and coiled-coil domain containing 1 [Source:MGI Symbol;Acc:MGI:3611451] |
| Gm45486 | 3,7269 | 5,499E-03 | 7 | antisense | predicted gene 45486 [Source:MGI Symbol;Acc:MGI:5791322] |
| Itga7 | -0,2562 | 5,577E-03 | 10 | protein_coding | integrin alpha 7 [Source:MGI Symbol;Acc:MGI:102700] |
| Coq4 | -0,2974 | 5,600E-03 | 2 | protein_coding | coenzyme Q4 [Source:MGI Symbol;Acc:MGI:1098826] |
| Pcdh15 | 0,3474 | 5,650E-03 | 10 | protein_coding | protocadherin 15 [Source:MGI Symbol;Acc:MGI:1891428] |
| Dnaaf2 | 1,0092 | 5,699E-03 | 12 | protein_coding | dynein, axonemal assembly factor 2 [Source:MGI Symbol;Acc:MGI:1923566] |
| Pdlim2 | -0,4292 | 5,714E-03 | 14 | protein_coding | PDZ and LIM domain 2 [Source:MGI Symbol;Acc:MGI:2384850] |
| Ticrr | 3,2186 | 5,823E-03 | 7 | protein_coding | TOPBP1-interacting checkpoint and replication regulator [Source:MGI Symbol;Acc:MGI:1924261] |
| Chst12 | -0,2612 | 5,834E-03 | 5 | protein_coding | carbohydrate sulfotransferase 12 [Source:MGI Symbol;Acc:MGI:1929064] |
| Gm11837 | 0,9882 | 5,970E-03 | 4 | antisense | predicted gene 11837 [Source:MGI Symbol;Acc:MGI:3702175] |
| Grk6 | -0,1800 | 5,997E-03 | 13 | protein_coding | G protein-coupled receptor kinase 6 [Source:MGI Symbol;Acc:MGI:1347078] |
| Mir7237 | -4,1601 | 6,053E-03 | 8 | miRNA | microRNA 7237 [Source:MGI Symbol;Acc:MGI:5530864] |
| Grik4 | -0,1893 | 6,066E-03 | 9 | protein_coding | glutamate receptor, ionotropic, kainate 4 [Source:MGI Symbol;Acc:MGI:95817] |
| Particl | -0,3219 | 6,081E-03 | 6 | TEC | promoter of Mat2a antisense radiation induced circulating long non-coding RNA [Source:MGI Symbol;Acc:MGI:1925358] |
| Slain2 | 0,3439 | 6,127E-03 | 5 | protein_coding | SLAIN motif family, member 2 [Source:MGI Symbol;Acc:MGI:1923241] |
| Shank3 | -0,1555 | 6,127E-03 | 15 | protein_coding | SH3 and multiple ankyrin repeat domains 3 [Source:MGI Symbol;Acc:MGI:1930016] |
| Ankrd55 | 0,6427 | 6,196E-03 | 13 | protein_coding | ankyrin repeat domain 55 [Source:MGI Symbol;Acc:MGI:1924568] |
| Gm37331 | 1,3191 | 6,387E-03 | 1 | TEC | predicted gene, 37331 [Source:MGI Symbol;Acc:MGI:5610559] |
| Gm26578 | 3,8263 | 6,442E-03 | 8 | lincRNA | predicted gene, 26578 [Source:MGI Symbol;Acc:MGI:5477072] |

| | | | | | |
|---|---|---|---|---|---|
| Fxn | 0,3798 | 6,447E-03 | 19 | protein_coding | frataxin [Source:MGI Symbol;Acc:MGI:1096879] |
| Gm10513 | 3,2449 | 6,483E-03 | 17 | TEC | predicted gene 10513 [Source:MGI Symbol;Acc:MGI:3641888] |
| Gm5617 | -0,5433 | 6,619E-03 | 9 | protein_coding | predicted gene 5617 [Source:MGI Symbol;Acc:MGI:3643566] |
| Olfr460 | 3,8872 | 6,629E-03 | 6 | protein_coding | olfactory receptor 460 [Source:MGI Symbol;Acc:MGI:3030294] |
| Epx | 3,6314 | 6,683E-03 | 11 | protein_coding | eosinophil peroxidase [Source:MGI Symbol;Acc:MGI:107569] |
| Pcdhga4 | 0,5718 | 6,743E-03 | 18 | protein_coding | protocadherin gamma subfamily A, 4 [Source:MGI Symbol;Acc:MGI:1935216] |
| Ddx10 | 0,3238 | 6,793E-03 | 9 | protein_coding | DEAD (Asp-Glu-Ala-Asp) box polypeptide 10 [Source:MGI Symbol;Acc:MGI:1924841] |
| Gm4631 | 2,2771 | 6,820E-03 | 2 | protein_coding | predicted gene 4631 [Source:MGI Symbol;Acc:MGI:3782813] |
| Cast | 0,3962 | 6,835E-03 | 13 | protein_coding | calpastatin [Source:MGI Symbol;Acc:MGI:1098236] |
| Lrrc32 | 0,6397 | 7,088E-03 | 7 | protein_coding | leucine rich repeat containing 32 [Source:MGI Symbol;Acc:MGI:93882] |
| Gm42585 | 1,6117 | 7,315E-03 | 5 | TEC | predicted gene 42585 [Source:MGI Symbol;Acc:MGI:5662722] |
| Mrpl28 | -0,2133 | 7,387E-03 | 17 | protein_coding | mitochondrial ribosomal protein L28 [Source:MGI Symbol;Acc:MGI:1915861] |
| Pnpt1 | 0,3286 | 7,402E-03 | 11 | protein_coding | polyribonucleotide nucleotidyltransferase 1 [Source:MGI Symbol;Acc:MGI:1918951] |
| Smarca5-ps | 3,4235 | 7,552E-03 | 4 | transcribed_processed_pseudogene | SWI/SNF related, matrix associated, actin depenent ragulator of chromatin, subfamily a, member 5, pseudogene [Source:MGI Symbol;Acc:MGI:2443330] |
| Fev | -3,8129 | 7,671E-03 | 1 | protein_coding | FEV (ETS oncogene family) [Source:MGI Symbol;Acc:MGI:2449712] |
| Gm42531 | 3,1282 | 7,736E-03 | 5 | sense_intronic | predicted gene 42531 [Source:MGI Symbol;Acc:MGI:5662668] |
| Slamf8 | -2,3451 | 7,739E-03 | 1 | protein_coding | SLAM family member 8 [Source:MGI Symbol;Acc:MGI:1921998] |
| Rps12-ps3 | -0,2619 | 7,849E-03 | 19 | protein_coding | ribosomal protein S12, pseudogene 3 [Source:MGI Symbol;Acc:MGI:3704503] |
| Gm17082 | -3,5938 | 7,882E-03 | 14 | processed_pseudogene | predicted gene 17082 [Source:MGI Symbol;Acc:MGI:4937909] |
| Mphosph8 | 0,2650 | 7,954E-03 | 14 | protein_coding | M-phase phosphoprotein 8 [Source:MGI Symbol;Acc:MGI:1922589] |
| Poln | 1,5759 | 8,003E-03 | 5 | protein_coding | DNA polymerase N [Source:MGI Symbol;Acc:MGI:2675617] |
| Acot12 | -1,4300 | 8,034E-03 | 13 | protein_coding | acyl-CoA thioesterase 12 [Source:MGI Symbol;Acc:MGI:1921406] |
| Dcakd | -0,2413 | 8,077E-03 | 11 | protein_coding | dephospho-CoA kinase domain containing [Source:MGI Symbol;Acc:MGI:1915337] |
| Zdhhc12 | -0,4258 | 8,146E-03 | 2 | protein_coding | zinc finger, DHHC domain containing 12 [Source:MGI Symbol;Acc:MGI:1913470] |
| Wdr43 | 0,2308 | 8,159E-03 | 17 | protein_coding | WD repeat domain 43 [Source:MGI Symbol;Acc:MGI:1919765] |
| Gm48449 | 3,6087 | 8,224E-03 | 14 | antisense | predicted gene, 48449 [Source:MGI Symbol;Acc:MGI:6097961] |
| Avpr1b | -1,2089 | 8,295E-03 | 1 | protein_coding | arginine vasopressin receptor 1B [Source:MGI Symbol;Acc:MGI:1347010] |

| | | | | | |
|---|---|---|---|---|---|
| Gbp4 | 0,6923 | 8,420E-03 | 5 | protein_coding | guanylate binding protein 4 [Source:MGI Symbol;Acc:MGI:97072] |
| D930016D06Rik | 0,3142 | 8,475E-03 | 5 | lincRNA | RIKEN cDNA D930016D06 gene [Source:MGI Symbol;Acc:MGI:2442700] |
| Gm32219 | 2,8406 | 8,617E-03 | 12 | lincRNA | predicted gene, 32219 [Source:MGI Symbol;Acc:MGI:5591378] |
| Gm48888 | 3,7556 | 8,621E-03 | 13 | TEC | predicted gene, 48888 [Source:MGI Symbol;Acc:MGI:6098648] |
| Gm5805 | -0,4093 | 8,636E-03 | 15 | processed_pseudogene | predicted gene 5805 [Source:MGI Symbol;Acc:MGI:3644633] |
| Gm26894 | -1,3602 | 8,709E-03 | 5 | lincRNA | predicted gene, 26894 [Source:MGI Symbol;Acc:MGI:5477388] |
| Ankrd13b | -0,2082 | 8,749E-03 | 11 | protein_coding | ankyrin repeat domain 13b [Source:MGI Symbol;Acc:MGI:2144501] |
| Ammecr1l | 0,2309 | 8,816E-03 | 18 | protein_coding | AMME chromosomal region gene 1-like [Source:MGI Symbol;Acc:MGI:2442711] |
| Hba-a1 | -0,5159 | 8,866E-03 | 11 | protein_coding | hemoglobin alpha, adult chain 1 [Source:MGI Symbol;Acc:MGI:96015] |
| Slc46a1 | -0,3420 | 8,895E-03 | 11 | protein_coding | solute carrier family 46, member 1 [Source:MGI Symbol;Acc:MGI:1098733] |
| Tsta3 | -0,2576 | 9,154E-03 | 15 | protein_coding | tissue specific transplantation antigen P35B [Source:MGI Symbol;Acc:MGI:98857] |
| Flt4 | -0,4464 | 9,201E-03 | 11 | protein_coding | FMS-like tyrosine kinase 4 [Source:MGI Symbol;Acc:MGI:95561] |
| Gm45494 | 1,8638 | 9,281E-03 | 9 | bidirectional_promoter_lncRNA | predicted gene 45494 [Source:MGI Symbol;Acc:MGI:5791330] |
| Gm9826 | -1,1923 | 9,338E-03 | 1 | processed_pseudogene | predicted gene 9826 [Source:MGI Symbol;Acc:MGI:3642725] |
| Gm47652 | -3,6001 | 9,418E-03 | 9 | TEC | predicted gene, 47652 [Source:MGI Symbol;Acc:MGI:6096735] |
| Klrd1 | 3,5458 | 9,427E-03 | 6 | protein_coding | killer cell lectin-like receptor, subfamily D, member 1 [Source:MGI Symbol;Acc:MGI:1196275] |
| Pop5 | -0,1870 | 9,444E-03 | 5 | protein_coding | processing of precursor 5, ribonuclease P/MRP family (S, cerevisiae) [Source:MGI Symbol;Acc:MGI:2151221] |
| Klk12 | -3,6387 | 9,512E-03 | 7 | protein_coding | kallikrein related-peptidase 12 [Source:MGI Symbol;Acc:MGI:1916761] |
| Spdl1 | -1,6255 | 9,592E-03 | 11 | protein_coding | spindle apparatus coiled-coil protein 1 [Source:MGI Symbol;Acc:MGI:1917635] |
| Clk3 | -0,1988 | 9,708E-03 | 9 | protein_coding | CDC-like kinase 3 [Source:MGI Symbol;Acc:MGI:1098670] |
| Cacnb3 | -0,1681 | 9,732E-03 | 15 | protein_coding | calcium channel, voltage-dependent, beta 3 subunit [Source:MGI Symbol;Acc:MGI:103307] |
| D3Ertd751e | 0,3864 | 9,736E-03 | 3 | protein_coding | DNA segment, Chr 3, ERATO Doi 751, expressed [Source:MGI Symbol;Acc:MGI:1289213] |
| Atp13a2 | -0,1561 | 9,754E-03 | 4 | protein_coding | ATPase type 13A2 [Source:MGI Symbol;Acc:MGI:1922022] |
| Rcvrn | -1,5937 | 9,764E-03 | 11 | protein_coding | recoverin [Source:MGI Symbol;Acc:MGI:97883] |
| Tmem169 | 0,3176 | 9,816E-03 | 1 | protein_coding | transmembrane protein 169 [Source:MGI Symbol;Acc:MGI:2442781] |
| Agbl3 | 0,3975 | 9,836E-03 | 6 | protein_coding | ATP/GTP binding protein-like 3 [Source:MGI Symbol;Acc:MGI:1923473] |
| Rpl38-ps2 | -0,4092 | 9,935E-03 | 6 | processed_pseudogene | ribosomal protein L38, pseudogene 2 [Source:MGI Symbol;Acc:MGI:3646625] |

| | | | | | |
|---|---|---|---|---|---|
| Gm15779 | -4,0590 | 1,003E-02 | 6 | processed_pseudogene | predicted gene 15779 [Source:MGI Symbol;Acc:MGI:3783221] |
| mt-Co1 | 0,1682 | 1,004E-02 | MT | protein_coding | mitochondrially encoded cytochrome c oxidase I [Source:MGI Symbol;Acc:MGI:102504] |
| Lcorl | 0,4201 | 1,027E-02 | 5 | protein_coding | ligand dependent nuclear receptor corepressor-like [Source:MGI Symbol;Acc:MGI:2651932] |
| Gm17747 | 1,5012 | 1,027E-02 | 14 | lincRNA | predicted gene, 17747 [Source:MGI Symbol;Acc:MGI:5009825] |
| Mettl17 | -0,2423 | 1,041E-02 | 14 | protein_coding | methyltransferase like 17 [Source:MGI Symbol;Acc:MGI:1098577] |
| Gm7347 | 3,9703 | 1,043E-02 | 5 | protein_coding | predicted gene 7347 [Source:MGI Symbol;Acc:MGI:3645983] |
| Cwc27 | 0,3817 | 1,051E-02 | 13 | protein_coding | CWC27 spliceosome-associated protein [Source:MGI Symbol;Acc:MGI:1914535] |
| Dcdc5 | -1,5331 | 1,067E-02 | 2 | transcribed_unitary_pseudogene | doublecortin domain containing 5 [Source:MGI Symbol;Acc:MGI:3045363] |
| Rpl21 | -0,3046 | 1,070E-02 | 5 | protein_coding | ribosomal protein L21 [Source:MGI Symbol;Acc:MGI:1278340] |
| Tnk2os | 2,8041 | 1,079E-02 | 16 | antisense | tyrosine kinase, non-receptor 2, opposite strand [Source:MGI Symbol;Acc:MGI:3642140] |
| Vps37d | -0,2109 | 1,090E-02 | 5 | protein_coding | vacuolar protein sorting 37D [Source:MGI Symbol;Acc:MGI:2159402] |
| Gm37470 | 3,3629 | 1,095E-02 | 1 | TEC | predicted gene, 37470 [Source:MGI Symbol;Acc:MGI:5610698] |
| Tom1l2 | -0,1324 | 1,098E-02 | 11 | protein_coding | target of myb1-like 2 (chicken) [Source:MGI Symbol;Acc:MGI:2443306] |
| Camsap3 | -0,2278 | 1,114E-02 | 8 | protein_coding | calmodulin regulated spectrin-associated protein family, member 3 [Source:MGI Symbol;Acc:MGI:1916947] |
| Mir3080 | 3,9412 | 1,136E-02 | 15 | miRNA | microRNA 3080 [Source:MGI Symbol;Acc:MGI:4834253] |
| Pcdhga10 | 2,8335 | 1,143E-02 | 18 | protein_coding | protocadherin gamma subfamily A, 10 [Source:MGI Symbol;Acc:MGI:1935227] |
| Ddx46 | 0,1688 | 1,150E-02 | 13 | protein_coding | DEAD (Asp-Glu-Ala-Asp) box polypeptide 46 [Source:MGI Symbol;Acc:MGI:1920895] |
| Tmem254c | 0,9685 | 1,159E-02 | 14 | protein_coding | transmembrane protein 254c [Source:MGI Symbol;Acc:MGI:3711260] |
| Hmgcll1 | 0,3220 | 1,163E-02 | 9 | protein_coding | 3-hydroxymethyl-3-methylglutaryl-Coenzyme A lyase-like 1 [Source:MGI Symbol;Acc:MGI:2446108] |
| Pdgfd | 0,5371 | 1,171E-02 | 9 | protein_coding | platelet-derived growth factor, D polypeptide [Source:MGI Symbol;Acc:MGI:1919035] |
| Gm43909 | 0,8132 | 1,174E-02 | 6 | TEC | predicted gene, 43909 [Source:MGI Symbol;Acc:MGI:5690301] |
| Zswim7 | -0,3680 | 1,176E-02 | 11 | protein_coding | zinc finger SWIM-type containing 7 [Source:MGI Symbol;Acc:MGI:1916997] |
| Man2a1 | 0,2711 | 1,178E-02 | 17 | protein_coding | mannosidase 2, alpha 1 [Source:MGI Symbol;Acc:MGI:104669] |
| Gm37120 | 1,2847 | 1,187E-02 | 1 | TEC | predicted gene, 37120 [Source:MGI Symbol;Acc:MGI:5610348] |
| 4833421K07Rik | 4,1896 | 1,191E-02 | 7 | TEC | RIKEN cDNA 4833421K07 gene [Source:MGI Symbol;Acc:MGI:1921844] |
| Plekhg4 | -0,6980 | 1,196E-02 | 8 | protein_coding | pleckstrin homology domain containing, family G (with RhoGef domain) member 4 [Source:MGI Symbol;Acc:MGI:2142544] |
| Fgl2 | 0,6793 | 1,203E-02 | 5 | protein_coding | fibrinogen-like protein 2 [Source:MGI Symbol;Acc:MGI:103266] |

| | | | | | |
|---|---|---|---|---|---|
| Gm7457 | 1,9986 | 1,207E-02 | 6 | processed_transcript | predicted gene 7457 [Source:MGI Symbol;Acc:MGI:3648132] |
| Gm23713 | 2,9903 | 1,212E-02 | 13 | snoRNA | predicted gene, 23713 [Source:MGI Symbol;Acc:MGI:5453490] |
| Sac3d1 | -0,2776 | 1,213E-02 | 19 | protein_coding | SAC3 domain containing 1 [Source:MGI Symbol;Acc:MGI:1913656] |
| Gm13029 | 4,1476 | 1,214E-02 | 4 | antisense | predicted gene 13029 [Source:MGI Symbol;Acc:MGI:3651764] |
| Bcr | -0,1208 | 1,217E-02 | 10 | protein_coding | breakpoint cluster region [Source:MGI Symbol;Acc:MGI:88141] |
| Adam21 | 0,8802 | 1,224E-02 | 12 | protein_coding | a disintegrin and metallopeptidase domain 21 [Source:MGI Symbol;Acc:MGI:1861229] |
| A330093E20Rik | 0,5826 | 1,227E-02 | 18 | antisense | RIKEN cDNA A330093E20 gene [Source:MGI Symbol;Acc:MGI:2147325] |
| Gm34821 | 3,3169 | 1,233E-02 | 7 | antisense | predicted gene, 34821 [Source:MGI Symbol;Acc:MGI:5593980] |
| 2310040G07Rik | 2,1075 | 1,249E-02 | 5 | antisense | RIKEN cDNA 2310040G07 gene [Source:MGI Symbol;Acc:MGI:1917534] |
| Magix | 1,4390 | 1,254E-02 | X | protein_coding | MAGI family member, X-linked [Source:MGI Symbol;Acc:MGI:1859644] |
| Ptprr | 0,2734 | 1,254E-02 | 10 | protein_coding | protein tyrosine phosphatase, receptor type, R [Source:MGI Symbol;Acc:MGI:109559] |
| Gm12966 | 0,4505 | 1,267E-02 | 4 | processed_pseudogene | predicted gene 12966 [Source:MGI Symbol;Acc:MGI:3652043] |
| Epb42 | -4,1685 | 1,277E-02 | 2 | protein_coding | erythrocyte membrane protein band 4,2 [Source:MGI Symbol;Acc:MGI:95402] |
| Fuk | -0,2301 | 1,279E-02 | 8 | protein_coding | fucokinase [Source:MGI Symbol;Acc:MGI:1916071] |
| Lax1 | 2,2261 | 1,281E-02 | 1 | protein_coding | lymphocyte transmembrane adaptor 1 [Source:MGI Symbol;Acc:MGI:2443362] |
| Slco1a4 | 0,2446 | 1,288E-02 | 6 | protein_coding | solute carrier organic anion transporter family, member 1a4 [Source:MGI Symbol;Acc:MGI:1351896] |
| Gm16835 | 0,5226 | 1,294E-02 | 14 | processed_transcript | predicted gene, 16835 [Source:MGI Symbol;Acc:MGI:4439759] |
| Panx2 | -0,1853 | 1,305E-02 | 15 | protein_coding | pannexin 2 [Source:MGI Symbol;Acc:MGI:1890615] |
| Mdm1 | 0,3284 | 1,305E-02 | 10 | protein_coding | transformed mouse 3T3 cell double minute 1 [Source:MGI Symbol;Acc:MGI:96951] |
| Hsp90aa1 | 0,1572 | 1,306E-02 | 12 | protein_coding | heat shock protein 90, alpha (cytosolic), class A member 1 [Source:MGI Symbol;Acc:MGI:96250] |
| Plin4 | -0,5147 | 1,324E-02 | 17 | protein_coding | perilipin 4 [Source:MGI Symbol;Acc:MGI:1929709] |
| Zfp758 | 0,3781 | 1,327E-02 | 17 | protein_coding | zinc finger protein 758 [Source:MGI Symbol;Acc:MGI:2385044] |
| Gm2464 | -0,6221 | 1,330E-02 | 3 | lincRNA | predicted gene 2464 [Source:MGI Symbol;Acc:MGI:3780631] |
| Uck1 | -0,1628 | 1,331E-02 | 2 | protein_coding | uridine-cytidine kinase 1 [Source:MGI Symbol;Acc:MGI:98904] |
| Gm14092 | 4,1453 | 1,332E-02 | 2 | antisense | predicted gene 14092 [Source:MGI Symbol;Acc:MGI:3651965] |
| AC154912,1 | 4,2251 | 1,338E-02 | 17 | antisense | novel transcript, antisense to Fkbp5 |
| Pnrc1 | 0,1883 | 1,349E-02 | 4 | protein_coding | proline-rich nuclear receptor coactivator 1 [Source:MGI Symbol;Acc:MGI:1917838] |

| | | | | | |
|---|---|---|---|---|---|
| Emp3 | 0,6876 | 1,356E-02 | 7 | protein_coding | epithelial membrane protein 3 [Source:MGI Symbol;Acc:MGI:1098729] |
| Phldb2 | 0,4371 | 1,357E-02 | 16 | protein_coding | pleckstrin homology like domain, family B, member 2 [Source:MGI Symbol;Acc:MGI:2444981] |
| Stoml1 | -0,1855 | 1,362E-02 | 9 | protein_coding | stomatin-like 1 [Source:MGI Symbol;Acc:MGI:1916356] |
| Inca1 | -0,5533 | 1,364E-02 | 11 | protein_coding | inhibitor of CDK, cyclin A1 interacting protein 1 [Source:MGI Symbol;Acc:MGI:2144284] |
| Gm29668 | -3,7474 | 1,379E-02 | 1 | processed_pseudogene | predicted gene 29668 [Source:MGI Symbol;Acc:MGI:5580374] |
| E130006D01Rik | -1,2674 | 1,403E-02 | 5 | lincRNA | RIKEN cDNA E130006D01 gene [Source:MGI Symbol;Acc:MGI:2685527] |
| Zfp575 | -0,3033 | 1,409E-02 | 7 | protein_coding | zinc finger protein 575 [Source:MGI Symbol;Acc:MGI:2141921] |
| Gm6517 | 3,5206 | 1,411E-02 | 5 | processed_pseudogene | predicted gene 6517 [Source:MGI Symbol;Acc:MGI:3643656] |
| Uba6 | 0,3243 | 1,411E-02 | 5 | protein_coding | ubiquitin-like modifier activating enzyme 6 [Source:MGI Symbol;Acc:MGI:1913894] |
| Ide | 0,3246 | 1,419E-02 | 19 | protein_coding | insulin degrading enzyme [Source:MGI Symbol;Acc:MGI:96412] |
| Gatd1 | -0,2212 | 1,423E-02 | 7 | protein_coding | glutamine amidotransferase like class 1 domain containing 1 [Source:MGI Symbol;Acc:MGI:2387178] |
| Ube2g2 | -0,1867 | 1,430E-02 | 10 | protein_coding | ubiquitin-conjugating enzyme E2G 2 [Source:MGI Symbol;Acc:MGI:1343188] |
| Prkag2os2 | -0,5975 | 1,432E-02 | 5 | antisense | protein kinase, AMP-activated, gamma 2 non-catalytic subunit, opposite strand 2 [Source:MGI Symbol;Acc:MGI:3783001] |
| Gtf3c1 | -0,1391 | 1,441E-02 | 7 | protein_coding | general transcription factor III C 1 [Source:MGI Symbol;Acc:MGI:107887] |
| Zfp850 | 0,3790 | 1,473E-02 | 7 | protein_coding | zinc finger protein 850 [Source:MGI Symbol;Acc:MGI:3036281] |
| Prrx1 | 0,3532 | 1,492E-02 | 1 | protein_coding | paired related homeobox 1 [Source:MGI Symbol;Acc:MGI:97712] |
| Gm37986 | 0,9553 | 1,496E-02 | 1 | lincRNA | predicted gene, 37986 [Source:MGI Symbol;Acc:MGI:5611214] |
| Eif6 | -0,2112 | 1,510E-02 | 2 | protein_coding | eukaryotic translation initiation factor 6 [Source:MGI Symbol;Acc:MGI:1196288] |
| Lrrc10b | -0,2482 | 1,517E-02 | 19 | protein_coding | leucine rich repeat containing 10B [Source:MGI Symbol;Acc:MGI:2685551] |
| 1700034G24Rik | 1,4163 | 1,521E-02 | 5 | lincRNA | RIKEN cDNA 1700034G24 gene [Source:MGI Symbol;Acc:MGI:1914575] |
| Vash1 | -0,2338 | 1,525E-02 | 12 | protein_coding | vasohibin 1 [Source:MGI Symbol;Acc:MGI:2442543] |
| B3glct | 0,2154 | 1,529E-02 | 5 | protein_coding | beta-3-glucosyltransferase [Source:MGI Symbol;Acc:MGI:2685903] |
| Tbc1d22a | -0,1749 | 1,533E-02 | 15 | protein_coding | TBC1 domain family, member 22a [Source:MGI Symbol;Acc:MGI:1289265] |
| Pla2g15 | -0,1972 | 1,536E-02 | 8 | protein_coding | phospholipase A2, group XV [Source:MGI Symbol;Acc:MGI:2178076] |
| mt-Co2 | 0,2389 | 1,549E-02 | MT | protein_coding | mitochondrially encoded cytochrome c oxidase II [Source:MGI Symbol;Acc:MGI:102503] |
| Gm43034 | -0,5568 | 1,551E-02 | 5 | TEC | predicted gene 43034 [Source:MGI Symbol;Acc:MGI:5663171] |
| Qser1 | 0,2719 | 1,556E-02 | 2 | protein_coding | glutamine and serine rich 1 [Source:MGI Symbol;Acc:MGI:2138986] |

| | | | | | |
|---|---|---|---|---|---|
| Gm11906 | -3,9202 | 1,570E-02 | 4 | processed_transcript | predicted gene 11906 [Source:MGI Symbol;Acc:MGI:3705114] |
| Gm26582 | -1,3311 | 1,585E-02 | 5 | lincRNA | predicted gene, 26582 [Source:MGI Symbol;Acc:MGI:5477076] |
| Scap | -0,1636 | 1,588E-02 | 9 | protein_coding | SREBF chaperone [Source:MGI Symbol;Acc:MGI:2135958] |
| Zfp300 | 0,6494 | 1,591E-02 | X | protein_coding | zinc finger protein 300 [Source:MGI Symbol;Acc:MGI:3045326] |
| Gm3257 | -1,8055 | 1,601E-02 | 7 | processed_pseudogene | predicted gene 3257 [Source:MGI Symbol;Acc:MGI:3781435] |
| Atp6ap2 | 0,1690 | 1,606E-02 | X | protein_coding | ATPase, H+ transporting, lysosomal accessory protein 2 [Source:MGI Symbol;Acc:MGI:1917745] |
| Gm18637 | -3,9485 | 1,610E-02 | 10 | processed_pseudogene | predicted gene, 18637 [Source:MGI Symbol;Acc:MGI:5010822] |
| Zfp975 | 0,4893 | 1,628E-02 | 7 | protein_coding | zinc finger protein 975 [Source:MGI Symbol;Acc:MGI:3648690] |
| Ffar4 | 3,4925 | 1,629E-02 | 19 | protein_coding | free fatty acid receptor 4 [Source:MGI Symbol;Acc:MGI:2147577] |
| Tsc2 | -0,1522 | 1,631E-02 | 17 | protein_coding | tuberous sclerosis 2 [Source:MGI Symbol;Acc:MGI:102548] |
| Gm46515 | -2,8230 | 1,648E-02 | 15 | lincRNA | predicted gene, 46515 [Source:MGI Symbol;Acc:MGI:5826152] |
| Ltbp1 | 0,4121 | 1,650E-02 | 17 | protein_coding | latent transforming growth factor beta binding protein 1 [Source:MGI Symbol;Acc:MGI:109151] |
| Gm21093 | 0,7613 | 1,661E-02 | 4 | protein_coding | predicted gene, 21093 [Source:MGI Symbol;Acc:MGI:5434448] |
| Shisal2a | 3,0848 | 1,669E-02 | 4 | protein_coding | shisa like 2A [Source:MGI Symbol;Acc:MGI:3651644] |
| Gm45772 | -3,4160 | 1,670E-02 | 8 | processed_pseudogene | predicted gene 45772 [Source:MGI Symbol;Acc:MGI:5804887] |
| Rps2-ps13 | -0,3076 | 1,677E-02 | X | processed_pseudogene | ribosomal protein S2, pseudogene 13 [Source:MGI Symbol;Acc:MGI:3705640] |
| Cys1 | -0,2837 | 1,678E-02 | 12 | protein_coding | cystin 1 [Source:MGI Symbol;Acc:MGI:2177632] |
| Tdrd5 | -0,6532 | 1,679E-02 | 1 | protein_coding | tudor domain containing 5 [Source:MGI Symbol;Acc:MGI:2684949] |
| Wdtc1 | -0,1611 | 1,686E-02 | 4 | protein_coding | WD and tetratricopeptide repeats 1 [Source:MGI Symbol;Acc:MGI:2685541] |
| Gm15082 | 3,2337 | 1,701E-02 | 6 | antisense | predicted gene 15082 [Source:MGI Symbol;Acc:MGI:3705124] |
| Gkap1 | 0,4036 | 1,713E-02 | 13 | protein_coding | G kinase anchoring protein 1 [Source:MGI Symbol;Acc:MGI:1891694] |
| 4921523L03Rik | -3,7806 | 1,721E-02 | 9 | antisense | RIKEN cDNA 4921523L03 gene [Source:MGI Symbol;Acc:MGI:1918163] |
| Ppp1r12c | -0,1519 | 1,722E-02 | 7 | protein_coding | protein phosphatase 1, regulatory subunit 12C [Source:MGI Symbol;Acc:MGI:1924258] |
| Gm44644 | 0,5850 | 1,736E-02 | 7 | lincRNA | predicted gene 44644 [Source:MGI Symbol;Acc:MGI:5753220] |
| Gm43923 | 2,9923 | 1,745E-02 | 6 | TEC | predicted gene, 43923 [Source:MGI Symbol;Acc:MGI:5690315] |
| Itga11 | -0,4021 | 1,745E-02 | 9 | protein_coding | integrin alpha 11 [Source:MGI Symbol;Acc:MGI:2442114] |
| Gm12413 | -3,5495 | 1,750E-02 | 4 | processed_pseudogene | predicted gene 12413 [Source:MGI Symbol;Acc:MGI:3650373] |

| | | | | | |
|---|---|---|---|---|---|
| Lrrc9 | 0,9313 | 1,752E-02 | 12 | protein_coding | leucine rich repeat containing 9 [Source:MGI Symbol;Acc:MGI:1925507] |
| Gm38273 | 1,6433 | 1,752E-02 | 2 | TEC | predicted gene, 38273 [Source:MGI Symbol;Acc:MGI:5611501] |
| Mvb12a | -0,2436 | 1,752E-02 | 8 | protein_coding | multivesicular body subunit 12A [Source:MGI Symbol;Acc:MGI:1920961] |
| Dnajb4 | 0,1799 | 1,760E-02 | 3 | protein_coding | DnaJ heat shock protein family (Hsp40) member B4 [Source:MGI Symbol;Acc:MGI:1914285] |
| Gm45516 | 2,3793 | 1,766E-02 | 8 | TEC | predicted gene 45516 [Source:MGI Symbol;Acc:MGI:5791352] |
| 5033417F24Rik | -0,5016 | 1,778E-02 | 1 | antisense | RIKEN cDNA 5033417F24 gene [Source:MGI Symbol;Acc:MGI:1923245] |
| Slc6a15 | 0,2000 | 1,780E-02 | 10 | protein_coding | solute carrier family 6 (neurotransmitter transporter), member 15 [Source:MGI Symbol;Acc:MGI:2143484] |
| Trip11 | 0,1843 | 1,797E-02 | 12 | protein_coding | thyroid hormone receptor interactor 11 [Source:MGI Symbol;Acc:MGI:1924393] |
| Slc35c2 | -0,1768 | 1,798E-02 | 2 | protein_coding | solute carrier family 35, member C2 [Source:MGI Symbol;Acc:MGI:2385166] |
| Vcan | 0,2487 | 1,805E-02 | 13 | protein_coding | versican [Source:MGI Symbol;Acc:MGI:102889] |
| Cpne9 | -0,2672 | 1,805E-02 | 6 | protein_coding | copine family member IX [Source:MGI Symbol;Acc:MGI:2443052] |
| Ppp1r37 | -0,1505 | 1,811E-02 | 7 | protein_coding | protein phosphatase 1, regulatory subunit 37 [Source:MGI Symbol;Acc:MGI:2687042] |
| Gpr139 | -0,7749 | 1,815E-02 | 7 | protein_coding | G protein-coupled receptor 139 [Source:MGI Symbol;Acc:MGI:2685341] |
| Chchd1 | -0,2088 | 1,815E-02 | 14 | protein_coding | coiled-coil-helix-coiled-coil-helix domain containing 1 [Source:MGI Symbol;Acc:MGI:1913371] |
| Eea1 | 0,2808 | 1,825E-02 | 10 | protein_coding | early endosome antigen 1 [Source:MGI Symbol;Acc:MGI:2442192] |
| Rfc5 | -0,2275 | 1,829E-02 | 5 | protein_coding | replication factor C (activator 1) 5 [Source:MGI Symbol;Acc:MGI:1919401] |
| Gm36500 | -1,3194 | 1,836E-02 | 13 | lincRNA | predicted gene, 36500 [Source:MGI Symbol;Acc:MGI:5595659] |
| AC131800,1 | -1,2233 | 1,845E-02 | 17 | bidirectional_promoter_lncRNA | novel transcript |
| Gm4853 | -3,0377 | 1,857E-02 | X | processed_pseudogene | predicted pseudogene 4853 [Source:MGI Symbol;Acc:MGI:3644655] |
| Dlx6 | 0,5320 | 1,872E-02 | 6 | protein_coding | distal-less homeobox 6 [Source:MGI Symbol;Acc:MGI:101927] |
| C230086J09Rik | 0,9009 | 1,873E-02 | 14 | TEC | RIKEN cDNA C230086J09 gene [Source:MGI Symbol;Acc:MGI:2443423] |
| Tfrc | 0,2833 | 1,881E-02 | 16 | protein_coding | transferrin receptor [Source:MGI Symbol;Acc:MGI:98822] |
| Mir32 | -3,9070 | 1,882E-02 | 4 | miRNA | microRNA 32 [Source:MGI Symbol;Acc:MGI:3619331] |
| Gm36028 | -4,0697 | 1,884E-02 | 16 | protein_coding | predicted gene, 36028 [Source:MGI Symbol;Acc:MGI:5595187] |
| Daam2 | -0,1831 | 1,890E-02 | 17 | protein_coding | dishevelled associated activator of morphogenesis 2 [Source:MGI Symbol;Acc:MGI:1923691] |
| Gm14434 | -3,0289 | 1,891E-02 | 2 | protein_coding | predicted gene 14434 [Source:MGI Symbol;Acc:MGI:3702417] |
| Tmem151b | -0,1638 | 1,892E-02 | 17 | protein_coding | transmembrane protein 151B [Source:MGI Symbol;Acc:MGI:2685169] |

| | | | | | |
|---|---|---|---|---|---|
| Melk | 3,7313 | 1,894E-02 | 4 | protein_coding | maternal embryonic leucine zipper kinase [Source:MGI Symbol;Acc:MGI:106924] |
| Cd19 | -3,3622 | 1,905E-02 | 7 | protein_coding | CD19 antigen [Source:MGI Symbol;Acc:MGI:88319] |
| Gm18225 | 3,5669 | 1,916E-02 | 9 | unprocessed_pseudogene | predicted gene, 18225 [Source:MGI Symbol;Acc:MGI:5010410] |
| Clec2l | -0,2267 | 1,918E-02 | 6 | protein_coding | C-type lectin domain family 2, member L [Source:MGI Symbol;Acc:MGI:2141402] |
| Mterf1b | 0,6514 | 1,926E-02 | 5 | protein_coding | mitochondrial transcription termination factor 1b [Source:MGI Symbol;Acc:MGI:3704243] |
| Pcmtd1 | 0,2246 | 1,930E-02 | 1 | protein_coding | protein-L-isoaspartate (D-aspartate) O-methyltransferase domain containing 1 [Source:MGI Symbol;Acc:MGI:2441773] |
| Gm38366 | -1,5770 | 1,930E-02 | 9 | TEC | predicted gene, 38366 [Source:MGI Symbol;Acc:MGI:5611594] |
| Zfp182 | 0,2820 | 1,938E-02 | X | protein_coding | zinc finger protein 182 [Source:MGI Symbol;Acc:MGI:2442220] |
| D230017M19Rik | -0,4151 | 1,948E-02 | 1 | lincRNA | RIKEN cDNA D230017M19 gene [Source:MGI Symbol;Acc:MGI:2445071] |
| Arhgap42 | 0,2792 | 1,950E-02 | 9 | protein_coding | Rho GTPase activating protein 42 [Source:MGI Symbol;Acc:MGI:1918794] |
| AC078895,1 | -0,4329 | 1,966E-02 | 16 | lincRNA | novel transcript |
| Tbx21 | -3,1761 | 1,973E-02 | 11 | protein_coding | T-box 21 [Source:MGI Symbol;Acc:MGI:1888984] |
| Nf2 | -0,1375 | 2,007E-02 | 11 | protein_coding | neurofibromin 2 [Source:MGI Symbol;Acc:MGI:97307] |
| Gm5422 | 0,7272 | 2,008E-02 | 10 | transcribed_processed_pseudogene | predicted pseudogene 5422 [Source:MGI Symbol;Acc:MGI:3643411] |
| Mc5r | 2,3195 | 2,009E-02 | 18 | protein_coding | melanocortin 5 receptor [Source:MGI Symbol;Acc:MGI:99420] |
| Snai3 | 2,7010 | 2,009E-02 | 8 | protein_coding | snail family zinc finger 3 [Source:MGI Symbol;Acc:MGI:1353563] |
| 4930461C15Rik | 3,4648 | 2,012E-02 | 16 | lincRNA | RIKEN cDNA 4930461C15 gene [Source:MGI Symbol;Acc:MGI:1922133] |
| Zfp867 | -0,3773 | 2,013E-02 | 11 | protein_coding | zinc finger protein 867 [Source:MGI Symbol;Acc:MGI:2681848] |
| Gm42470 | 3,6529 | 2,019E-02 | 9 | lincRNA | predicted gene 42470 [Source:MGI Symbol;Acc:MGI:5662607] |
| Klk1b5 | 4,0657 | 2,020E-02 | 7 | protein_coding | kallikrein 1-related peptidase b5 [Source:MGI Symbol;Acc:MGI:892020] |
| Gm10686 | 1,2717 | 2,020E-02 | 9 | TEC | predicted gene 10686 [Source:MGI Symbol;Acc:MGI:3704331] |
| Fam32a | -0,1479 | 2,020E-02 | 8 | protein_coding | family with sequence similarity 32, member A [Source:MGI Symbol;Acc:MGI:1915172] |
| Aspscr1 | -0,1868 | 2,024E-02 | 11 | protein_coding | alveolar soft part sarcoma chromosome region, candidate 1 (human) [Source:MGI Symbol;Acc:MGI:1916188] |
| Kctd9 | 0,2822 | 2,028E-02 | 14 | protein_coding | potassium channel tetramerisation domain containing 9 [Source:MGI Symbol;Acc:MGI:2145579] |
| Tbc1d14 | -0,1594 | 2,039E-02 | 5 | protein_coding | TBC1 domain family, member 14 [Source:MGI Symbol;Acc:MGI:1098708] |
| Gm45209 | -3,5013 | 2,050E-02 | 7 | processed_pseudogene | predicted gene 45209 [Source:MGI Symbol;Acc:MGI:5753785] |
| Gm4974 | -3,6021 | 2,051E-02 | 7 | processed_pseudogene | predicted gene 4974 [Source:MGI Symbol;Acc:MGI:3646627] |

| | | | | | |
|---|---|---|---|---|---|
| Acadl | 0,2402 | 2,057E-02 | 1 | protein_coding | acyl-Coenzyme A dehydrogenase, long-chain [Source:MGI Symbol;Acc:MGI:87866] |
| Il13ra1 | 0,3655 | 2,061E-02 | X | protein_coding | interleukin 13 receptor, alpha 1 [Source:MGI Symbol;Acc:MGI:105052] |
| Asphd1 | -0,2346 | 2,073E-02 | 7 | protein_coding | aspartate beta-hydroxylase domain containing 1 [Source:MGI Symbol;Acc:MGI:2685014] |
| Ppp1r16b | -0,1596 | 2,076E-02 | 2 | protein_coding | protein phosphatase 1, regulatory subunit 16B [Source:MGI Symbol;Acc:MGI:2151841] |
| 4933432I03Rik | -1,7097 | 2,080E-02 | 14 | lincRNA | RIKEN cDNA 4933432I03 gene [Source:MGI Symbol;Acc:MGI:1918514] |
| Gm43534 | 3,6721 | 2,084E-02 | 3 | TEC | predicted gene 43534 [Source:MGI Symbol;Acc:MGI:5663671] |
| Gm44866 | -0,9068 | 2,086E-02 | 7 | lincRNA | predicted gene 44866 [Source:MGI Symbol;Acc:MGI:5753442] |
| Gm10925 | 0,4659 | 2,100E-02 | 1 | unprocessed_pseudogene | predicted gene 10925 [Source:MGI Symbol;Acc:MGI:3809095] |
| Kcnip2 | -0,1306 | 2,108E-02 | 19 | protein_coding | Kv channel-interacting protein 2 [Source:MGI Symbol;Acc:MGI:2135916] |
| Ccpg1 | 0,8763 | 2,113E-02 | 9 | protein_coding | cell cycle progression 1 [Source:MGI Symbol;Acc:MGI:1196419] |
| Klhl26 | -0,1389 | 2,120E-02 | 8 | protein_coding | kelch-like 26 [Source:MGI Symbol;Acc:MGI:2443079] |
| C230057M02Rik | -0,2269 | 2,125E-02 | 8 | lincRNA | RIKEN cDNA C230057M02 gene [Source:MGI Symbol;Acc:MGI:2442079] |
| Gm40055 | 3,4799 | 2,126E-02 | 3 | lincRNA | predicted gene, 40055 [Source:MGI Symbol;Acc:MGI:5622940] |
| Hspa13 | 0,2438 | 2,128E-02 | 16 | protein_coding | heat shock protein 70 family, member 13 [Source:MGI Symbol;Acc:MGI:1309463] |
| Il11ra2 | -4,2723 | 2,131E-02 | JH584293,1 | protein_coding | interleukin-11 receptor subunit alpha-2-like [Source:NCBI gene;Acc:100861969] |
| Lin37 | -0,2361 | 2,142E-02 | 7 | protein_coding | lin-37 homolog (C, elegans) [Source:MGI Symbol;Acc:MGI:1922910] |
| Gm45605 | -0,3881 | 2,146E-02 | 11 | lincRNA | predicted gene 45605 [Source:MGI Symbol;Acc:MGI:5791441] |
| Gm44586 | -2,0415 | 2,148E-02 | 7 | TEC | predicted gene 44586 [Source:MGI Symbol;Acc:MGI:5753162] |
| 1700123O20Rik | -0,2144 | 2,151E-02 | 14 | protein_coding | RIKEN cDNA 1700123O20 gene [Source:MGI Symbol;Acc:MGI:1920893] |
| 5430427N15Rik | -1,8197 | 2,158E-02 | 5 | lincRNA | RIKEN cDNA 5430427N15 gene [Source:MGI Symbol;Acc:MGI:2441699] |
| Gm37402 | 3,1758 | 2,159E-02 | 17 | TEC | predicted gene, 37402 [Source:MGI Symbol;Acc:MGI:5610630] |
| Grhpr | -0,1866 | 2,169E-02 | 4 | protein_coding | glyoxylate reductase/hydroxypyruvate reductase [Source:MGI Symbol;Acc:MGI:1923488] |
| Nudt14 | -0,3171 | 2,170E-02 | 12 | protein_coding | nudix (nucleoside diphosphate linked moiety X)-type motif 14 [Source:MGI Symbol;Acc:MGI:1913424] |
| Gm26761 | 3,4502 | 2,174E-02 | 5 | lincRNA | predicted gene, 26761 [Source:MGI Symbol;Acc:MGI:5477255] |
| Gm47798 | 0,8558 | 2,176E-02 | 14 | sense_intronic | predicted gene, 47798 [Source:MGI Symbol;Acc:MGI:6096974] |
| Nxt1 | -0,2861 | 2,183E-02 | 2 | protein_coding | NTF2-related export protein 1 [Source:MGI Symbol;Acc:MGI:1929619] |
| Hmgb3 | 0,2964 | 2,185E-02 | X | protein_coding | high mobility group box 3 [Source:MGI Symbol;Acc:MGI:1098219] |

| | | | | | |
|---|---|---|---|---|---|
| Tpmt | -0,3839 | 2,201E-02 | 13 | protein_coding | thiopurine methyltransferase [Source:MGI Symbol;Acc:MGI:98812] |
| Filip1l | 0,4355 | 2,214E-02 | 16 | protein_coding | filamin A interacting protein 1-like [Source:MGI Symbol;Acc:MGI:1925999] |
| Use1 | -0,1937 | 2,221E-02 | 8 | protein_coding | unconventional SNARE in the ER 1 homolog (S, cerevisiae) [Source:MGI Symbol;Acc:MGI:1914273] |
| AW822252 | -0,6974 | 2,223E-02 | X | transcribed_unprocessed_pseudogene | expressed sequence AW822252 [Source:MGI Symbol;Acc:MGI:2148030] |
| E330035G20Rik | 3,5987 | 2,226E-02 | 12 | lincRNA | RIKEN cDNA E330035G20 gene [Source:MGI Symbol;Acc:MGI:2443932] |
| Gm12944 | 2,8311 | 2,227E-02 | 4 | processed_pseudogene | predicted gene 12944 [Source:MGI Symbol;Acc:MGI:3650498] |
| Parp14 | 0,3895 | 2,228E-02 | 16 | protein_coding | poly (ADP-ribose) polymerase family, member 14 [Source:MGI Symbol;Acc:MGI:1919489] |
| Naf1 | 0,2710 | 2,232E-02 | 8 | protein_coding | nuclear assembly factor 1 ribonucleoprotein [Source:MGI Symbol;Acc:MGI:2682306] |
| Gm28319 | 2,2694 | 2,232E-02 | 1 | antisense | predicted gene 28319 [Source:MGI Symbol;Acc:MGI:5579025] |
| Gm28439 | 0,5594 | 2,232E-02 | 1 | unprocessed_pseudogene | predicted gene 28439 [Source:MGI Symbol;Acc:MGI:5579145] |
| Cntn1 | 0,1584 | 2,233E-02 | 15 | protein_coding | contactin 1 [Source:MGI Symbol;Acc:MGI:105980] |
| E230015B07Rik | 2,8898 | 2,235E-02 | 6 | bidirectional_promoter_lncRNA | RIKEN cDNA E230015B07 gene [Source:MGI Symbol;Acc:MGI:2443140] |
| Cul2 | 0,1751 | 2,241E-02 | 18 | protein_coding | cullin 2 [Source:MGI Symbol;Acc:MGI:1918995] |
| Frat1 | -0,2342 | 2,246E-02 | 19 | protein_coding | frequently rearranged in advanced T cell lymphomas [Source:MGI Symbol;Acc:MGI:109450] |
| Rnase1 | 1,1856 | 2,257E-02 | 14 | protein_coding | ribonuclease, RNase A family, 1 (pancreatic) [Source:MGI Symbol;Acc:MGI:97919] |
| Kat8 | -0,2119 | 2,260E-02 | 7 | protein_coding | K(lysine) acetyltransferase 8 [Source:MGI Symbol;Acc:MGI:1915023] |
| Bpifb5 | 3,6321 | 2,264E-02 | 2 | protein_coding | BPI fold containing family B, member 5 [Source:MGI Symbol;Acc:MGI:2385160] |
| Cwc15 | 0,1851 | 2,265E-02 | 9 | protein_coding | CWC15 spliceosome-associated protein [Source:MGI Symbol;Acc:MGI:1913320] |
| Kcnk13 | -0,4000 | 2,266E-02 | 12 | protein_coding | potassium channel, subfamily K, member 13 [Source:MGI Symbol;Acc:MGI:2384976] |
| Adam12 | -0,3758 | 2,267E-02 | 7 | protein_coding | a disintegrin and metallopeptidase domain 12 (meltrin alpha) [Source:MGI Symbol;Acc:MGI:105378] |
| Itfg2 | -0,2349 | 2,272E-02 | 6 | protein_coding | integrin alpha FG-GAP repeat containing 2 [Source:MGI Symbol;Acc:MGI:1915450] |
| Paqr7 | -0,1993 | 2,274E-02 | 4 | protein_coding | progestin and adipoQ receptor family member VII [Source:MGI Symbol;Acc:MGI:1919154] |
| Gm12847 | -2,7677 | 2,276E-02 | 4 | lincRNA | predicted gene 12847 [Source:MGI Symbol;Acc:MGI:3650822] |
| Kcnj11 | -0,2245 | 2,276E-02 | 7 | protein_coding | potassium inwardly rectifying channel, subfamily J, member 11 [Source:MGI Symbol;Acc:MGI:107501] |
| Ska3 | 0,8041 | 2,286E-02 | 14 | protein_coding | spindle and kinetochore associated complex subunit 3 [Source:MGI Symbol;Acc:MGI:3041235] |
| Tmem141 | -0,2901 | 2,291E-02 | 2 | protein_coding | transmembrane protein 141 [Source:MGI Symbol;Acc:MGI:1098773] |
| Perm1 | 0,3094 | 2,318E-02 | 4 | protein_coding | PPARGC1 and ESRR induced regulator, muscle 1 [Source:MGI Symbol;Acc:MGI:1921433] |

| | | | | | |
|---|---|---|---|---|---|
| Fosl2 | 0,1697 | 2,333E-02 | 5 | protein_coding | fos-like antigen 2 [Source:MGI Symbol;Acc:MGI:102858] |
| Tusc1 | -0,3051 | 2,340E-02 | 4 | protein_coding | tumor suppressor candidate 1 [Source:MGI Symbol;Acc:MGI:2684283] |
| Kcnab3 | -0,2543 | 2,343E-02 | 11 | protein_coding | potassium voltage-gated channel, shaker-related subfamily, beta member 3 [Source:MGI Symbol;Acc:MGI:1336208] |
| Map1lc3b | -0,1226 | 2,344E-02 | 8 | protein_coding | microtubule-associated protein 1 light chain 3 beta [Source:MGI Symbol;Acc:MGI:1914693] |
| Lbr | 0,2801 | 2,347E-02 | 1 | protein_coding | lamin B receptor [Source:MGI Symbol;Acc:MGI:2138281] |
| Sik1 | 0,2848 | 2,348E-02 | 17 | protein_coding | salt inducible kinase 1 [Source:MGI Symbol;Acc:MGI:104754] |
| Ddx18 | 0,2398 | 2,357E-02 | 1 | protein_coding | DEAD (Asp-Glu-Ala-Asp) box polypeptide 18 [Source:MGI Symbol;Acc:MGI:1914192] |
| Slc12a8 | 0,7117 | 2,368E-02 | 16 | protein_coding | solute carrier family 12 (potassium/chloride transporters), member 8 [Source:MGI Symbol;Acc:MGI:2443672] |
| Gm17201 | -0,4025 | 2,371E-02 | 10 | antisense | predicted gene 17201 [Source:MGI Symbol;Acc:MGI:4938028] |
| Itfg1 | 0,1557 | 2,373E-02 | 8 | protein_coding | integrin alpha FG-GAP repeat containing 1 [Source:MGI Symbol;Acc:MGI:106419] |
| Ahr | 0,3365 | 2,373E-02 | 12 | protein_coding | aryl-hydrocarbon receptor [Source:MGI Symbol;Acc:MGI:105043] |
| Hnf1b | -1,0206 | 2,377E-02 | 11 | protein_coding | HNF1 homeobox B [Source:MGI Symbol;Acc:MGI:98505] |
| Gm10544 | -1,3227 | 2,378E-02 | 18 | processed_transcript | predicted gene 10544 [Source:MGI Symbol;Acc:MGI:3642892] |
| Commd4 | -0,1627 | 2,380E-02 | 9 | protein_coding | COMM domain containing 4 [Source:MGI Symbol;Acc:MGI:1913449] |
| Sipa1l3 | -0,2152 | 2,397E-02 | 7 | protein_coding | signal-induced proliferation-associated 1 like 3 [Source:MGI Symbol;Acc:MGI:1921456] |
| Rnf208 | -0,1604 | 2,400E-02 | 2 | protein_coding | ring finger protein 208 [Source:MGI Symbol;Acc:MGI:1916096] |
| Acap3 | -0,1476 | 2,402E-02 | 4 | protein_coding | ArfGAP with coiled-coil, ankyrin repeat and PH domains 3 [Source:MGI Symbol;Acc:MGI:2153589] |
| Gm13625 | 1,8132 | 2,405E-02 | 2 | processed_transcript | predicted gene 13625 [Source:MGI Symbol;Acc:MGI:3709621] |
| Rgl3 | -0,4252 | 2,407E-02 | 9 | protein_coding | ral guanine nucleotide dissociation stimulator-like 3 [Source:MGI Symbol;Acc:MGI:1918996] |
| Otogl | -0,7234 | 2,411E-02 | 10 | protein_coding | otogelin-like [Source:MGI Symbol;Acc:MGI:3647600] |
| Nabp2 | -0,1982 | 2,416E-02 | 10 | protein_coding | nucleic acid binding protein 2 [Source:MGI Symbol;Acc:MGI:1917167] |
| mt-Nd2 | 0,2525 | 2,417E-02 | MT | protein_coding | mitochondrially encoded NADH dehydrogenase 2 [Source:MGI Symbol;Acc:MGI:102500] |
| Mios | 0,2630 | 2,418E-02 | 6 | protein_coding | meiosis regulator for oocyte development [Source:MGI Symbol;Acc:MGI:2182066] |
| Gm37804 | -3,9358 | 2,418E-02 | 12 | TEC | predicted gene, 37804 [Source:MGI Symbol;Acc:MGI:5611032] |
| Wdr3 | 0,2454 | 2,420E-02 | 3 | protein_coding | WD repeat domain 3 [Source:MGI Symbol;Acc:MGI:2443143] |
| Slc25a45 | 0,4788 | 2,421E-02 | 19 | protein_coding | solute carrier family 25, member 45 [Source:MGI Symbol;Acc:MGI:2147731] |
| Gm3411 | 1,2857 | 2,428E-02 | 14 | protein_coding | predicted gene 3411 [Source:MGI Symbol;Acc:MGI:3781589] |

| | | | | | |
|---|---|---|---|---|---|
| Zdhhc13 | 0,2476 | 2,439E-02 | 7 | protein_coding | zinc finger, DHHC domain containing 13 [Source:MGI Symbol;Acc:MGI:1919227] |
| Tpgs1 | -0,2262 | 2,446E-02 | 10 | protein_coding | tubulin polyglutamylase complex subunit 1 [Source:MGI Symbol;Acc:MGI:106618] |
| Sart3 | -0,1774 | 2,447E-02 | 5 | protein_coding | squamous cell carcinoma antigen recognized by T cells 3 [Source:MGI Symbol;Acc:MGI:1858230] |
| Mir1291 | 2,2650 | 2,461E-02 | 15 | miRNA | microRNA 1291 [Source:MGI Symbol;Acc:MGI:5562744] |
| 4930520O04Rik | -0,7088 | 2,469E-02 | 9 | processed_transcript | RIKEN cDNA 4930520O04 gene [Source:MGI Symbol;Acc:MGI:1922366] |
| Phf10 | 0,1926 | 2,472E-02 | 17 | protein_coding | PHD finger protein 10 [Source:MGI Symbol;Acc:MGI:1919307] |
| Gm42775 | 3,2346 | 2,472E-02 | 9 | lincRNA | predicted gene 42775 [Source:MGI Symbol;Acc:MGI:5662912] |
| CT010429,1 | 3,6941 | 2,474E-02 | 17 | processed_pseudogene | ribosomal protein S27 (Rps27) pseudogene |
| Mapre1 | 0,1388 | 2,485E-02 | 2 | protein_coding | microtubule-associated protein, RP/EB family, member 1 [Source:MGI Symbol;Acc:MGI:891995] |
| Nxpe4 | 0,2743 | 2,489E-02 | 9 | protein_coding | neurexophilin and PC-esterase domain family, member 4 [Source:MGI Symbol;Acc:MGI:1924792] |
| Wipi1 | 0,1991 | 2,491E-02 | 11 | protein_coding | WD repeat domain, phosphoinositide interacting 1 [Source:MGI Symbol;Acc:MGI:1261864] |
| Gm14762 | 3,3783 | 2,498E-02 | X | lincRNA | predicted gene 14762 [Source:MGI Symbol;Acc:MGI:3705212] |
| Gm5518 | 0,2833 | 2,498E-02 | 19 | processed_pseudogene | predicted gene 5518 [Source:MGI Symbol;Acc:MGI:3648673] |
| Mtbp | 0,7129 | 2,502E-02 | 15 | protein_coding | Mdm2, transformed 3T3 cell double minute p53 binding protein [Source:MGI Symbol;Acc:MGI:2146005] |
| Mapk8ip1 | -0,1475 | 2,511E-02 | 2 | protein_coding | mitogen-activated protein kinase 8 interacting protein 1 [Source:MGI Symbol;Acc:MGI:1309464] |
| Exosc5 | -0,2813 | 2,524E-02 | 7 | protein_coding | exosome component 5 [Source:MGI Symbol;Acc:MGI:107889] |
| A330009N23Rik | -0,5119 | 2,527E-02 | 15 | antisense | RIKEN cDNA A330009N23 gene [Source:MGI Symbol;Acc:MGI:2443491] |
| Gm14412 | 1,0220 | 2,528E-02 | 2 | protein_coding | predicted gene 14412 [Source:MGI Symbol;Acc:MGI:3652251] |
| Rbm41 | 0,4696 | 2,532E-02 | X | protein_coding | RNA binding motif protein 41 [Source:MGI Symbol;Acc:MGI:2444923] |
| Gm14104 | 2,6079 | 2,534E-02 | 2 | antisense | predicted gene 14104 [Source:MGI Symbol;Acc:MGI:3651500] |
| Gpr137 | -0,1812 | 2,547E-02 | 19 | protein_coding | G protein-coupled receptor 137 [Source:MGI Symbol;Acc:MGI:2147529] |
| Msh6 | 0,2374 | 2,547E-02 | 17 | protein_coding | mutS homolog 6 [Source:MGI Symbol;Acc:MGI:1343961] |
| Gm42600 | 0,9680 | 2,551E-02 | 6 | TEC | predicted gene 42600 [Source:MGI Symbol;Acc:MGI:5662737] |
| Gm48904 | 3,3104 | 2,555E-02 | 12 | TEC | predicted gene, 48904 [Source:MGI Symbol;Acc:MGI:6098675] |
| Gm27252 | 1,1548 | 2,558E-02 | 7 | processed_transcript | predicted gene 27252 [Source:MGI Symbol;Acc:MGI:5521095] |
| Depdc5 | -0,1539 | 2,571E-02 | 5 | protein_coding | DEP domain containing 5 [Source:MGI Symbol;Acc:MGI:2141101] |
| Gm29113 | 3,9317 | 2,572E-02 | 1 | lincRNA | predicted gene 29113 [Source:MGI Symbol;Acc:MGI:5579819] |

| | | | | | |
|---|---|---|---|---|---|
| Ak6 | -0,3183 | 2,576E-02 | 13 | protein_coding | adenylate kinase 6 [Source:MGI Symbol;Acc:MGI:5510732] |
| Slamf1 | 3,6803 | 2,577E-02 | 1 | protein_coding | signaling lymphocytic activation molecule family member 1 [Source:MGI Symbol;Acc:MGI:1351314] |
| Ngp | -0,9833 | 2,578E-02 | 9 | protein_coding | neutrophilic granule protein [Source:MGI Symbol;Acc:MGI:105983] |
| Inpp5j | -0,1875 | 2,579E-02 | 11 | protein_coding | inositol polyphosphate 5-phosphatase J [Source:MGI Symbol;Acc:MGI:2158663] |
| Rnaseh1 | -0,2761 | 2,580E-02 | 12 | protein_coding | ribonuclease H1 [Source:MGI Symbol;Acc:MGI:1335073] |
| Commd5 | -1,3174 | 2,592E-02 | 15 | protein_coding | COMM domain containing 5 [Source:MGI Symbol;Acc:MGI:1913648] |
| Klhdc1 | 0,2696 | 2,593E-02 | 12 | protein_coding | kelch domain containing 1 [Source:MGI Symbol;Acc:MGI:2672853] |
| 0610012G03Rik | -0,2340 | 2,603E-02 | 16 | protein_coding | RIKEN cDNA 0610012G03 gene [Source:MGI Symbol;Acc:MGI:1913301] |
| 4930439A04Rik | 3,8703 | 2,614E-02 | 1 | antisense | RIKEN cDNA 4930439A04 gene [Source:MGI Symbol;Acc:MGI:1925369] |
| Slc25a38 | -0,2050 | 2,620E-02 | 9 | protein_coding | solute carrier family 25, member 38 [Source:MGI Symbol;Acc:MGI:2384782] |
| Gm44220 | 0,5117 | 2,622E-02 | 6 | TEC | predicted gene, 44220 [Source:MGI Symbol;Acc:MGI:5690612] |
| A830019P07Rik | 0,7893 | 2,625E-02 | 19 | lincRNA | RIKEN cDNA A830019P07 gene [Source:MGI Symbol;Acc:MGI:3028035] |
| Tagln | 1,1356 | 2,628E-02 | 9 | protein_coding | transgelin [Source:MGI Symbol;Acc:MGI:106012] |
| Stac | -0,3445 | 2,631E-02 | 9 | protein_coding | src homology three (SH3) and cysteine rich domain [Source:MGI Symbol;Acc:MGI:1201400] |
| Gsk3a | -0,1596 | 2,633E-02 | 7 | protein_coding | glycogen synthase kinase 3 alpha [Source:MGI Symbol;Acc:MGI:2152453] |
| Rab1b | -0,1674 | 2,635E-02 | 19 | protein_coding | RAB1B, member RAS oncogene family [Source:MGI Symbol;Acc:MGI:1923558] |
| Gm1604a | 3,5931 | 2,647E-02 | 17 | protein_coding | predicted gene 1604A [Source:MGI Symbol;Acc:MGI:3807545] |
| A730017L22Rik | 0,3117 | 2,651E-02 | 2 | processed_transcript | RIKEN cDNA A730017L22 gene [Source:MGI Symbol;Acc:MGI:3584452] |
| Zfp710 | -0,1867 | 2,655E-02 | 7 | protein_coding | zinc finger protein 710 [Source:MGI Symbol;Acc:MGI:1921747] |
| Selenow | -0,1648 | 2,660E-02 | 7 | protein_coding | selenoprotein W [Source:MGI Symbol;Acc:MGI:1100878] |
| Pwwp2b | -0,2319 | 2,666E-02 | 7 | protein_coding | PWWP domain containing 2B [Source:MGI Symbol;Acc:MGI:2142008] |
| Gm12472 | -2,1863 | 2,668E-02 | 4 | antisense | predicted gene 12472 [Source:MGI Symbol;Acc:MGI:3649439] |
| Oplah | -0,1881 | 2,681E-02 | 15 | protein_coding | 5-oxoprolinase (ATP-hydrolysing) [Source:MGI Symbol;Acc:MGI:1922725] |
| F12 | -1,2892 | 2,682E-02 | 13 | protein_coding | coagulation factor XII (Hageman factor) [Source:MGI Symbol;Acc:MGI:1891012] |
| Gm16539 | 3,8802 | 2,688E-02 | 1 | unprocessed_pseudogene | predicted gene 16539 [Source:MGI Symbol;Acc:MGI:4414959] |
| Rfx4 | 0,2778 | 2,689E-02 | 10 | protein_coding | regulatory factor X, 4 (influences HLA class II expression) [Source:MGI Symbol;Acc:MGI:1918387] |
| 1600002K03Rik | -0,4984 | 2,699E-02 | 10 | protein_coding | RIKEN cDNA 1600002K03 gene [Source:MGI Symbol;Acc:MGI:1917020] |

| | | | | | |
|---|---|---|---|---|---|
| Rasa2 | 0,2374 | 2,700E-02 | 9 | protein_coding | RAS p21 protein activator 2 [Source:MGI Symbol;Acc:MGI:2149960] |
| Gm47715 | -2,2320 | 2,701E-02 | 10 | lincRNA | predicted gene, 47715 [Source:MGI Symbol;Acc:MGI:6096840] |
| Gbp5 | 0,5008 | 2,708E-02 | 3 | protein_coding | guanylate binding protein 5 [Source:MGI Symbol;Acc:MGI:2429943] |
| Ptpn12 | 0,1983 | 2,710E-02 | 5 | protein_coding | protein tyrosine phosphatase, non-receptor type 12 [Source:MGI Symbol;Acc:MGI:104673] |
| Gabrg1 | 0,3293 | 2,712E-02 | 5 | protein_coding | gamma-aminobutyric acid (GABA) A receptor, subunit gamma 1 [Source:MGI Symbol;Acc:MGI:103156] |
| Gm22663 | 3,2753 | 2,725E-02 | 12 | snRNA | predicted gene, 22663 [Source:MGI Symbol;Acc:MGI:5452440] |
| Gm43421 | 2,1099 | 2,730E-02 | 5 | TEC | predicted gene 43421 [Source:MGI Symbol;Acc:MGI:5663558] |
| Gm26834 | -1,4905 | 2,733E-02 | 16 | lincRNA | predicted gene, 26834 [Source:MGI Symbol;Acc:MGI:5477328] |
| AC154707,1 | 0,4704 | 2,735E-02 | 17 | processed_pseudogene | novel C2H2 zinc finger pseudogene |
| Btbd2 | -0,1622 | 2,738E-02 | 10 | protein_coding | BTB (POZ) domain containing 2 [Source:MGI Symbol;Acc:MGI:1933831] |
| Chchd4 | -0,2059 | 2,742E-02 | 6 | protein_coding | coiled-coil-helix-coiled-coil-helix domain containing 4 [Source:MGI Symbol;Acc:MGI:1919420] |
| Rpl24 | -1,1112 | 2,747E-02 | 16 | protein_coding | ribosomal protein L24 [Source:MGI Symbol;Acc:MGI:1915443] |
| Zfp219 | -0,2440 | 2,754E-02 | 14 | protein_coding | zinc finger protein 219 [Source:MGI Symbol;Acc:MGI:1917140] |
| Htra1 | -0,2022 | 2,761E-02 | 7 | protein_coding | HtrA serine peptidase 1 [Source:MGI Symbol;Acc:MGI:1929076] |
| Yes1 | 0,3221 | 2,762E-02 | 5 | protein_coding | YES proto-oncogene 1, Src family tyrosine kinase [Source:MGI Symbol;Acc:MGI:99147] |
| 4930538E20Rik | 1,1210 | 2,766E-02 | 11 | antisense | RIKEN cDNA 4930538E20 gene [Source:MGI Symbol;Acc:MGI:1925450] |
| Sbf1 | -0,1335 | 2,766E-02 | 15 | protein_coding | SET binding factor 1 [Source:MGI Symbol;Acc:MGI:1925230] |
| Lect2 | 3,3780 | 2,771E-02 | 13 | protein_coding | leukocyte cell-derived chemotaxin 2 [Source:MGI Symbol;Acc:MGI:1278342] |
| Gm36939 | 3,8321 | 2,783E-02 | 16 | TEC | predicted gene, 36939 [Source:MGI Symbol;Acc:MGI:5610167] |
| Gm13340 | 0,3293 | 2,785E-02 | 2 | unprocessed_pseudogene | predicted gene 13340 [Source:MGI Symbol;Acc:MGI:3650227] |
| Gm26765 | -1,5883 | 2,789E-02 | 10 | lincRNA | predicted gene, 26765 [Source:MGI Symbol;Acc:MGI:5477259] |
| Npas1 | -0,2195 | 2,795E-02 | 7 | protein_coding | neuronal PAS domain protein 1 [Source:MGI Symbol;Acc:MGI:109205] |
| Vegfb | -0,1938 | 2,799E-02 | 19 | protein_coding | vascular endothelial growth factor B [Source:MGI Symbol;Acc:MGI:106199] |
| Tnfrsf4 | -0,7562 | 2,821E-02 | 4 | protein_coding | tumor necrosis factor receptor superfamily, member 4 [Source:MGI Symbol;Acc:MGI:104512] |
| Nt5m | -0,1943 | 2,823E-02 | 11 | protein_coding | 5',3'-nucleotidase, mitochondrial [Source:MGI Symbol;Acc:MGI:1917127] |
| Dync2li1 | 0,2514 | 2,831E-02 | 17 | protein_coding | dynein cytoplasmic 2 light intermediate chain 1 [Source:MGI Symbol;Acc:MGI:1913996] |
| Mettl7a3 | -2,6690 | 2,839E-02 | 15 | protein_coding | methyltransferase like 7A3 [Source:MGI Symbol;Acc:MGI:3710670] |

| | | | | | |
|---|---|---|---|---|---|
| Zfp1 | 0,2141 | 2,840E-02 | 8 | protein_coding | zinc finger protein 1 [Source:MGI Symbol;Acc:MGI:99154] |
| Nsmf | -0,1596 | 2,842E-02 | 2 | protein_coding | NMDA receptor synaptonuclear signaling and neuronal migration factor [Source:MGI Symbol;Acc:MGI:1861755] |
| Orai2 | -0,1467 | 2,846E-02 | 5 | protein_coding | ORAI calcium release-activated calcium modulator 2 [Source:MGI Symbol;Acc:MGI:2443195] |
| BC023105 | 2,7111 | 2,852E-02 | 18 | pseudogene | cDNA sequence BC023105 [Source:MGI Symbol;Acc:MGI:2384767] |
| 4930404H11Rik | 3,5768 | 2,854E-02 | 12 | lincRNA | RIKEN cDNA 4930404H11 gene [Source:MGI Symbol;Acc:MGI:1921065] |
| Gm12031 | 3,3917 | 2,855E-02 | 11 | lincRNA | predicted gene 12031 [Source:MGI Symbol;Acc:MGI:3651406] |
| Gprasp1 | 0,1094 | 2,855E-02 | X | protein_coding | G protein-coupled receptor associated sorting protein 1 [Source:MGI Symbol;Acc:MGI:1917418] |
| Olfr56 | 3,9231 | 2,859E-02 | 11 | protein_coding | olfactory receptor 56 [Source:MGI Symbol;Acc:MGI:1333785] |
| Phxr4 | -2,3740 | 2,870E-02 | 9 | TEC | per-hexamer repeat gene 4 [Source:MGI Symbol;Acc:MGI:104522] |
| Gm12925 | 0,8105 | 2,871E-02 | 4 | antisense | predicted gene 12925 [Source:MGI Symbol;Acc:MGI:3651536] |
| Sptbn2 | -0,1449 | 2,874E-02 | 19 | protein_coding | spectrin beta, non-erythrocytic 2 [Source:MGI Symbol;Acc:MGI:1313261] |
| Slc51a | 3,6688 | 2,874E-02 | 16 | protein_coding | solute carrier family 51, alpha subunit [Source:MGI Symbol;Acc:MGI:2146634] |
| Tsga10ip | 2,7258 | 2,887E-02 | 19 | protein_coding | testis specific 10 interacting protein [Source:MGI Symbol;Acc:MGI:1925556] |
| Fbxo44 | -0,1559 | 2,893E-02 | 4 | protein_coding | F-box protein 44 [Source:MGI Symbol;Acc:MGI:1354744] |
| Arl13b | 0,3284 | 2,895E-02 | 16 | protein_coding | ADP-ribosylation factor-like 13B [Source:MGI Symbol;Acc:MGI:1915396] |
| Gpx1 | -0,2067 | 2,900E-02 | 9 | protein_coding | glutathione peroxidase 1 [Source:MGI Symbol;Acc:MGI:104887] |
| Eme1 | -2,8871 | 2,906E-02 | 11 | protein_coding | essential meiotic structure-specific endonuclease 1 [Source:MGI Symbol;Acc:MGI:3576783] |
| Esf1 | 0,2541 | 2,907E-02 | 2 | protein_coding | ESF1 nucleolar pre-rRNA processing protein homolog [Source:MGI Symbol;Acc:MGI:1913830] |
| Gm34388 | -3,3722 | 2,919E-02 | 13 | lincRNA | predicted gene, 34388 [Source:MGI Symbol;Acc:MGI:5593547] |
| Pold1 | -0,2981 | 2,942E-02 | 7 | protein_coding | polymerase (DNA directed), delta 1, catalytic subunit [Source:MGI Symbol;Acc:MGI:97741] |
| Rap1gap | -0,1590 | 2,953E-02 | 4 | protein_coding | Rap1 GTPase-activating protein [Source:MGI Symbol;Acc:MGI:109338] |
| Rpl21-ps8 | 0,4139 | 2,971E-02 | 18 | processed_pseudogene | ribosomal protein L21, pseudogene 8 [Source:MGI Symbol;Acc:MGI:3648345] |
| Gm31410 | 3,5553 | 2,972E-02 | 9 | lincRNA | predicted gene, 31410 [Source:MGI Symbol;Acc:MGI:5590569] |
| Sin3a | 0,1796 | 2,975E-02 | 9 | protein_coding | transcriptional regulator, SIN3A (yeast) [Source:MGI Symbol;Acc:MGI:107157] |
| Arhgap35 | -0,1142 | 2,979E-02 | 7 | protein_coding | Rho GTPase activating protein 35 [Source:MGI Symbol;Acc:MGI:1929494] |
| Aspa | 0,2553 | 2,995E-02 | 11 | protein_coding | aspartoacylase [Source:MGI Symbol;Acc:MGI:87914] |
| Sema5b | -0,2684 | 3,002E-02 | 16 | protein_coding | (semaphorin) 5B [Source:MGI Symbol;Acc:MGI:107555] |

| | | | | | |
|---|---|---|---|---|---|
| Akap2 | 3,2079 | 3,009E-02 | 4 | protein_coding | A kinase (PRKA) anchor protein 2 [Source:MGI Symbol;Acc:MGI:1306795] |
| Gm13217 | 1,1431 | 3,015E-02 | 2 | unprocessed_pseudogene | predicted gene 13217 [Source:MGI Symbol;Acc:MGI:3649713] |
| Uba3 | 0,2184 | 3,016E-02 | 6 | protein_coding | ubiquitin-like modifier activating enzyme 3 [Source:MGI Symbol;Acc:MGI:1341217] |
| Fam69b | -0,1727 | 3,018E-02 | 2 | protein_coding | family with sequence similarity 69, member B [Source:MGI Symbol;Acc:MGI:1927576] |
| Gm47594 | -3,0345 | 3,031E-02 | 10 | TEC | predicted gene, 47594 [Source:MGI Symbol;Acc:MGI:6096642] |
| Prelid1 | -0,1606 | 3,033E-02 | 13 | protein_coding | PRELI domain containing 1 [Source:MGI Symbol;Acc:MGI:1913744] |
| Gm44646 | -0,9312 | 3,038E-02 | 7 | processed_transcript | predicted gene 44646 [Source:MGI Symbol;Acc:MGI:5753222] |
| Adgrg5 | 4,0453 | 3,043E-02 | 8 | protein_coding | adhesion G protein-coupled receptor G5 [Source:MGI Symbol;Acc:MGI:2685955] |
| Gm10392 | 2,9820 | 3,058E-02 | 11 | protein_coding | predicted gene 10392 [Source:MGI Symbol;Acc:MGI:3704377] |
| Fgfbp3 | 0,4188 | 3,065E-02 | 19 | protein_coding | fibroblast growth factor binding protein 3 [Source:MGI Symbol;Acc:MGI:1919764] |
| 4930556I23Rik | -0,6050 | 3,065E-02 | 1 | antisense | RIKEN cDNA 4930556I23 gene [Source:MGI Symbol;Acc:MGI:1922520] |
| Mir6982 | -1,0838 | 3,072E-02 | 18 | miRNA | microRNA 6982 [Source:MGI Symbol;Acc:MGI:5531272] |
| Ndnf | 0,3457 | 3,074E-02 | 6 | protein_coding | neuron-derived neurotrophic factor [Source:MGI Symbol;Acc:MGI:1915419] |
| Map6d1 | -0,1698 | 3,079E-02 | 16 | protein_coding | MAP6 domain containing 1 [Source:MGI Symbol;Acc:MGI:3607784] |
| Snapc1 | 0,2045 | 3,083E-02 | 12 | protein_coding | small nuclear RNA activating complex, polypeptide 1 [Source:MGI Symbol;Acc:MGI:1922877] |
| Ccdc62 | 0,4637 | 3,084E-02 | 5 | protein_coding | coiled-coil domain containing 62 [Source:MGI Symbol;Acc:MGI:2684996] |
| Clip2 | -0,1292 | 3,088E-02 | 5 | protein_coding | CAP-GLY domain containing linker protein 2 [Source:MGI Symbol;Acc:MGI:1313136] |
| B930086L07Rik | 2,8759 | 3,090E-02 | 7 | TEC | RIKEN cDNA B930086L07 gene [Source:MGI Symbol;Acc:MGI:2443202] |
| Sarm1 | -0,1803 | 3,090E-02 | 11 | protein_coding | sterile alpha and HEAT/Armadillo motif containing 1 [Source:MGI Symbol;Acc:MGI:2136419] |
| Timd4 | -4,0506 | 3,091E-02 | 11 | protein_coding | T cell immunoglobulin and mucin domain containing 4 [Source:MGI Symbol;Acc:MGI:2445125] |
| Gm21399 | 0,9799 | 3,099E-02 | 8 | processed_pseudogene | predicted gene, 21399 [Source:MGI Symbol;Acc:MGI:5434754] |
| Efcab2 | 0,3375 | 3,100E-02 | 1 | protein_coding | EF-hand calcium binding domain 2 [Source:MGI Symbol;Acc:MGI:1915476] |
| Ramp3 | -0,3998 | 3,105E-02 | 11 | protein_coding | receptor (calcitonin) activity modifying protein 3 [Source:MGI Symbol;Acc:MGI:1860292] |
| Kdm6a | 0,2850 | 3,105E-02 | X | protein_coding | lysine (K)-specific demethylase 6A [Source:MGI Symbol;Acc:MGI:1095419] |
| Lipo3 | 0,3628 | 3,105E-02 | 19 | protein_coding | lipase, member O3 [Source:MGI Symbol;Acc:MGI:2147592] |
| Usp28 | 0,2111 | 3,105E-02 | 9 | protein_coding | ubiquitin specific peptidase 28 [Source:MGI Symbol;Acc:MGI:2442293] |
| CT025652,1 | 1,0055 | 3,109E-02 | 17 | antisense | novel transcript, antisense to Fance |

| | | | | | |
|---|---|---|---|---|---|
| Nbn | 0,3385 | 3,114E-02 | 4 | protein_coding | nibrin [Source:MGI Symbol;Acc:MGI:1351625] |
| Rttn | 0,4304 | 3,115E-02 | 18 | protein_coding | rotatin [Source:MGI Symbol;Acc:MGI:2179288] |
| Gm43670 | -1,7407 | 3,137E-02 | 5 | TEC | predicted gene 43670 [Source:MGI Symbol;Acc:MGI:5663807] |
| Tlr2 | 0,6899 | 3,146E-02 | 3 | protein_coding | toll-like receptor 2 [Source:MGI Symbol;Acc:MGI:1346060] |
| Neurog1 | 1,6555 | 3,165E-02 | 13 | protein_coding | neurogenin 1 [Source:MGI Symbol;Acc:MGI:107754] |
| Best1 | -0,5838 | 3,175E-02 | 19 | protein_coding | bestrophin 1 [Source:MGI Symbol;Acc:MGI:1346332] |
| Trim80 | 2,9306 | 3,178E-02 | 11 | protein_coding | tripartite motif-containing 80 [Source:MGI Symbol;Acc:MGI:3588186] |
| Gm19461 | -1,2458 | 3,180E-02 | 1 | antisense | predicted gene, 19461 [Source:MGI Symbol;Acc:MGI:5011646] |
| Gm21057 | -2,5569 | 3,181E-02 | 7 | antisense | predicted gene, 21057 [Source:MGI Symbol;Acc:MGI:5434414] |
| Gm37824 | 0,6848 | 3,186E-02 | 2 | TEC | predicted gene, 37824 [Source:MGI Symbol;Acc:MGI:5611052] |
| Gm6505 | -1,2038 | 3,188E-02 | 3 | processed_pseudogene | predicted pseudogene 6505 [Source:MGI Symbol;Acc:MGI:3648080] |
| Gm26691 | 1,0927 | 3,188E-02 | 3 | antisense | predicted gene, 26691 [Source:MGI Symbol;Acc:MGI:5477185] |
| Ccar1 | 0,2368 | 3,190E-02 | 10 | protein_coding | cell division cycle and apoptosis regulator 1 [Source:MGI Symbol;Acc:MGI:1914750] |
| 4930435N07Rik | 1,3968 | 3,191E-02 | 8 | TEC | RIKEN cDNA 4930435N07 gene [Source:MGI Symbol;Acc:MGI:1923047] |
| Mrps11 | -0,2805 | 3,194E-02 | 7 | protein_coding | mitochondrial ribosomal protein S11 [Source:MGI Symbol;Acc:MGI:1915244] |
| Ccdc22 | -3,8571 | 3,200E-02 | X | protein_coding | coiled-coil domain containing 22 [Source:MGI Symbol;Acc:MGI:1859608] |
| Efemp1 | 0,3404 | 3,200E-02 | 11 | protein_coding | epidermal growth factor-containing fibulin-like extracellular matrix protein 1 [Source:MGI Symbol;Acc:MGI:1339998] |
| Gm6238 | -3,6812 | 3,202E-02 | X | processed_pseudogene | predicted pseudogene 6238 [Source:MGI Symbol;Acc:MGI:3648913] |
| Cyb5a | 0,1777 | 3,204E-02 | 18 | protein_coding | cytochrome b5 type A (microsomal) [Source:MGI Symbol;Acc:MGI:1926952] |
| Tyw5 | 0,3382 | 3,205E-02 | 1 | protein_coding | tRNA-yW synthesizing protein 5 [Source:MGI Symbol;Acc:MGI:1915986] |
| Gm8116 | -0,5839 | 3,224E-02 | 9 | processed_pseudogene | predicted gene 8116 [Source:MGI Symbol;Acc:MGI:3648797] |
| Dgkz | -0,1545 | 3,225E-02 | 2 | protein_coding | diacylglycerol kinase zeta [Source:MGI Symbol;Acc:MGI:1278339] |
| Zfp65 | 0,3073 | 3,226E-02 | 13 | protein_coding | zinc finger protein 65 [Source:MGI Symbol;Acc:MGI:107769] |
| AC167169,1 | 1,6913 | 3,232E-02 | 16 | antisense | novel transcript, antisense to Dscam |
| Exoc1 | 0,1598 | 3,233E-02 | 5 | protein_coding | exocyst complex component 1 [Source:MGI Symbol;Acc:MGI:2445020] |
| Gm9924 | -0,9501 | 3,238E-02 | 5 | TEC | predicted gene 9924 [Source:MGI Symbol;Acc:MGI:3642216] |
| Gtf2h2 | 0,1968 | 3,239E-02 | 13 | protein_coding | general transcription factor II H, polypeptide 2 [Source:MGI Symbol;Acc:MGI:1345669] |

| | | | | | |
|---|---|---|---|---|---|
| Rprml | -0,1830 | 3,243E-02 | 11 | protein_coding | reprimo-like [Source:MGI Symbol;Acc:MGI:2144486] |
| Gm14863 | -3,2785 | 3,266E-02 | X | processed_pseudogene | predicted gene 14863 [Source:MGI Symbol;Acc:MGI:3705489] |
| Ccdc13 | -0,3391 | 3,269E-02 | 9 | protein_coding | coiled-coil domain containing 13 [Source:MGI Symbol;Acc:MGI:1920144] |
| Gm24270 | 0,5978 | 3,275E-02 | 9 | miRNA | predicted gene, 24270 [Source:MGI Symbol;Acc:MGI:5454047] |
| Gm47445 | -2,0990 | 3,279E-02 | 10 | TEC | predicted gene, 47445 [Source:MGI Symbol;Acc:MGI:6096399] |
| Gm42748 | -1,6271 | 3,279E-02 | 5 | TEC | predicted gene 42748 [Source:MGI Symbol;Acc:MGI:5662885] |
| Gm10231 | 2,0362 | 3,279E-02 | 17 | processed_pseudogene | predicted pseudogene 10231 [Source:MGI Symbol;Acc:MGI:3704468] |
| Lrrc45 | -0,1740 | 3,280E-02 | 11 | protein_coding | leucine rich repeat containing 45 [Source:MGI Symbol;Acc:MGI:2387183] |
| Slc7a7 | -0,4452 | 3,280E-02 | 14 | protein_coding | solute carrier family 7 (cationic amino acid transporter, y+ system), member 7 [Source:MGI Symbol;Acc:MGI:1337120] |
| Gm9824 | -0,8502 | 3,290E-02 | 10 | processed_pseudogene | predicted pseudogene 9824 [Source:MGI Symbol;Acc:MGI:3641964] |
| Gm43174 | 1,2084 | 3,297E-02 | 5 | TEC | predicted gene 43174 [Source:MGI Symbol;Acc:MGI:5663311] |
| 4933431G14Rik | 1,8522 | 3,300E-02 | 6 | antisense | RIKEN cDNA 4933431G14 gene [Source:MGI Symbol;Acc:MGI:1918515] |
| Gm17833 | -2,8311 | 3,301E-02 | 5 | processed_pseudogene | predicted gene, 17833 [Source:MGI Symbol;Acc:MGI:5010018] |
| Zfp943 | 0,2554 | 3,302E-02 | 17 | protein_coding | zinc finger prtoein 943 [Source:MGI Symbol;Acc:MGI:1921920] |
| Gm21844 | -2,2560 | 3,320E-02 | 19 | antisense | predicted gene, 21844 [Source:MGI Symbol;Acc:MGI:5434008] |
| Gm10475 | 3,4327 | 3,335E-02 | 5 | antisense | predicted gene 10475 [Source:MGI Symbol;Acc:MGI:3642283] |
| Cxcl13 | -4,0383 | 3,343E-02 | 5 | protein_coding | chemokine (C-X-C motif) ligand 13 [Source:MGI Symbol;Acc:MGI:1888499] |
| Gm33543 | -1,3273 | 3,355E-02 | 10 | protein_coding | predicted gene, 33543 [Source:MGI Symbol;Acc:MGI:5592702] |
| Sema3c | -0,2254 | 3,355E-02 | 5 | protein_coding | sema domain, immunoglobulin domain (Ig), short basic domain, secreted, (semaphorin) 3C [Source:MGI Symbol;Acc:MGI:107557] |
| Vps13a | 0,1894 | 3,361E-02 | 19 | protein_coding | vacuolar protein sorting 13A [Source:MGI Symbol;Acc:MGI:2444304] |
| C230035I16Rik | 0,9899 | 3,371E-02 | 13 | lincRNA | RIKEN cDNA C230035I16 gene [Source:MGI Symbol;Acc:MGI:2444850] |
| Cdon | 0,1782 | 3,380E-02 | 9 | protein_coding | cell adhesion molecule-related/down-regulated by oncogenes [Source:MGI Symbol;Acc:MGI:1926387] |
| Gm14005 | -2,3052 | 3,382E-02 | 2 | lincRNA | predicted gene 14005 [Source:MGI Symbol;Acc:MGI:3652191] |
| 2810459M11Rik | 0,1830 | 3,388E-02 | 1 | protein_coding | RIKEN cDNA 2810459M11 gene [Source:MGI Symbol;Acc:MGI:1920042] |
| Baat | 2,1405 | 3,389E-02 | 4 | protein_coding | bile acid-Coenzyme A: amino acid N-acyltransferase [Source:MGI Symbol;Acc:MGI:106642] |
| Slfn9 | -1,2556 | 3,392E-02 | 11 | protein_coding | schlafen 9 [Source:MGI Symbol;Acc:MGI:2445121] |
| Dgkb | 0,1816 | 3,395E-02 | 12 | protein_coding | diacylglycerol kinase, beta [Source:MGI Symbol;Acc:MGI:2442474] |

| | | | | | |
|---|---|---|---|---|---|
| Map2k2 | -0,1659 | 3,402E-02 | 10 | protein_coding | mitogen-activated protein kinase kinase 2 [Source:MGI Symbol;Acc:MGI:1346867] |
| Gm20655 | 1,2957 | 3,407E-02 | 10 | antisense | predicted gene 20655 [Source:MGI Symbol;Acc:MGI:5313102] |
| Chm | 0,2923 | 3,410E-02 | X | protein_coding | choroidermia (RAB escort protein 1) [Source:MGI Symbol;Acc:MGI:892979] |
| Gm9484 | -1,3911 | 3,411E-02 | 5 | lincRNA | predicted gene 9484 [Source:MGI Symbol;Acc:MGI:3779894] |
| Gm9825 | 1,3061 | 3,411E-02 | 6 | processed_pseudogene | predicted gene 9825 [Source:MGI Symbol;Acc:MGI:3708729] |
| Cnppd1 | -0,1866 | 3,422E-02 | 1 | protein_coding | cyclin Pas1/PHO80 domain containing 1 [Source:MGI Symbol;Acc:MGI:1916421] |
| Rab31 | -0,1696 | 3,428E-02 | 17 | protein_coding | RAB31, member RAS oncogene family [Source:MGI Symbol;Acc:MGI:1914603] |
| Gm13905 | 2,5136 | 3,433E-02 | 2 | lincRNA | predicted gene 13905 [Source:MGI Symbol;Acc:MGI:3651198] |
| Tcea2 | -0,1879 | 3,433E-02 | 2 | protein_coding | transcription elongation factor A (SII), 2 [Source:MGI Symbol;Acc:MGI:107368] |
| Gssos1 | 3,2652 | 3,434E-02 | 2 | antisense | glutathione synthase, opposite strand 1 [Source:MGI Symbol;Acc:MGI:1915623] |
| Ksr1 | -0,1330 | 3,440E-02 | 11 | protein_coding | kinase suppressor of ras 1 [Source:MGI Symbol;Acc:MGI:105051] |
| mt-Tv | 1,2723 | 3,441E-02 | MT | Mt_tRNA | mitochondrially encoded tRNA valine [Source:MGI Symbol;Acc:MGI:102472] |
| Gm36423 | 2,3003 | 3,443E-02 | 13 | antisense | predicted gene, 36423 [Source:MGI Symbol;Acc:MGI:5595582] |
| Ints12 | 0,2770 | 3,448E-02 | 3 | protein_coding | integrator complex subunit 12 [Source:MGI Symbol;Acc:MGI:1919043] |
| Cela1 | -0,5771 | 3,450E-02 | 15 | protein_coding | chymotrypsin-like elastase family, member 1 [Source:MGI Symbol;Acc:MGI:95314] |
| Gm26613 | -1,5107 | 3,459E-02 | 11 | lincRNA | predicted gene, 26613 [Source:MGI Symbol;Acc:MGI:5477107] |
| Pcm1 | 0,2068 | 3,460E-02 | 8 | protein_coding | pericentriolar material 1 [Source:MGI Symbol;Acc:MGI:1277958] |
| Taok2 | -0,1157 | 3,461E-02 | 7 | protein_coding | TAO kinase 2 [Source:MGI Symbol;Acc:MGI:1915919] |
| Fastkd2 | -0,2514 | 3,464E-02 | 1 | protein_coding | FAST kinase domains 2 [Source:MGI Symbol;Acc:MGI:1922869] |
| Lrch2 | 0,3174 | 3,476E-02 | X | protein_coding | leucine-rich repeats and calponin homology (CH) domain containing 2 [Source:MGI Symbol;Acc:MGI:2147870] |
| Gm21284 | 2,0475 | 3,485E-02 | 6 | antisense | predicted gene, 21284 [Source:MGI Symbol;Acc:MGI:5434639] |
| Gm4956 | 3,2909 | 3,495E-02 | 1 | transcribed_unprocessed_pseudogene | predicted gene 4956 [Source:MGI Symbol;Acc:MGI:3647976] |
| Dscaml1 | -0,1889 | 3,498E-02 | 9 | protein_coding | DS cell adhesion molecule like 1 [Source:MGI Symbol;Acc:MGI:2150309] |
| Gm25716 | 3,7641 | 3,502E-02 | 14 | snRNA | predicted gene, 25716 [Source:MGI Symbol;Acc:MGI:5455493] |
| Cldn9 | -1,5232 | 3,504E-02 | 17 | protein_coding | claudin 9 [Source:MGI Symbol;Acc:MGI:1913100] |
| Dlgap3 | -0,1684 | 3,507E-02 | 4 | protein_coding | DLG associated protein 3 [Source:MGI Symbol;Acc:MGI:3039563] |
| Zfp354c | 0,1894 | 3,508E-02 | 11 | protein_coding | zinc finger protein 354C [Source:MGI Symbol;Acc:MGI:1353621] |

| | | | | | |
|---|---|---|---|---|---|
| Gm42972 | -1,2319 | 3,512E-02 | 3 | TEC | predicted gene 42972 [Source:MGI Symbol;Acc:MGI:5663109] |
| Pprc1 | 0,1643 | 3,518E-02 | 19 | protein_coding | peroxisome proliferative activated receptor, gamma, coactivator-related 1 [Source:MGI Symbol;Acc:MGI:2385096] |
| Srp54c | 0,2205 | 3,521E-02 | 12 | protein_coding | signal recognition particle 54C [Source:MGI Symbol;Acc:MGI:3714359] |
| Entpd7 | 0,2485 | 3,533E-02 | 19 | protein_coding | ectonucleoside triphosphate diphosphohydrolase 7 [Source:MGI Symbol;Acc:MGI:2135885] |
| Arhgap39 | -0,1480 | 3,535E-02 | 15 | protein_coding | Rho GTPase activating protein 39 [Source:MGI Symbol;Acc:MGI:107858] |
| Gm44093 | -3,1550 | 3,539E-02 | 6 | TEC | predicted gene, 44093 [Source:MGI Symbol;Acc:MGI:5690485] |
| Gm45579 | 3,8132 | 3,549E-02 | 4 | lincRNA | predicted gene 45579 [Source:MGI Symbol;Acc:MGI:5791415] |
| Fbxl18 | -0,1499 | 3,552E-02 | 5 | protein_coding | F-box and leucine-rich repeat protein 18 [Source:MGI Symbol;Acc:MGI:2444450] |
| Usp18 | 0,7684 | 3,557E-02 | 6 | protein_coding | ubiquitin specific peptidase 18 [Source:MGI Symbol;Acc:MGI:1344364] |
| Pfn1 | -0,1714 | 3,558E-02 | 11 | protein_coding | profilin 1 [Source:MGI Symbol;Acc:MGI:97549] |
| Erp27 | -2,2750 | 3,564E-02 | 6 | protein_coding | endoplasmic reticulum protein 27 [Source:MGI Symbol;Acc:MGI:1916437] |
| Gm26575 | 0,9394 | 3,569E-02 | 18 | sense_intronic | predicted gene, 26575 [Source:MGI Symbol;Acc:MGI:5477069] |
| Gm44911 | -0,6377 | 3,573E-02 | 8 | processed_pseudogene | predicted gene 44911 [Source:MGI Symbol;Acc:MGI:5753487] |
| Diaph2 | 0,2963 | 3,573E-02 | X | protein_coding | diaphanous related formin 2 [Source:MGI Symbol;Acc:MGI:1858500] |
| Irx2 | -3,0189 | 3,590E-02 | 13 | protein_coding | Iroquois homeobox 2 [Source:MGI Symbol;Acc:MGI:1197526] |
| Nln | 0,2027 | 3,604E-02 | 13 | protein_coding | neurolysin (metallopeptidase M3 family) [Source:MGI Symbol;Acc:MGI:1923055] |
| Bcl2l1 | -0,1503 | 3,608E-02 | 2 | protein_coding | BCL2-like 1 [Source:MGI Symbol;Acc:MGI:88139] |
| Dnm2 | -0,1502 | 3,608E-02 | 9 | protein_coding | dynamin 2 [Source:MGI Symbol;Acc:MGI:109547] |
| Gm48949 | -1,9708 | 3,614E-02 | 14 | sense_intronic | predicted gene, 48949 [Source:MGI Symbol;Acc:MGI:6118274] |
| Aatf | 0,2765 | 3,614E-02 | 11 | protein_coding | apoptosis antagonizing transcription factor [Source:MGI Symbol;Acc:MGI:1929608] |
| Gm15443 | 3,5521 | 3,618E-02 | 1 | processed_pseudogene | predicted gene 15443 [Source:MGI Symbol;Acc:MGI:3642525] |
| Zfp945 | 0,3116 | 3,618E-02 | 17 | protein_coding | zinc finger protein 945 [Source:MGI Symbol;Acc:MGI:2445132] |
| Myom1 | 0,4605 | 3,641E-02 | 17 | protein_coding | myomesin 1 [Source:MGI Symbol;Acc:MGI:1341430] |
| Pdzd2 | -0,1903 | 3,650E-02 | 15 | protein_coding | PDZ domain containing 2 [Source:MGI Symbol;Acc:MGI:1922394] |
| Rpl15-ps2 | 1,7552 | 3,650E-02 | 9 | processed_pseudogene | ribosomal protein L15, pseudogene 2 [Source:MGI Symbol;Acc:MGI:3648255] |
| Fcrl1 | 1,1294 | 3,652E-02 | 3 | protein_coding | Fc receptor-like 1 [Source:MGI Symbol;Acc:MGI:2442862] |
| Gm48382 | 1,6301 | 3,660E-02 | 12 | TEC | predicted gene, 48382 [Source:MGI Symbol;Acc:MGI:6097861] |

| | | | | | |
|---|---|---|---|---|---|
| Tmem87b | 0,1712 | 3,660E-02 | 2 | protein_coding | transmembrane protein 87B [Source:MGI Symbol;Acc:MGI:1919727] |
| Cdk8 | 0,1492 | 3,660E-02 | 5 | protein_coding | cyclin-dependent kinase 8 [Source:MGI Symbol;Acc:MGI:1196224] |
| Rab40c | -0,1740 | 3,661E-02 | 17 | protein_coding | Rab40C, member RAS oncogene family [Source:MGI Symbol;Acc:MGI:2183454] |
| Gm44068 | 2,0802 | 3,664E-02 | 6 | TEC | predicted gene, 44068 [Source:MGI Symbol;Acc:MGI:5690460] |
| Slc39a12 | 0,2220 | 3,664E-02 | 2 | protein_coding | solute carrier family 39 (zinc transporter), member 12 [Source:MGI Symbol;Acc:MGI:2139274] |
| Sorbs2os | 0,5981 | 3,667E-02 | 8 | antisense | sorbin and SH3 domain containing 2, opposite strand [Source:MGI Symbol;Acc:MGI:2443013] |
| Fam208b | 0,2423 | 3,668E-02 | 13 | protein_coding | family with sequence similarity 208, member B [Source:MGI Symbol;Acc:MGI:2145274] |
| Map3k10 | -0,1512 | 3,674E-02 | 7 | protein_coding | mitogen-activated protein kinase kinase kinase 10 [Source:MGI Symbol;Acc:MGI:1346879] |
| Ptgs2os | 0,9884 | 3,676E-02 | 1 | processed_transcript | prostaglandin-endoperoxide synthase 2, opposite strand [Source:MGI Symbol;Acc:MGI:2443180] |
| Gm20125 | 1,5877 | 3,676E-02 | 10 | processed_transcript | predicted gene, 20125 [Source:MGI Symbol;Acc:MGI:5012310] |
| Gm20407 | 2,9886 | 3,678E-02 | 19 | processed_transcript | predicted gene 20407 [Source:MGI Symbol;Acc:MGI:5141872] |
| Hmgxb3 | -0,1489 | 3,690E-02 | 18 | protein_coding | HMG box domain containing 3 [Source:MGI Symbol;Acc:MGI:2441817] |
| Galnt1 | 0,1853 | 3,696E-02 | 18 | protein_coding | polypeptide N-acetylgalactosaminyltransferase 1 [Source:MGI Symbol;Acc:MGI:894693] |
| Git1 | -0,1460 | 3,709E-02 | 11 | protein_coding | G protein-coupled receptor kinase-interactor 1 [Source:MGI Symbol;Acc:MGI:1927140] |
| Gpd1 | -0,1687 | 3,710E-02 | 15 | protein_coding | glycerol-3-phosphate dehydrogenase 1 (soluble) [Source:MGI Symbol;Acc:MGI:95679] |
| Adamts1 | 0,2681 | 3,712E-02 | 16 | protein_coding | a disintegrin-like and metallopeptidase (reprolysin type) with thrombospondin type 1 motif, 1 [Source:MGI Symbol;Acc:MGI:109249] |
| Nudt22 | -0,4257 | 3,713E-02 | 19 | protein_coding | nudix (nucleoside diphosphate linked moiety X)-type motif 22 [Source:MGI Symbol;Acc:MGI:1915573] |
| Rfng | -0,1511 | 3,713E-02 | 11 | protein_coding | RFNG O-fucosylpeptide 3-beta-N-acetylglucosaminyltransferase [Source:MGI Symbol;Acc:MGI:894275] |
| Gm20616 | 3,8123 | 3,716E-02 | 19 | processed_transcript | predicted gene 20616 [Source:MGI Symbol;Acc:MGI:5313063] |
| Gm48508 | -0,9197 | 3,721E-02 | 12 | lincRNA | predicted gene, 48508 [Source:MGI Symbol;Acc:MGI:6098039] |
| A330074H02Rik | 0,7740 | 3,738E-02 | 7 | TEC | RIKEN cDNA A330074H02 gene [Source:MGI Symbol;Acc:MGI:3028065] |
| Glrp1 | 2,8875 | 3,743E-02 | 1 | protein_coding | glutamine repeat protein 1 [Source:MGI Symbol;Acc:MGI:108038] |
| Gm44007 | -3,3070 | 3,752E-02 | 6 | TEC | predicted gene, 44007 [Source:MGI Symbol;Acc:MGI:5690399] |
| Gm44813 | 2,5422 | 3,759E-02 | 7 | lincRNA | predicted gene 44813 [Source:MGI Symbol;Acc:MGI:5753389] |
| Gm43137 | -2,8776 | 3,760E-02 | 5 | antisense | predicted gene 43137 [Source:MGI Symbol;Acc:MGI:5663274] |
| Zfp445 | 0,1468 | 3,770E-02 | 9 | protein_coding | zinc finger protein 445 [Source:MGI Symbol;Acc:MGI:2143340] |
| Alg12 | -0,2289 | 3,786E-02 | 15 | protein_coding | asparagine-linked glycosylation 12 (alpha-1,6-mannosyltransferase) [Source:MGI Symbol;Acc:MGI:2385025] |

| | | | | | |
|---|---|---|---|---|---|
| St8sia6 | 0,4140 | 3,788E-02 | 2 | protein_coding | ST8 alpha-N-acetyl-neuraminide alpha-2,8-sialyltransferase 6 [Source:MGI Symbol;Acc:MGI:2386797] |
| Gtpbp3 | -0,1859 | 3,789E-02 | 8 | protein_coding | GTP binding protein 3 [Source:MGI Symbol;Acc:MGI:1917609] |
| Pitpnm2 | -0,1164 | 3,791E-02 | 5 | protein_coding | phosphatidylinositol transfer protein, membrane-associated 2 [Source:MGI Symbol;Acc:MGI:1336192] |
| C130013H08Rik | 0,7299 | 3,797E-02 | 3 | antisense | RIKEN cDNA C130013H08 gene [Source:MGI Symbol;Acc:MGI:3697343] |
| Gm29797 | -0,8695 | 3,800E-02 | 2 | protein_coding | predicted gene, 29797 [Source:MGI Symbol;Acc:MGI:5588956] |
| 1700001G11Rik | 1,5813 | 3,810E-02 | 14 | processed_transcript | RIKEN cDNA 1700001G11 gene [Source:MGI Symbol;Acc:MGI:1916553] |
| Wdr24 | -0,1720 | 3,823E-02 | 17 | protein_coding | WD repeat domain 24 [Source:MGI Symbol;Acc:MGI:2446285] |
| Fam243 | -3,9609 | 3,831E-02 | 16 | protein_coding | family with sequence similarity 243 [Source:MGI Symbol;Acc:MGI:1922578] |
| Gm10566 | 2,5491 | 3,846E-02 | 1 | processed_pseudogene | predicted gene 10566 [Source:MGI Symbol;Acc:MGI:3642220] |
| Gm37941 | 1,0729 | 3,848E-02 | 9 | TEC | predicted gene, 37941 [Source:MGI Symbol;Acc:MGI:5611169] |
| Sltm | 0,1728 | 3,855E-02 | 9 | protein_coding | SAFB-like, transcription modulator [Source:MGI Symbol;Acc:MGI:1913910] |
| Ddx58 | 0,3835 | 3,861E-02 | 4 | protein_coding | DEAD (Asp-Glu-Ala-Asp) box polypeptide 58 [Source:MGI Symbol;Acc:MGI:2442858] |
| Gm5089 | 2,7193 | 3,862E-02 | 14 | protein_coding | predicted gene 5089 [Source:MGI Symbol;Acc:MGI:3644731] |
| Tesc | -0,2033 | 3,868E-02 | 5 | protein_coding | tescalcin [Source:MGI Symbol;Acc:MGI:1930803] |
| Tgtp2 | 0,8650 | 3,878E-02 | 11 | protein_coding | T cell specific GTPase 2 [Source:MGI Symbol;Acc:MGI:3710083] |
| Gm3716 | 2,2552 | 3,880E-02 | 5 | lincRNA | predicted gene 3716 [Source:MGI Symbol;Acc:MGI:3781892] |
| F2rl3 | 0,6580 | 3,885E-02 | 8 | protein_coding | coagulation factor II (thrombin) receptor-like 3 [Source:MGI Symbol;Acc:MGI:1298207] |
| Timm22 | -0,1853 | 3,890E-02 | 11 | protein_coding | translocase of inner mitochondrial membrane 22 [Source:MGI Symbol;Acc:MGI:1929742] |
| Gm33045 | -1,3975 | 3,890E-02 | 13 | lincRNA | predicted gene, 33045 [Source:MGI Symbol;Acc:MGI:5592204] |
| Nelfb | -0,1702 | 3,895E-02 | 2 | protein_coding | negative elongation factor complex member B [Source:MGI Symbol;Acc:MGI:1931035] |
| Agtr1b | 2,3666 | 3,899E-02 | 3 | protein_coding | angiotensin II receptor, type 1b [Source:MGI Symbol;Acc:MGI:87965] |
| Zmiz1os1 | 0,7058 | 3,899E-02 | 14 | antisense | Zmiz1 opposite strand 1 [Source:MGI Symbol;Acc:MGI:3041228] |
| Zfp524 | -0,6102 | 3,901E-02 | 7 | protein_coding | zinc finger protein 524 [Source:MGI Symbol;Acc:MGI:1916740] |
| A930015D03Rik | -2,4604 | 3,904E-02 | 17 | antisense | RIKEN cDNA A930015D03 gene [Source:MGI Symbol;Acc:MGI:1925060] |
| Ppil6 | -0,5326 | 3,907E-02 | 10 | protein_coding | peptidylprolyl isomerase (cyclophilin)-like 6 [Source:MGI Symbol;Acc:MGI:1920325] |
| Rnd2 | -0,1590 | 3,907E-02 | 11 | protein_coding | Rho family GTPase 2 [Source:MGI Symbol;Acc:MGI:1338755] |
| Gm49493 | 1,1752 | 3,909E-02 | 7 | TEC | predicted gene, 49493 [Source:MGI Symbol;Acc:MGI:6155171] |

| | | | | | |
|---|---|---|---|---|---|
| Gm23954 | -2,3705 | 3,912E-02 | 18 | misc_RNA | predicted gene, 23954 [Source:MGI Symbol;Acc:MGI:5453731] |
| Zfp932 | 0,2341 | 3,915E-02 | 5 | protein_coding | zinc finger protein 932 [Source:MGI Symbol;Acc:MGI:1916754] |
| Bbip1 | 0,1847 | 3,924E-02 | 19 | protein_coding | BBSome interacting protein 1 [Source:MGI Symbol;Acc:MGI:1913610] |
| Gm26753 | 3,8104 | 3,924E-02 | 17 | lincRNA | predicted gene, 26753 [Source:MGI Symbol;Acc:MGI:5477247] |
| 4930558K02Rik | 3,7658 | 3,926E-02 | 1 | protein_coding | RIKEN cDNA 4930558K02 gene [Source:MGI Symbol;Acc:MGI:1922618] |
| Gfra4 | -0,1864 | 3,932E-02 | 2 | protein_coding | glial cell line derived neurotrophic factor family receptor alpha 4 [Source:MGI Symbol;Acc:MGI:1341873] |
| Zfp955b | 0,3290 | 3,938E-02 | 17 | protein_coding | zinc finger protein 955B [Source:MGI Symbol;Acc:MGI:4834573] |
| Mir22hg | 0,2596 | 3,938E-02 | 11 | processed_transcript | Mir22 host gene (non-protein coding) [Source:MGI Symbol;Acc:MGI:1914348] |
| Cavin2 | 0,3485 | 3,940E-02 | 1 | protein_coding | caveolae associated 2 [Source:MGI Symbol;Acc:MGI:99513] |
| Npas3 | 0,2711 | 3,945E-02 | 12 | protein_coding | neuronal PAS domain protein 3 [Source:MGI Symbol;Acc:MGI:1351610] |
| Treml4 | 2,4196 | 3,948E-02 | 17 | protein_coding | triggering receptor expressed on myeloid cells-like 4 [Source:MGI Symbol;Acc:MGI:1923239] |
| Ttll10 | -2,1416 | 3,949E-02 | 4 | protein_coding | tubulin tyrosine ligase-like family, member 10 [Source:MGI Symbol;Acc:MGI:1921855] |
| Snph | -0,1210 | 3,973E-02 | 2 | protein_coding | syntaphilin [Source:MGI Symbol;Acc:MGI:2139270] |
| Plppr2 | -0,1472 | 3,975E-02 | 9 | protein_coding | phospholipid phosphatase related 2 [Source:MGI Symbol;Acc:MGI:2384575] |
| Gm13246 | 3,4370 | 3,980E-02 | 4 | processed_pseudogene | predicted gene 13246 [Source:MGI Symbol;Acc:MGI:3649657] |
| Smarca5 | 0,2338 | 3,991E-02 | 8 | protein_coding | SWI/SNF related, matrix associated, actin dependent regulator of chromatin, subfamily a, member 5 [Source:MGI Symbol;Acc:MGI:1935[illegible] |
| Ccdc39 | 0,2631 | 3,995E-02 | 3 | protein_coding | coiled-coil domain containing 39 [Source:MGI Symbol;Acc:MGI:1289263] |
| Alkbh1 | 0,2545 | 3,997E-02 | 12 | protein_coding | alkB homolog 1, histone H2A dioxygenase [Source:MGI Symbol;Acc:MGI:2384034] |
| Tomm5 | -0,2023 | 4,008E-02 | 4 | protein_coding | translocase of outer mitochondrial membrane 5 [Source:MGI Symbol;Acc:MGI:1915762] |
| Sertad3 | -0,5591 | 4,018E-02 | 7 | protein_coding | SERTA domain containing 3 [Source:MGI Symbol;Acc:MGI:2180697] |
| Zbtb42 | -0,6633 | 4,032E-02 | 12 | protein_coding | zinc finger and BTB domain containing 42 [Source:MGI Symbol;Acc:MGI:3644133] |
| Cnih2 | -0,1616 | 4,044E-02 | 19 | protein_coding | cornichon family AMPA receptor auxiliary protein 2 [Source:MGI Symbol;Acc:MGI:1277225] |
| Wdr91 | -0,1686 | 4,052E-02 | 6 | protein_coding | WD repeat domain 91 [Source:MGI Symbol;Acc:MGI:2141558] |
| Gm48632 | 2,8073 | 4,053E-02 | 12 | TEC | predicted gene, 48632 [Source:MGI Symbol;Acc:MGI:6098236] |
| 4930502E09Rik | 3,7063 | 4,064E-02 | 11 | processed_transcript | RIKEN cDNA 4930502E09 gene [Source:MGI Symbol;Acc:MGI:1922245] |
| Cfap36 | 0,1565 | 4,068E-02 | 11 | protein_coding | cilia and flagella associated protein 36 [Source:MGI Symbol;Acc:MGI:1913994] |
| Gm49214 | -3,6421 | 4,069E-02 | 15 | processed_pseudogene | predicted gene, 49214 [Source:MGI Symbol;Acc:MGI:6118668] |

| | | | | | |
|---|---|---|---|---|---|
| Gm26902 | 0,7465 | 4,076E-02 | 19 | lincRNA | predicted gene, 26902 [Source:MGI Symbol;Acc:MGI:5477396] |
| Gm38387 | 0,7411 | 4,084E-02 | 1 | TEC | predicted gene, 38387 [Source:MGI Symbol;Acc:MGI:5613622] |
| Ect2l | 0,7867 | 4,095E-02 | 10 | protein_coding | epithelial cell transforming sequence 2 oncogene-like [Source:MGI Symbol;Acc:MGI:3641723] |
| Gm42693 | -1,3917 | 4,103E-02 | 3 | TEC | predicted gene 42693 [Source:MGI Symbol;Acc:MGI:5662830] |
| Psmb8 | 0,4370 | 4,106E-02 | 17 | protein_coding | proteasome (prosome, macropain) subunit, beta type 8 (large multifunctional peptidase 7) [Source:MGI Symbol;Acc:MGI:1346527] |
| Dclk2 | -0,1296 | 4,108E-02 | 3 | protein_coding | doublecortin-like kinase 2 [Source:MGI Symbol;Acc:MGI:1918012] |
| Tmem159 | 0,2374 | 4,109E-02 | 7 | protein_coding | transmembrane protein 159 [Source:MGI Symbol;Acc:MGI:1925752] |
| Gm44641 | 2,7342 | 4,115E-02 | 7 | processed_transcript | predicted gene 44641 [Source:MGI Symbol;Acc:MGI:5753217] |
| Il20rb | -0,3905 | 4,117E-02 | 9 | protein_coding | interleukin 20 receptor beta [Source:MGI Symbol;Acc:MGI:2143266] |
| Mkl1 | -0,1565 | 4,120E-02 | 15 | protein_coding | MKL (megakaryoblastic leukemia)/myocardin-like 1 [Source:MGI Symbol;Acc:MGI:2384495] |
| Cklf | 0,4238 | 4,129E-02 | 8 | protein_coding | chemokine-like factor [Source:MGI Symbol;Acc:MGI:1922708] |
| Cdip1 | -0,1005 | 4,130E-02 | 16 | protein_coding | cell death inducing Trp53 target 1 [Source:MGI Symbol;Acc:MGI:1913876] |
| Gm9821 | -0,5690 | 4,131E-02 | 2 | protein_coding | predicted gene 9821 [Source:MGI Symbol;Acc:MGI:3704208] |
| Gm18336 | -2,9948 | 4,132E-02 | X | protein_coding | predicted gene, 18336 [Source:MGI Symbol;Acc:MGI:5010521] |
| Naxe | -0,1726 | 4,134E-02 | 3 | protein_coding | NAD(P)HX epimerase [Source:MGI Symbol;Acc:MGI:2180167] |
| Zer1 | -0,1175 | 4,142E-02 | 2 | protein_coding | zyg-11 related, cell cycle regulator [Source:MGI Symbol;Acc:MGI:2442511] |
| Nek3 | 0,5844 | 4,146E-02 | 8 | protein_coding | NIMA (never in mitosis gene a)-related expressed kinase 3 [Source:MGI Symbol;Acc:MGI:1344371] |
| Lsm6 | 0,1845 | 4,157E-02 | 8 | protein_coding | LSM6 homolog, U6 small nuclear RNA and mRNA degradation associated [Source:MGI Symbol;Acc:MGI:1925901] |
| Dctn2 | -0,1604 | 4,172E-02 | 10 | protein_coding | dynactin 2 [Source:MGI Symbol;Acc:MGI:107733] |
| Hspbp1 | -0,1672 | 4,175E-02 | 7 | protein_coding | HSPA (heat shock 70kDa) binding protein, cytoplasmic cochaperone 1 [Source:MGI Symbol;Acc:MGI:1913495] |
| Chrna6 | -2,3790 | 4,180E-02 | 8 | protein_coding | cholinergic receptor, nicotinic, alpha polypeptide 6 [Source:MGI Symbol;Acc:MGI:106213] |
| Gm8369 | -2,1970 | 4,186E-02 | 19 | protein_coding | predicted gene 8369 [Source:MGI Symbol;Acc:MGI:3645380] |
| Sp3 | 0,2098 | 4,187E-02 | 2 | protein_coding | trans-acting transcription factor 3 [Source:MGI Symbol;Acc:MGI:1277166] |
| Fmnl2 | 0,1864 | 4,190E-02 | 2 | protein_coding | formin-like 2 [Source:MGI Symbol;Acc:MGI:1918659] |
| Limk1 | -0,1726 | 4,196E-02 | 5 | protein_coding | LIM-domain containing, protein kinase [Source:MGI Symbol;Acc:MGI:104572] |
| Gm18194 | -0,7428 | 4,196E-02 | 7 | transcribed_processed_pseudogene | predicted gene, 18194 [Source:MGI Symbol;Acc:MGI:5010379] |
| Ino80b | -0,2561 | 4,200E-02 | 6 | protein_coding | INO80 complex subunit B [Source:MGI Symbol;Acc:MGI:1917270] |

| | | | | | |
|---|---|---|---|---|---|
| Ifih1 | 0,4836 | 4,201E-02 | 2 | protein_coding | interferon induced with helicase C domain 1 [Source:MGI Symbol;Acc:MGI:1918836] |
| Dnttip2 | 0,1993 | 4,203E-02 | 3 | protein_coding | deoxynucleotidyltransferase, terminal, interacting protein 2 [Source:MGI Symbol;Acc:MGI:1923173] |
| Ccdc142os | 1,2744 | 4,205E-02 | 6 | antisense | coiled-coil domain containing 142, opposite strand [Source:MGI Symbol;Acc:MGI:3783052] |
| Zfp111 | 0,3235 | 4,209E-02 | 7 | protein_coding | zinc finger protein 111 [Source:MGI Symbol;Acc:MGI:1929114] |
| Cd79b | -1,5770 | 4,209E-02 | 11 | protein_coding | CD79B antigen [Source:MGI Symbol;Acc:MGI:96431] |
| Ublcp1 | 0,1395 | 4,215E-02 | 11 | protein_coding | ubiquitin-like domain containing CTD phosphatase 1 [Source:MGI Symbol;Acc:MGI:1933105] |
| Tenm2 | 0,1653 | 4,216E-02 | 11 | protein_coding | teneurin transmembrane protein 2 [Source:MGI Symbol;Acc:MGI:1345184] |
| Cox4i2 | 0,5294 | 4,234E-02 | 2 | protein_coding | cytochrome c oxidase subunit 4I2 [Source:MGI Symbol;Acc:MGI:2135755] |
| Gzmm | -0,5495 | 4,234E-02 | 10 | protein_coding | granzyme M (lymphocyte met-ase 1) [Source:MGI Symbol;Acc:MGI:99549] |
| Zfp580 | -0,1999 | 4,248E-02 | 7 | protein_coding | zinc finger protein 580 [Source:MGI Symbol;Acc:MGI:1916242] |
| Rbp3 | 1,8219 | 4,252E-02 | 14 | protein_coding | retinol binding protein 3, interstitial [Source:MGI Symbol;Acc:MGI:97878] |
| Borcs6 | -0,2080 | 4,253E-02 | 11 | protein_coding | BLOC-1 related complex subunit 6 [Source:MGI Symbol;Acc:MGI:1919173] |
| Src | -0,1422 | 4,261E-02 | 2 | protein_coding | Rous sarcoma oncogene [Source:MGI Symbol;Acc:MGI:98397] |
| Hepacam | -0,1363 | 4,262E-02 | 9 | protein_coding | hepatocyte cell adhesion molecule [Source:MGI Symbol;Acc:MGI:1920177] |
| Ttc9b | -0,2000 | 4,265E-02 | 7 | protein_coding | tetratricopeptide repeat domain 9B [Source:MGI Symbol;Acc:MGI:1920282] |
| Tnk2 | -0,1474 | 4,286E-02 | 16 | protein_coding | tyrosine kinase, non-receptor, 2 [Source:MGI Symbol;Acc:MGI:1858308] |
| Ube2c | 1,9939 | 4,287E-02 | 2 | protein_coding | ubiquitin-conjugating enzyme E2C [Source:MGI Symbol;Acc:MGI:1915862] |
| Gm37494 | 0,2559 | 4,287E-02 | 7 | lincRNA | predicted gene, 37494 [Source:MGI Symbol;Acc:MGI:5610722] |
| Gm16061 | 3,0545 | 4,288E-02 | 5 | processed_pseudogene | predicted gene 16061 [Source:MGI Symbol;Acc:MGI:3801920] |
| Tmem121b | -0,1420 | 4,290E-02 | 6 | protein_coding | transmembrane protein 121B [Source:MGI Symbol;Acc:MGI:2136977] |
| Coro7 | -0,1491 | 4,306E-02 | 16 | protein_coding | coronin 7 [Source:MGI Symbol;Acc:MGI:1926135] |
| Gm7072 | 0,3537 | 4,310E-02 | 17 | protein_coding | predicted gene 7072 [Source:NCBI gene;Acc:631624] |
| Tbc1d13 | -0,1546 | 4,316E-02 | 2 | protein_coding | TBC1 domain family, member 13 [Source:MGI Symbol;Acc:MGI:2385326] |
| Psmb5 | -0,1642 | 4,325E-02 | 14 | protein_coding | proteasome (prosome, macropain) subunit, beta type 5 [Source:MGI Symbol;Acc:MGI:1194513] |
| Gm21887 | -0,6385 | 4,325E-02 | X | protein_coding | predicted gene, 21887 [Source:MGI Symbol;Acc:MGI:5434051] |
| Gm15387 | 0,5034 | 4,328E-02 | 15 | processed_pseudogene | predicted gene 15387 [Source:MGI Symbol;Acc:MGI:3705374] |
| Rps27a | -0,5485 | 4,331E-02 | 11 | protein_coding | ribosomal protein S27A [Source:MGI Symbol;Acc:MGI:1925544] |

| | | | | | |
|---|---|---|---|---|---|
| Ppa2 | 0,1422 | 4,337E-02 | 3 | protein_coding | pyrophosphatase (inorganic) 2 [Source:MGI Symbol;Acc:MGI:1922026] |
| Ggta1 | 0,3912 | 4,343E-02 | 2 | protein_coding | glycoprotein galactosyltransferase alpha 1, 3 [Source:MGI Symbol;Acc:MGI:95704] |
| Slc30a1 | 0,1865 | 4,353E-02 | 1 | protein_coding | solute carrier family 30 (zinc transporter), member 1 [Source:MGI Symbol;Acc:MGI:1345281] |
| Gm14036 | -2,2933 | 4,354E-02 | 2 | processed_pseudogene | predicted gene 14036 [Source:MGI Symbol;Acc:MGI:3649581] |
| Mb | 1,4228 | 4,359E-02 | 15 | protein_coding | myoglobin [Source:MGI Symbol;Acc:MGI:96922] |
| Mcrip1 | -0,1754 | 4,364E-02 | 11 | protein_coding | MAPK regulated corepressor interacting protein 1 [Source:MGI Symbol;Acc:MGI:2384752] |
| Gm5641 | 0,5566 | 4,375E-02 | 3 | transcribed_processed_pseudogene | predicted gene 5641 [Source:MGI Symbol;Acc:MGI:3645731] |
| 9630050E16Rik | -2,0643 | 4,376E-02 | 14 | TEC | RIKEN cDNA 9630050E16 gene [Source:MGI Symbol;Acc:MGI:2443518] |
| Zfp281 | 0,1792 | 4,380E-02 | 1 | protein_coding | zinc finger protein 281 [Source:MGI Symbol;Acc:MGI:3029290] |
| Hbb-bs | -0,3853 | 4,385E-02 | 7 | protein_coding | hemoglobin, beta adult s chain [Source:MGI Symbol;Acc:MGI:5474852] |
| Tmem181b-ps | 0,2070 | 4,387E-02 | 17 | transcribed_unprocessed_pseudogene | transmembrane protein 181B, pseudogene [Source:MGI Symbol;Acc:MGI:3779544] |
| Gm15895 | -3,3484 | 4,398E-02 | 8 | antisense | predicted gene 15895 [Source:MGI Symbol;Acc:MGI:3802071] |
| Ftl1 | -0,1669 | 4,401E-02 | 7 | protein_coding | ferritin light polypeptide 1 [Source:MGI Symbol;Acc:MGI:95589] |
| Asic2 | -0,1582 | 4,403E-02 | 11 | protein_coding | acid-sensing (proton-gated) ion channel 2 [Source:MGI Symbol;Acc:MGI:1100867] |
| Ubl7 | -0,1623 | 4,404E-02 | 9 | protein_coding | ubiquitin-like 7 (bone marrow stromal cell-derived) [Source:MGI Symbol;Acc:MGI:1916709] |
| Maff | 0,5452 | 4,420E-02 | 15 | protein_coding | v-maf musculoaponeurotic fibrosarcoma oncogene family, protein F (avian) [Source:MGI Symbol;Acc:MGI:96910] |
| Atxn7l3 | -0,1294 | 4,441E-02 | 11 | protein_coding | ataxin 7-like 3 [Source:MGI Symbol;Acc:MGI:3036270] |
| Pcna | 0,1966 | 4,448E-02 | 2 | protein_coding | proliferating cell nuclear antigen [Source:MGI Symbol;Acc:MGI:97503] |
| Brf2 | -0,2053 | 4,449E-02 | 8 | protein_coding | BRF2, RNA polymerase III transcription initiation factor 50kDa subunit [Source:MGI Symbol;Acc:MGI:1913903] |
| Gm24187 | 2,8577 | 4,455E-02 | 13 | miRNA | predicted gene, 24187 [Source:MGI Symbol;Acc:MGI:5453964] |
| Wnt2 | -0,5041 | 4,460E-02 | 6 | protein_coding | wingless-type MMTV integration site family, member 2 [Source:MGI Symbol;Acc:MGI:98954] |
| Sgsm2 | -0,1127 | 4,464E-02 | 11 | protein_coding | small G protein signaling modulator 2 [Source:MGI Symbol;Acc:MGI:2144695] |
| Gm37834 | -2,3460 | 4,467E-02 | 3 | TEC | predicted gene, 37834 [Source:MGI Symbol;Acc:MGI:5611062] |
| 1700011M02Rik | 3,8781 | 4,467E-02 | X | protein_coding | RIKEN cDNA 1700011M02 gene [Source:MGI Symbol;Acc:MGI:1922693] |
| E2f5 | 0,2706 | 4,471E-02 | 3 | protein_coding | E2F transcription factor 5 [Source:MGI Symbol;Acc:MGI:105091] |
| Plch2 | -0,1446 | 4,480E-02 | 4 | protein_coding | phospholipase C, eta 2 [Source:MGI Symbol;Acc:MGI:2443078] |
| Gm43573 | 3,8754 | 4,501E-02 | 3 | antisense | predicted gene 43573 [Source:MGI Symbol;Acc:MGI:5663710] |

| | | | | | |
|---|---|---|---|---|---|
| Nlrp6 | -0,7154 | 4,503E-02 | 7 | protein_coding | NLR family, pyrin domain containing 6 [Source:MGI Symbol;Acc:MGI:2141990] |
| Slc26a6 | -0,2555 | 4,505E-02 | 9 | protein_coding | solute carrier family 26, member 6 [Source:MGI Symbol;Acc:MGI:2159728] |
| Pcbd1 | -0,4641 | 4,506E-02 | 10 | protein_coding | pterin 4 alpha carbinolamine dehydratase/dimerization cofactor of hepatocyte nuclear factor 1 alpha (TCF1) 1 [Source:MGI Symbol;Acc:MGI:94873] |
| Svop | -0,1409 | 4,507E-02 | 5 | protein_coding | SV2 related protein [Source:MGI Symbol;Acc:MGI:1915916] |
| Tmem30b | -2,0227 | 4,507E-02 | 12 | protein_coding | transmembrane protein 30B [Source:MGI Symbol;Acc:MGI:2442082] |
| Gm6710 | 0,8795 | 4,524E-02 | 2 | protein_coding | predicted gene 6710 [Source:MGI Symbol;Acc:MGI:3779623] |
| Rps12-ps4 | 1,3535 | 4,526E-02 | 7 | processed_pseudogene | ribosomal protein S12, pseudogene 4 [Source:MGI Symbol;Acc:MGI:3780139] |
| Ddah2 | -0,3863 | 4,532E-02 | 17 | protein_coding | dimethylarginine dimethylaminohydrolase 2 [Source:MGI Symbol;Acc:MGI:1859016] |
| Dera | -0,5400 | 4,538E-02 | 6 | protein_coding | deoxyribose-phosphate aldolase (putative) [Source:MGI Symbol;Acc:MGI:1913762] |
| Snrpb | -0,1618 | 4,542E-02 | 2 | protein_coding | small nuclear ribonucleoprotein B [Source:MGI Symbol;Acc:MGI:98342] |
| Pcdhga5 | 0,3776 | 4,554E-02 | 18 | protein_coding | protocadherin gamma subfamily A, 5 [Source:MGI Symbol;Acc:MGI:1935217] |
| Gm42725 | 1,0001 | 4,555E-02 | 5 | TEC | predicted gene 42725 [Source:MGI Symbol;Acc:MGI:5662862] |
| Gm19589 | -3,9431 | 4,557E-02 | 1 | lincRNA | predicted gene, 19589 [Source:MGI Symbol;Acc:MGI:5011774] |
| Gm46364 | 3,3073 | 4,560E-02 | 12 | processed_pseudogene | predicted gene, 46364 [Source:MGI Symbol;Acc:MGI:5826001] |
| Adprh | -0,1522 | 4,569E-02 | 16 | protein_coding | ADP-ribosylarginine hydrolase [Source:MGI Symbol;Acc:MGI:1098234] |
| Gpr62 | -0,2330 | 4,570E-02 | 9 | protein_coding | G protein-coupled receptor 62 [Source:MGI Symbol;Acc:MGI:3525078] |
| Fntb | -0,1636 | 4,573E-02 | 12 | protein_coding | farnesyltransferase, CAAX box, beta [Source:MGI Symbol;Acc:MGI:1861305] |
| Dmtf1 | 0,1582 | 4,584E-02 | 5 | protein_coding | cyclin D binding myb-like transcription factor 1 [Source:MGI Symbol;Acc:MGI:1344415] |
| Tcap | 0,8394 | 4,585E-02 | 11 | protein_coding | titin-cap [Source:MGI Symbol;Acc:MGI:1330233] |
| Ssb | 0,2450 | 4,586E-02 | 2 | protein_coding | Sjogren syndrome antigen B [Source:MGI Symbol;Acc:MGI:98423] |
| Th | -0,8688 | 4,590E-02 | 7 | protein_coding | tyrosine hydroxylase [Source:MGI Symbol;Acc:MGI:98735] |
| Arl2 | -0,1738 | 4,591E-02 | 19 | protein_coding | ADP-ribosylation factor-like 2 [Source:MGI Symbol;Acc:MGI:1928393] |
| Gm25492 | -0,7077 | 4,597E-02 | 5 | scaRNA | predicted gene, 25492 [Source:MGI Symbol;Acc:MGI:5455269] |
| Cyp2t4 | -0,6221 | 4,597E-02 | 7 | protein_coding | cytochrome P450, family 2, subfamily t, polypeptide 4 [Source:MGI Symbol;Acc:MGI:2686296] |
| Pdcd4 | 0,1788 | 4,597E-02 | 19 | protein_coding | programmed cell death 4 [Source:MGI Symbol;Acc:MGI:107490] |
| Ms4a6c | 1,3781 | 4,599E-02 | 19 | protein_coding | membrane-spanning 4-domains, subfamily A, member 6C [Source:MGI Symbol;Acc:MGI:2385644] |
| Tmf1 | 0,2704 | 4,612E-02 | 6 | protein_coding | TATA element modulatory factor 1 [Source:MGI Symbol;Acc:MGI:2684999] |

| | | | | | |
|---|---|---|---|---|---|
| N4bp2 | 0,2835 | 4,614E-02 | 5 | protein_coding | NEDD4 binding protein 2 [Source:MGI Symbol;Acc:MGI:2684414] |
| Nolc1 | 0,1287 | 4,614E-02 | 19 | protein_coding | nucleolar and coiled-body phosphoprotein 1 [Source:MGI Symbol;Acc:MGI:1918019] |
| Cdh9 | 0,2366 | 4,617E-02 | 15 | protein_coding | cadherin 9 [Source:MGI Symbol;Acc:MGI:107433] |
| Gpbp1 | 0,1727 | 4,620E-02 | 13 | protein_coding | GC-rich promoter binding protein 1 [Source:MGI Symbol;Acc:MGI:1920524] |
| Dmc1 | 0,7740 | 4,624E-02 | 15 | protein_coding | DNA meiotic recombinase 1 [Source:MGI Symbol;Acc:MGI:105393] |
| Usp51 | 0,6896 | 4,627E-02 | X | protein_coding | ubiquitin specific protease 51 [Source:MGI Symbol;Acc:MGI:3588217] |
| Numbl | -0,1398 | 4,632E-02 | 7 | protein_coding | numb-like [Source:MGI Symbol;Acc:MGI:894702] |
| Gm44415 | -3,5720 | 4,633E-02 | 6 | TEC | predicted gene, 44415 [Source:MGI Symbol;Acc:MGI:5690807] |
| Lox | 1,1939 | 4,635E-02 | 18 | protein_coding | lysyl oxidase [Source:MGI Symbol;Acc:MGI:96817] |
| Cbx3-ps7 | -1,4029 | 4,640E-02 | 1 | processed_pseudogene | chromobox 3, pseudogene 7 [Source:MGI Symbol;Acc:MGI:3704187] |
| Rpl23a-ps14 | -3,9334 | 4,642E-02 | 11 | processed_pseudogene | ribosomal protein L23A, pseudogene 14 [Source:MGI Symbol;Acc:MGI:3649865] |
| Mitd1 | 0,3282 | 4,646E-02 | 1 | protein_coding | MIT, microtubule interacting and transport, domain containing 1 [Source:MGI Symbol;Acc:MGI:1916278] |
| Plin5 | 0,7676 | 4,649E-02 | 17 | protein_coding | perilipin 5 [Source:MGI Symbol;Acc:MGI:1914218] |
| Col4a5 | 0,4431 | 4,657E-02 | X | protein_coding | collagen, type IV, alpha 5 [Source:MGI Symbol;Acc:MGI:88456] |
| Pi4k2a | -0,1171 | 4,666E-02 | 19 | protein_coding | phosphatidylinositol 4-kinase type 2 alpha [Source:MGI Symbol;Acc:MGI:1934031] |
| Gm45425 | 2,4118 | 4,669E-02 | 8 | lincRNA | predicted gene 45425 [Source:MGI Symbol;Acc:MGI:5791261] |
| Gm15665 | -2,7207 | 4,671E-02 | 16 | processed_pseudogene | predicted gene 15665 [Source:MGI Symbol;Acc:MGI:3783107] |
| Gm20784 | 1,1904 | 4,684E-02 | 13 | processed_pseudogene | predicted gene, 20784 [Source:MGI Symbol;Acc:MGI:5434140] |
| Tmem39b | -0,2455 | 4,692E-02 | 4 | protein_coding | transmembrane protein 39b [Source:MGI Symbol;Acc:MGI:2682939] |
| Ptprh | -0,6870 | 4,701E-02 | 7 | protein_coding | protein tyrosine phosphatase, receptor type, H [Source:MGI Symbol;Acc:MGI:3026877] |
| Sh3glb2 | -0,1496 | 4,716E-02 | 2 | protein_coding | SH3-domain GRB2-like endophilin B2 [Source:MGI Symbol;Acc:MGI:2385131] |
| Tram1 | 0,1854 | 4,727E-02 | 1 | protein_coding | translocating chain-associating membrane protein 1 [Source:MGI Symbol;Acc:MGI:1919515] |
| Slc30a7 | 0,2105 | 4,729E-02 | 3 | protein_coding | solute carrier family 30 (zinc transporter), member 7 [Source:MGI Symbol;Acc:MGI:1913750] |
| Ptdss2 | -0,1506 | 4,732E-02 | 7 | protein_coding | phosphatidylserine synthase 2 [Source:MGI Symbol;Acc:MGI:1351664] |
| Gdf7 | 1,3415 | 4,740E-02 | 12 | protein_coding | growth differentiation factor 7 [Source:MGI Symbol;Acc:MGI:95690] |
| Taf12 | 0,2226 | 4,769E-02 | 4 | protein_coding | TATA-box binding protein associated factor 12 [Source:MGI Symbol;Acc:MGI:1913714] |
| C230012O17Rik | -0,6857 | 4,769E-02 | 4 | processed_transcript | RIKEN cDNA C230012O17 gene [Source:MGI Symbol;Acc:MGI:2442283] |

| | | | | | |
|---|---|---|---|---|---|
| Baalc | -0,1149 | 4,769E-02 | 15 | protein_coding | brain and acute leukemia, cytoplasmic [Source:MGI Symbol;Acc:MGI:1928704] |
| Zfp329 | 0,2150 | 4,772E-02 | 7 | protein_coding | zinc finger protein 329 [Source:MGI Symbol;Acc:MGI:1921283] |
| Wdr90 | -0,2001 | 4,775E-02 | 17 | protein_coding | WD repeat domain 90 [Source:MGI Symbol;Acc:MGI:1921267] |
| Sptb | -0,1315 | 4,776E-02 | 12 | protein_coding | spectrin beta, erythrocytic [Source:MGI Symbol;Acc:MGI:98387] |
| Slc22a12 | -0,7936 | 4,783E-02 | 19 | protein_coding | solute carrier family 22 (organic anion/cation transporter), member 12 [Source:MGI Symbol;Acc:MGI:1195269] |
| D3Ertd254e | 0,2056 | 4,788E-02 | 3 | protein_coding | DNA segment, Chr 3, ERATO Doi 254, expressed [Source:MGI Symbol;Acc:MGI:1098769] |
| CT025671,2 | -2,6875 | 4,788E-02 | 17 | processed_pseudogene | periphilin 1 (Pphln1) pseudogene |
| Gm13509 | -0,1813 | 4,788E-02 | 2 | processed_pseudogene | predicted gene 13509 [Source:MGI Symbol;Acc:MGI:3649640] |
| Dennd5a | -0,1087 | 4,790E-02 | 7 | protein_coding | DENN/MADD domain containing 5A [Source:MGI Symbol;Acc:MGI:1201681] |
| Stc2 | 0,2904 | 4,801E-02 | 11 | protein_coding | stanniocalcin 2 [Source:MGI Symbol;Acc:MGI:1316731] |
| Lims2 | -0,2339 | 4,808E-02 | 18 | protein_coding | LIM and senescent cell antigen like domains 2 [Source:MGI Symbol;Acc:MGI:2385067] |
| Evc2 | -0,2072 | 4,826E-02 | 5 | protein_coding | EvC ciliary complex subunit 2 [Source:MGI Symbol;Acc:MGI:1915775] |
| Slc2a6 | -0,2026 | 4,841E-02 | 2 | protein_coding | solute carrier family 2 (facilitated glucose transporter), member 6 [Source:MGI Symbol;Acc:MGI:2443286] |
| Gm18009 | -2,1243 | 4,843E-02 | 6 | processed_pseudogene | predicted gene, 18009 [Source:MGI Symbol;Acc:MGI:5010194] |
| Gm26830 | 2,9748 | 4,853E-02 | 11 | lincRNA | predicted gene, 26830 [Source:MGI Symbol;Acc:MGI:5477324] |
| H3f3b | 0,1105 | 4,853E-02 | 11 | protein_coding | H3 histone, family 3B [Source:MGI Symbol;Acc:MGI:1101768] |
| Enho | -0,2000 | 4,856E-02 | 4 | protein_coding | energy homeostasis associated [Source:MGI Symbol;Acc:MGI:1916888] |
| Gm4740 | 3,5528 | 4,861E-02 | 15 | processed_pseudogene | predicted gene 4740 [Source:MGI Symbol;Acc:MGI:3648965] |
| Gm16702 | 0,3763 | 4,888E-02 | 17 | lincRNA | predicted gene, 16702 [Source:MGI Symbol;Acc:MGI:4439626] |
| Myoz2 | -3,1595 | 4,901E-02 | 3 | protein_coding | myozenin 2 [Source:MGI Symbol;Acc:MGI:1913063] |
| Orc4 | 0,1845 | 4,903E-02 | 2 | protein_coding | origin recognition complex, subunit 4 [Source:MGI Symbol;Acc:MGI:1347043] |
| Phip | 0,2228 | 4,906E-02 | 9 | protein_coding | pleckstrin homology domain interacting protein [Source:MGI Symbol;Acc:MGI:1932404] |
| Gm7541 | -2,3236 | 4,929E-02 | 16 | processed_pseudogene | predicted gene 7541 [Source:MGI Symbol;Acc:MGI:3648278] |
| Rabl6 | -0,1168 | 4,931E-02 | 2 | protein_coding | RAB, member RAS oncogene family-like 6 [Source:MGI Symbol;Acc:MGI:2442633] |
| Klhdc8a | -0,3141 | 4,936E-02 | 1 | protein_coding | kelch domain containing 8A [Source:MGI Symbol;Acc:MGI:2442630] |
| Gm20468 | -2,8058 | 4,937E-02 | 17 | lincRNA | predicted gene 20468 [Source:MGI Symbol;Acc:MGI:5141933] |
| Rarb | 0,6110 | 4,942E-02 | 14 | protein_coding | retinoic acid receptor, beta [Source:MGI Symbol;Acc:MGI:97857] |

| | | | | | |
|---|---|---|---|---|---|
| Plcxd1 | -0,3042 | 4,942E-02 | 5 | protein_coding | phosphatidylinositol-specific phospholipase C, X domain containing 1 [Source:MGI Symbol;Acc:MGI:2685422] |
| Gm37498 | -2,4984 | 4,943E-02 | 3 | TEC | predicted gene, 37498 [Source:MGI Symbol;Acc:MGI:5610726] |
| Npy5r | -0,2684 | 4,952E-02 | 8 | protein_coding | neuropeptide Y receptor Y5 [Source:MGI Symbol;Acc:MGI:108082] |
| Lrrn2 | -0,1545 | 4,956E-02 | 1 | protein_coding | leucine rich repeat protein 2, neuronal [Source:MGI Symbol;Acc:MGI:106037] |
| Spryd7 | 0,1909 | 4,959E-02 | 14 | protein_coding | SPRY domain containing 7 [Source:MGI Symbol;Acc:MGI:1913924] |
| Uchl1 | -0,1599 | 4,966E-02 | 5 | protein_coding | ubiquitin carboxy-terminal hydrolase L1 [Source:MGI Symbol;Acc:MGI:103149] |
| Nxph2 | 0,7428 | 4,966E-02 | 2 | protein_coding | neurexophilin 2 [Source:MGI Symbol;Acc:MGI:107491] |
| Tas1r1 | -0,4331 | 4,970E-02 | 4 | protein_coding | taste receptor, type 1, member 1 [Source:MGI Symbol;Acc:MGI:1927505] |
| Plpp2 | -0,4839 | 4,974E-02 | 10 | protein_coding | phospholipid phosphatase 2 [Source:MGI Symbol;Acc:MGI:1354945] |
| Prpf40a | 0,2584 | 4,978E-02 | 2 | protein_coding | pre-mRNA processing factor 40A [Source:MGI Symbol;Acc:MGI:1860512] |
| Ntn4 | 0,3719 | 4,979E-02 | 10 | protein_coding | netrin 4 [Source:MGI Symbol;Acc:MGI:1888978] |
| Mus81 | -0,2003 | 4,990E-02 | 19 | protein_coding | MUS81 structure-specific endonuclease subunit [Source:MGI Symbol;Acc:MGI:1918961] |
| St3gal2 | -0,1237 | 4,998E-02 | 8 | protein_coding | ST3 beta-galactoside alpha-2,3-sialyltransferase 2 [Source:MGI Symbol;Acc:MGI:99427] |
