## Supplemental Table 2 for "The ketamine metabolite (*2R,6R*)-hydroxynorketamine rescues hippocampal mRNA translation, synaptic plasticity and memory in mouse models of Alzheimer’s disease"

| **Supplementary Table 2. Differentially expressed genes (DEGs) in APP/PS1 mice (compared to WT littermates) that were corrected by treatment with HNK.** |
|---|
| 1700025N21Rik |
| 1700099I09Rik |
| 2010001A14Rik |
| 2610012C04Rik |
| 4732496C06Rik |
| 4930461C15Rik |
| 4930505M18Rik |
| 4930550C14Rik |
| 4930579G24Rik |
| 4933407E24Rik |
| 4933428P19Rik |
| 9330175E14Rik |
| Abi3 |
| AC133505.1 |
| AC154200.1 |
| AC165079.1 |
| Acat2 |
| Acbd7 |
| Acot1 |
| Acp7 |
| Adgre4 |
| Adipor1 |
| Aga |
| Aipl1 |
| Akap12 |
| Alas2 |
| Aldh1a7 |
| Angptl1 |
| Angptl4 |
| Ankrd61 |
| Anxa2 |
| Anxa5 |
| Aox1 |
| Ap1s2 |
| Apob |
| Apold1 |
| Arhgap30 |
| Arrdc2 |
| Art5 |
| Atp6v0e |
| B4galt1 |
| Barhl1 |
| Bcl2a1a |
| Bicdl2 |

| |
|---|
| Bmf |
| Btf3 |
| C030005K06Rik |
| C130012C08Rik |
| C230035I16Rik |
| Cacng6 |
| Car13 |
| Car5b |
| Ccl9 |
| Ccnb1 |
| Ccni |
| Cd200r1 |
| Cd244a |
| Cd300a |
| Cd33 |
| Cd44 |
| Cd63 |
| Cd63-ps |
| Cd86 |
| Cdc6 |
| Cdk19os |
| Cdkn1a |
| Cenpn |
| Chil3 |
| Cldn8 |
| Clec4a1 |
| Clic1 |
| Cog2 |
| Cpt2 |
| Cript |
| Crygs |
| Cryl1 |
| Csf1 |
| CT010429.1 |
| CT025584.1 |
| CT025659.3 |
| Ctla2a |
| Cxcl1 |
| Cyp4f16 |
| Daw1 |
| Dct |
| Dhrs7b |
| Dkk2 |
| Dmrta1 |
| Dnajc12 |
| Dnase2a |

| |
|---|
| Dock2 |
| Egln3 |
| Elk3 |
| Eme2 |
| Entpd4 |
| Epha2 |
| Esyt1 |
| F2rl3 |
| Fadd |
| Fam107b |
| Fam229b |
| Fbln5 |
| Fbxo40 |
| Fcgr4 |
| Fermt3 |
| Fhad1 |
| Foxb2 |
| Frmd3 |
| Ftl1-ps1 |
| Fyb |
| Galnt15 |
| Ggcx |
| Ghrl |
| Gimap4 |
| Gm10252 |
| Gm11748 |
| Gm11772 |
| Gm12411 |
| Gm13246 |
| Gm13787 |
| Gm13986 |
| Gm14412 |
| Gm14416 |
| Gm14648 |
| Gm15238 |
| Gm17018 |
| Gm17066 |
| Gm17231 |
| Gm17344 |
| Gm17660 |
| Gm18529 |
| Gm19195 |
| Gm19385 |
| Gm1966 |
| Gm20482 |
| Gm20517 |

| |
|---|
| Gm20559 |
| Gm22260 |
| Gm22613 |
| Gm23608 |
| Gm2379 |
| Gm26594 |
| Gm26652 |
| Gm26812 |
| Gm27252 |
| Gm27651 |
| Gm28299 |
| Gm2861 |
| Gm29257 |
| Gm29361 |
| Gm29383 |
| Gm2a |
| Gm30023 |
| Gm32296 |
| Gm35248 |
| Gm35595 |
| Gm36988 |
| Gm37238 |
| Gm37403 |
| Gm37529 |
| Gm37621 |
| Gm37861 |
| Gm38244 |
| Gm38273 |
| Gm3896 |
| Gm3943 |
| Gm40309 |
| Gm41271 |
| Gm42047 |
| Gm42462 |
| Gm42510 |
| Gm42789 |
| Gm42830 |
| Gm42853 |
| Gm42970 |
| Gm43046 |
| Gm43262 |
| Gm43420 |
| Gm43435 |
| Gm43460 |
| Gm43610 |
| Gm43848 |

| |
|---|
| Gm43912 |
| Gm44090 |
| Gm44220 |
| Gm44242 |
| Gm44265 |
| Gm44815 |
| Gm44877 |
| Gm44891 |
| Gm44986 |
| Gm45124 |
| Gm45129 |
| Gm45177 |
| Gm45299 |
| Gm45493 |
| Gm45632 |
| Gm45847 |
| Gm46350 |
| Gm4707 |
| Gm47073 |
| Gm4735 |
| Gm47381 |
| Gm47590 |
| Gm47692 |
| Gm47860 |
| Gm48181 |
| Gm48632 |
| Gm48755 |
| Gm48885 |
| Gm49154 |
| Gm49226 |
| Gm4952 |
| Gm49539 |
| Gm5415 |
| Gm5855 |
| Gm6285 |
| Gm6598 |
| Gm7571 |
| Gm7656 |
| Gm7985 |
| Gm8186 |
| Gm8430 |
| Gm853 |
| Gm9402 |
| Gm9616 |
| Gm9978 |
| Gnb3 |

| |
|---|
| Gpr146 |
| Gpsm3 |
| H19 |
| Hbb-bs |
| Hbb-bt |
| Hfm1 |
| Hist1h2ae |
| Hist1h4h |
| Hist2h2ac |
| Hyal1 |
| Id2 |
| Ier3 |
| Il6 |
| Ip6k3 |
| Irf5 |
| Jagn1 |
| Kcnk1 |
| Khdrbs2 |
| Kirrel3 |
| Klhl10 |
| Klhl5 |
| Krt80 |
| Lfng |
| Lgals3 |
| Lhfpl5 |
| Lipe |
| Llgl2 |
| Loxl3 |
| Lpcat2 |
| Lpin3 |
| Lrrc9 |
| Ly6a |
| Lyl1 |
| Maff |
| Mcl1 |
| Med18 |
| Mettl13 |
| Mfap4 |
| Mfsd2a |
| Mir124a-3 |
| Mir467f |
| Mmd |
| Mrps30 |
| Ms4a4c |
| Msrb1 |
| mt-Nd3 |

| |
|---|
| mt-Ti |
| mt-Tq |
| mt-Ts2 |
| Mtmr10 |
| Naa20 |
| Ndufv2 |
| Nek3 |
| Nfkbia |
| Nipsnap3b |
| Nkapl |
| Nmur1 |
| Nod1 |
| Nr5a1 |
| Nrtn |
| Nxph3 |
| Oaz3 |
| Oscar |
| P2rx2 |
| P2ry13 |
| P2ry2 |
| P2ry6 |
| Paqr5 |
| Parp12 |
| Pcp4l1 |
| Pdk4 |
| Pex19 |
| Pfkfb4 |
| Pgm2 |
| Phox2b |
| Pirb |
| Pla1a |
| Platr17 |
| Plaur |
| Plin2 |
| Pole |
| Popdc3 |
| Prcp |
| Prdx4 |
| Prkag2os1 |
| Prkch |
| Prkd2 |
| Psat1 |
| Rcsd1 |
| Renbp |
| Rhoa |
| Rhoc |

| |
|---|
| Rhoq |
| Rin3 |
| Ripply3 |
| Rpl14 |
| Rpl14-ps1 |
| Rpl17 |
| Rpl17-ps3 |
| Rpl19-ps11 |
| Rpl23a-ps3 |
| Rpl26 |
| Rpl34-ps1 |
| Rpl36a |
| Rpl7a-ps3 |
| Rplp0 |
| Rps11-ps1 |
| Rps6 |
| Rps6ka1 |
| Rpsa |
| Rsph3a |
| Sar1b |
| Sash3 |
| Scpep1 |
| Selenof |
| Selp |
| Serpina3g |
| Serpinb1b |
| Sgk1 |
| Sgk3 |
| Sh2d6 |
| Slc12a7 |
| Slc1a5 |
| Slc25a13 |
| Slc25a45 |
| Slc2a1 |
| Slc46a3 |
| Slfn2 |
| Smarca5-ps |
| Smyd4 |
| Snord17 |
| Snrpe |
| Soat2 |
| Sord |
| Sparc |
| Spi1 |
| Srd5a2 |
| Srgn |

| |
|---|
| Stat5a |
| Stk17b |
| Sult1a1 |
| Sult6b1 |
| Sumf2 |
| Svopl |
| Tagln2 |
| Tapbp |
| Tbxas1 |
| Tlr7 |
| Tmem254c |
| Tmem37 |
| Tmem9 |
| Tnfrsf10b |
| Tnk2os |
| Tph1 |
| Tpi-rs11 |
| Tram1 |
| Treml2 |
| Trim16 |
| Trip10 |
| Trpm8 |
| Tsacc |
| Tspo |
| Tuft1 |
| Txnip |
| Ush1c |
| Vmn2r56 |
| Wdr38 |
| Wrb |
| Zbtb7b |
| Zfp36 |
| Zfp58 |
| Zfp954 |
| Zfp968 |
| Zfp981 |
